# Tensions in tillage: Reduction in tillage intensity associates with lower wheat growth and nutritional grain quality despite enhanced soil biological indicators

**DOI:** 10.1101/2024.12.09.627503

**Authors:** Matthias Waibel, Jennifer Michel, Maurine Antoine, Iñaki Balanzategui-Guijarro, Da Cao, Pierre Delaplace, Jacques Le Gouis, David Alvarez, Claire Léon, Sandy Manfroy, Jordi Moya-Laraño, Sibille Perrochon, Sara Sanchez-Moreno, Inés Santin-Montanya, José Luis Tenorio, Cécile Thonar, Hervé Vanderschuren, Dominique Van Der Straeten, Thomas Verlinde, Markus Weinmann, Sarah Symanczik

**Author notes:** Corresponding author: Sarah Symanczik.

## Abstract

Dryland ecosystems are particularly susceptible to the adverse effects of intensive agriculture, with intensive tillage exerting a major impact on soil health and its biotic components. The implementation of less disturbing soil management practices can be essential for preserving the soil environment and maintaining the diverse communities of microorganisms, micro- and mesofauna, which are essential contributors to soil fertility. In this study, we assessed soil chemical properties, soil biodiversity and functionality, and wheat crop growth across a tillage gradient encompassing no-tillage (NT), minimum tillage (MT), and standard tillage (ST). Results showed that NT resulted in increased soil macronutrient levels compared to MT and ST. In general, reduced tillage increased the abundance of soil biota, with significantly higher levels of bacterial and fungal marker genes observed in MT and NT compared to ST. Nematode abundance increased by 25% in MT and 50% in NT, compared to ST and predatory acari were significantly more abundant in NT, while numbers of total acari were higher in both NT and ST compared to MT. Community structure analysis revealed that tillage strongly influenced bacterial, fungal and acari community composition, reflecting a gradient of soil disturbance intensity. Corresponding to the increased abundance of soil biota, reduced tillage increased microbial activity and soil functionality along the disturbance gradient. This was evident in the potential activity of carbon, nitrogen and phosphorus cycling enzymes, as well as the microbial capacity for carbon utilisation. In addition, evidence of the formation of biocrust as a possible source of carbon input was found. Furthermore, we observed important wheat pathogens to decrease and fungal antagonists to increase in NT compared to ST. Despite enhanced soil biological indicators under reduced tillage, wheat growth, nitrogen uptake and grain B vitamin contents were higher in ST compared to NT. In addition, we observed a shift in technological grain properties across tillage practices. The higher root:shoot ratio (an indicator of nitrogen deficiency) and median root diameter (hormone-driven lateral expansion) in NT suggest that soil compaction could be a potential cause of reduced wheat performance. These results suggest that despite improved soil biological indicators, other factors such as a low rates of N mineralization potential and prevalence of soil compaction may be limiting wheat performance in NT systems.

**Highlights:**

- Enhanced microbial activity and functionality under reduced tillage
- Tillage intensity shaped community structure of microbes, nematodes and acari
- Soil biocrust development under NT may increase soil organic carbon
- Root traits revealed soil compaction and nutrient limitation in NT systems
- Reduced tillage impaired wheat quality and changed technological grain properties

## 1. Introduction

Dryland agro-ecosystems are particularly vulnerable to the detrimental effects of intensive tillage, which causes upturn and disruption of soil structure, reduction of soil organic matter (SOM) content, and ultimately altering soil biodiversity and functioning (Degrune et al., 2016; Haddaway et al., 2017; Kraut-Cohen et al., 2020; Li et al., 2019; Peng et al., 2020). As studies have shown, reducing tillage intensity has the potential to increase topsoil SOM, reduce soil erosion, enhance water retention, and thus it can help to preserve the integrity of soils (Haddaway et al., 2017; Holland, 2004; Lal, 2009).

Soil organisms provide multiple agroecosystem services including nutrient cycling, mitigation of biotic and abiotic stresses and thus soil and plant health (Bender et al., 2023; Delgado-Baquerizo et al., 2016, Delgado-Baquerizo et al. 2020; Wagg et al. 2019, Wagg et al., 2021). Several studies have documented the beneficial impacts of reduced tillage on increasing the abundance of soil microorganisms (Chen et al., 2020; Essel et al., 2019; Helgason et al., 2009; Mathew et al., 2012; Morugán-Coronado et al., 2022) and micro- and mesofauna (Brévault et al., 2007, Postma-Blaauw et al. 2010, Betancur-Corredor et al. 2022).

Through the disruption of fungal mycelia, intensive tillage is suspected to negatively impact fungal communities (Roger-Estrade et al. 2010), while higher soil organic carbon (SOC) under reduced tillage practices should enhance microbial abundance and activity (Lori et al. 2017, Ramírez et al., 2020). However, a recent meta-analysis found that the response of soil microbial diversity to tillage is highly variable and strongly depends on pedoclimate and plant growth stage (Li et al., 2024, de Graaff et al., 2019): While minimum tillage increased bacterial diversity by 7%, there was no significant effect on fungal diversity compared to ploughed systems. Similarly, Capelle et al. (2012) have shown that the effect of tillage intensity on the abundance of soil biota might differ depending on soil texture. Given the central role of local pedoclimatic conditions in modulating the impact of tillage practices on soil microbial communities, it is essential to approach each environmental context individually to gain a full understanding of their responses. This involves to further identify key organisms responsible in shaping the soil environment, including those within the soil micro- and mesofauna, which are often neglected.

So far, most field studies investigating belowground effects of tillage practices have focussed either on the abundance and/or community structure of soil biota or on soil functional properties but without a link to aboveground performance (Capelle et al. 2012, Roger-Estrade et al. 2010). Furthermore, there is still limited understanding about the driving factors behind crop performance under different tillage practices, as literature presents contradictory results: the effects of no-tillage and minimum tillage reported in meta-analyses are ranging from negative to neutral effects (Young et al. 2021). The meta-analysis of Pittelkow et al. (2015) demonstrated best performance of no-tillage under rainfed conditions in dry climates, matching conventional tillage yields, while average yield of 50 crops reduced by 5.1%. Thus, further studies are needed that link soil biodiversity and functioning with crop growth and grain quality.

In the present study we investigated how tillage intensity in a long-term field experiment affects soil properties and the interplay between soil biodiversity, functionality and crop performance. Winter wheat was grown in crop rotation comparing no-tillage (NT), minimum/reduced tillage (MT) and standard inversion tillage (ST). We assessed soil chemical, biological and functional properties to identify tillage practices enhancing belowground functioning. Soil chemical factors comprised a range of soil macro and micronutrients and soil pH. Soil biological variables included the abundance, diversity and community structure of soil micro- and macrobiota including bacteria, fungi, nematodes and acari. Soil functional properties focused on the C-, N- and P-cycling potential of the soil, substrate utilization potential and microbial activity in general. Further, in an effort to establish connections to various agronomic qualitity indicators of winter wheat, we characterised shoot and root traits, grain nutrients and B vitamins as indicators for nutritional quality as well as starch size distribution and gliadin and glutenin contents in grains as technological quality indicators.

We hypothesised that: i) tillage intensity would be negatively correlated to SOC, soil biodiversity and functionality, ii) tillage intensity would drive shifts in the community structure of soil biota, namely bacteria, fungi, nematodes and acari, and iii) that an increase in SOC and higher nutrient cycling potential could translate into an enhanced uptake of nutrients leading to higher wheat biomass and grain nutrient and B vitamin content.

## 2. Material and Methods

### 2.1 Site and soil characteristics

The experimental field for this investigation was the “La Canaleja” long-term tillage comparison trial of the Spanish National Research Institute for Food Research and Technology (INIA-CSIC), established in 1994 outside Alcalá de Henares (Madrid, Spain, 40°30’55.0" N 3°18’37.1" W; 600 m a.s.l.). The soil texture is classed as a Loam (USDA classification) and the soil type as a calcic Haploxeralf. The climate is semi-arid Mediterranean with the majority of the average annual precipitation of 367 mm yr^−1^ [1994-2022] having occurred in the autumn, winter, and spring months. In recent years, the average monthly temperature in summer has increased by ca. 1.3 °C and rainfall amounts have decreased in the months October to May by average total of 67 mm (period 2017-2022 vs. 1994-2022, own climatic data recorded on-site). The fallow period in the year before and the full winter wheat growing season 2021-2022 in the vegetative period (October to sample taking at flowering stage in May) did not exhibit a large total precipitation deficit (+99 mm and -13 mm, for the periods respectively) as compared to the averages of the previous years (2017-2022). However, some months in the period January to May of 2022 were much drier as compared against the average 2017-2022 (total 14 mm vs. 39 mm). Indeed, the soil moisture in this experiment was extremely low at ca. 2.7%, where another study by Santin-Montanya et al., (2020) reported around 10% at wheat flowering stage for the years 2014 to 2016.

### 2.2 Experimental design

The three field tillage treatments consist of standard/conventional tillage (ST), minimum/reduced tillage (MT), and no tillage (NT) and are arranged in a randomized complete block design (4 blocks = treatment replicates) with 5 split-plots within each block of which four contain crop rotations (Fallow, winter wheat, vetch, barley) and the fifth containing a wheat monoculture. In the crop rotation before winter wheat, one year of bare fallow is employed to accumulate rainwater for subsequent wheat cropping. The tillage practices for ST consist of mouldboard ploughing (30 cm), chisel (non-inversion) ploughing (20/15 cm) for MT, and no tillage for NT with herbicide treatments when needed for weed control middle of May and always at the beginning of October before sowing. Further details on seeding, nutrient, pest, weed, and residue management, as well as crop performance, soil physico-chemical, and climate aspects on this long-term trial were reported in preceding studies (Gandía et al., 2021, Santin-Montanya et al., 2020, 2017, 2016, 2013, Tellez-Rio et al., 2017, Guardia et al., 2016, Martin-Lammerding et al., 2015, 2013, 2011, Martin-Rueda, et al., 2007).

### 2.3 Field sampling and sample processing

Shoots, roots and soil from the split-plots of winter wheat (*Triticum aestivum* var. Marius) in crop rotation were sampled at flowering at the end of May 2022. Rhizosphere soil for the analysis of soil microbial communities (bacteria, fungi), microfauna (nematodes), and mesofauna (acari) was sampled as composite sample of five samples per plot. First, plants were cut at the crown and then a shovel was used to extract the root system and adjacent soil from an area diameter of 15 cm at two depths 0-10 cm and 10-20 cm. Nematodes were sampled from a combined depth of 0-20 cm only. The soil at the center of the extracted root system was shaken off and transferred to plastic bags for subsequent DNA and soil functional analyses. For mesofauna, samples were carefully taken at the edges of the sampling holes as intact pieces of soil blocks with a total volume of 250 ml per plot and kept in sturdy plastic containers to avoid structural disturbance. For nematodes, a combined sample of 200 g soil per plot was taken and kept in sealed plastic bags. In between wheat rows, 10-20 additional soil samples were taken from 0-20 cm depth and combined to obtain 2 kg per plot for soil chemical analyses. For analysis of root system architecture, two additional samples were taken per plot: One root sample was taken with a precise volume (h=20 cm, d=9 cm) using a pneumatic soil corer (Royal Eijkelkamp B.V, The Netherlands) to determine root length and other quantitative parameters of root system architecture within that soil cylinder and an additional sample was taken from the top 10 cm extracting the root crown to count first to third order lateral roots and measure the respective root angles. For each root sample, corresponding aboveground plant biomass was sampled and kept in paper bags for drying at room temperature. All soil and root samples were immediately cooled to 4°C and transported to the lab. For further analysis, soil samples were sieved at 2 mm and stored at 4°C for MicroResp® analyses or frozen at -20°C for DNA and enzymatic anaylses.

Root samples were carefully washed with water over 1 mm sieves to separate roots from soil. Clean roots were subsequently stored in 70% ethanol. For quantitative root traits such as total root length and network area, the washed root samples were transferred to petri dishes and arranged without overlap of roots. Samples were scanned (Epson flatbed, 600 DPI, Seiko Epson Corporation, Japan) and images were subsequently processed in RhizoVision Explorer (Seethepalli et al., 2021). The root crown samples were digitized using Nikon D3400 (Nikon, Tokyo, Japan) and images were processed in RhizoVision Explorer (Seethepalli et al., 2021) and phenotyped for root angles and lateral root growth (Trachsel et al. 2011). Afterwards, root samples were dried at 40°C in paper envelopes and root dry mass was determined. Corresponding aboveground biomass was separated into leaves, shoots and heads and dry weight determined.

### 2.4 Soil chemical analysis

Soil gravimetric moisture content was determined by drying over night at 105 °C. Soil C and N (%) were determined on a Vario Max Cube (Elementar Analysensysteme GmbH, Langenselbold, Germany). Further soil elements were determined on inductively coupled plasma optical emission spectroscopy (ICP-OES, Agilent 5110, Agilent Technologies Inc, USA) with extractants as detailed in the following. CaCl2 extraction matrix was used for pH determination and estimation of plant available Mg. Calcium Acetate Lactate (CAL) extraction matrix was used for reflecting plant available K and P (VDLUFA 2016, Scherer and Weichmann (1991). Calcium chloride/DTPA (CAT) extraction matrix was used for estimating plant available Cu, Fe, Mn, P, and Zn (VDLUFA 2008). Nitric acid and hydrochloric acid (Aqua regia (AR)/ King’s water) digestion was applied to determine total B, Ca, Cu, Fe, K, Mg, Mn, P, and Zn concentrations in soil (VDLUFA 1991). B concentrations were below the detection limit of 10 mg kg^-1^.

### 2.5 Soil micro- and mesofauna analyses

Acari and Collembola were extracted from 500 ml of soil using Berlese-Tullgren funnels with LED lamps (non incandescent 7 W bulbs of 600 lumens) for 15 days at constant temperature (23 °C) and stored in 70% ethanol. To identify the collected specimens to family level, dichotomous keys were used, e.g. Krantz & Walter (2009) for Acari and Jordana & Arbea (1989) for Collembola. The specimens were preserved in 70% ethanol or mounted in semi-permanent slides with Hoyer’s medium, and have been deposited in the "Mesofauna collection" at the Arid Zones Experimental Station (EEZA-CSIC). Only 4 specimens of Collembola were collected in total and we will therefore omit them from further discussion. To assign mite families to trophic guild (predator, fungivore, herbivore) the information in Krantz & Walter (2009), in addition to expert knowledge (e.g. shape of the mouth parts), were considered.

Nematodes were extracted by a modification of wet sieving and the Baermann funnel method (Barker, 1984). All nematodes were counted under a dissecting microscope and at least 100 individuals were identified to genus level under a light microscope after Bongers (1990). Nematode abundances were expressed as number of individuals 100 g fresh soil^-1^. Nematode genera were classified into five trophic groups (bacterivores, fungivores, herbivores, omnivores, and predators) after Yeates (1994), and into the colonizer-persister (cp) scale, that classifies nematodes into five cp groups (cp1-5). Nematodes in low cp groups (cp 1-2) are commonly r-strategist bacterivores and fungivores with high reproduction rates, short life cycles, and resistance to environmental perturbation, while high cp groups (cp 3-5) are K-strategists with progressively longer life cycles, lower reproduction rates and higher sensitivity to perturbation and environmental stress (Bongers, 1990). Based on such functional classifications, nematode participation on nutrient cycling was assessed through the use of three nematode-based indices (Ferris, et al., 2001; Ferris, 2010). The Enrichment Index (EI) is based on the relative abundances of bacterivores with low cp value which are enrichment- opportunists, and is used as an indicator of fast organic matter decomposition mediated by bacteria and active nutrient cycling. The Channel Index (CI) is based on the ratio fungivores to opportunistic bacterivores and is used as an indicator of slow organic matter decomposition mediated by fungi. Both the EI and the CI range from 0 (no enrichment opportunists (EI) or fungal feeders (CI) present in the community) to 100 (absolute dominance of such functional groups). Finally, the Enrichment Footprint (EF) is a calculation of the potential amount of C mineralized (µg C kg^-1^ soil) by enrichment opportunistic nematodes and thus an indicator of the magnitude of the participation of such nematodes in the nutrient mineralization soil service.

### 2.6 Molecular microbial analysis

#### 2.6.1 DNA extraction

DNA was extracted from 350 mg fresh weight soil using the Macherey Nagel NucleoSpin 96 Genomic DNA soil kit with the enhancer solution following the manufacturers recommendations. Prior to extraction, samples were homogenized twice for 5 min in a TissueLyser (Qiagen, Hilden, Germany) at 30 Hz in two different orientations and stored at - 20 °C. Extract quality was checked with Nanodrop 260/230 and 260/280 nm absorption and on 1.5% agarose gel electrophoresis. DNA concentration was quantified with Qubit dsDNA HS Assay Kit on 384 well plate reader (Tecan 200 Pro M Nano+).

#### 2.6.2 Quantitative PCR

DNA extracts were diluted 10-times for quantitative PCR in a CFX384 Real-Time System (Bio-Rad Laboratories, CA, USA). Prokaryotic 16S marker gene quantification was performed in triplicate PCR reactions of 15 μL consisted of 7.5 μL 1x KAPA SYBR Fast qPCR Kit Master Mix 2× Universal (Axonlab, Baden, Switzerland), 1.8 μL of each 10 μM forward and reverse primer (BactQuant, Liu et al. 2012) and 2 μL of DNA template. Reaction conditions consisted of an initial denaturation at 95 °C for 3 min, and 39 cycles of denaturation at 95 °C for 15 s, annealing at 62 °C at 15 s, with final extension at 72 °C for 30 s. Melt curve analysis was performed from 65-98 °C. The amplification efficiency was between 68.9%-70.6% and R^2^ 99.7- 99.8%. Prokaryotic copies were quantified against a ten-fold dilution series in triplicate of a 465 bp long target region on pJET 2974 plasmid (CloneJET PCR Cloning Kit, Thermo Fisher Scientific, Switzerland). Eukaryotic/fungal 18S marker gene quantification was performed in triplicate PCR reactions of 15 μL consisted of 7.5 μL 1x KAPA SYBR Fast qPCR Kit Master Mix 2× Universal (Axonlab, Baden, Switzerland), 0.75 μL of each 10 μM forward and reverse primer (FR1 and FF390, VAINIO and HANTULA, 2000) and 2 μL of DNA template. Reaction conditions differed to the bacterial qPCR in that 35 cycles were carried out with annealing at 50°C. The amplification efficiency was between 92.8%-95.9% and R^2^ 99.6-99.9%. Eukaryotic copies were quantified against a ten-fold dilution series in triplicate of a 390 bp long target region on pJET 2974 plasmid (CloneJET PCR Cloning Kit, Thermo Fisher Scientific, Switzerland).

#### 2.6.3 Metabarcode analysis of bacterial and fungal community

Bacterial and fungal amplicon libraries were performed in a limited cycle PCR amplification approach followed by an Illumina barcode indexing PCR for sample pooling (Illumina 16S Metagenomic Sequencing Library Preparation, Genetic Diversity Center ETH Zürich). For amplification of the 16S V3-V4 target region, primers 341F (5’- CCTAYGGGDBGCWSCAG- 3’) and 806R (5’-GGACTACNVGGGTHTCTAAT-3’) (Frey et al., 2016) were used with added nextera and sequencing adaptors. The fungal ITS2 target region enclosed by the 5.8S and 28S gene cassette was amplified with ITS3ngs (5’-CANCGATGAAGAACGYRG-3’) and ITS4ngs (5’- CCTSCSCTTANTDATATGC-3’) (Tedersoo and Lindahl, 2016) and the same adaptors as used for bacteria. The first PCR was performed in triplicate PCR reactions of 15 μL consistent of 7.5 μL 1x KAPA SYBR Fast qPCR Kit Master Mix 2× Universal (Axonlab, Baden, Switzerland), 0.4 μL each of 10 μM forward and reverse primers, and 2 uL of DNA template of different DNA template dilutions (1:5, 1:10, 1:25 for bacteria and 2 x original concentration and 1:5). The reaction conditions for bacterial and fungal sets consisted of initial denaturation at 95 °C for 3 minutes, followed by 35 cycles (bacteria) or 40 cycles (fungi) of denaturing 95 °C for 20 s, annealing 58 °C (bacteria) and 60 °C (fungi) for 15 s and extension 72 °C for 40 s and a final extension at 72 °C for 10 min. Triplicate reactions were pooled and purified using a magnetic bead solution (https://openwetware.org/wiki/SPRI_bead_mix) and visualized on agarose gel (1.5%). The indexing PCR step was carried out as per default instructions of the Ilumina indexing kit where the template concentration was prediluted to 0.1-0.5 ng DNA μL-1. Index PCR reactions were purified using a magnetic bead solution (https://openwetware.org/wiki/SPRI_bead_mix), visualized on agarose gel (1.5%) and quantified with Qubit dsDNA HS Assay Kit on a 384 well plate reader (Tecan 200 Pro M Nano+) before library pooling. Final libraries were quantified and size checked with TapeStation (4150 TapeStation System, Agilent Technologies, Switzerland). Both libraries including non-template PCR/library control (n=1) and processing blanks (n=2) were sequenced separately on Illumina MiSeq v3 (Illumina Inc., San Diego, CA, USA) paired end 2x300 bp sequencing at Genome Quebec CES (Montreal, Canada) with ∼7% PhiX per sequencing run.

Sequencing reads were prepared at Genetic Diversity Center (GDC), ETH Zürich, in the USEARCH (Edgar, 2010) pipeline. Reads were demultiplexed, PhiX and low complexity samples were removed, and reads trimmed at each end with 30 or 55 bp length. Read pairs were synchronized, merged with minimum 30 bp overlap, 100 bp length, and 60% identity, and primer sites removed in-silicio. Size and quality of amplicons was selected at 300-500 bp length for 16S and between 200-500 bp for ITS2 with minimum mean Q score of 20 leaving 6.4 mil. (5.5 mil. unique) and 11.2 mil. (1.7 mil. unique) amplicons for 16S and ITS2 sets, respectively. UNOISE3 was used to deduplicate amplicons and denoise sequences with zero radius operational taxonomic units clustered de-novo at 97% similarity (97% zOTUs). Amplicons were backmapped to OTUs (counting). Taxonomy assignment was made with SINTAX for 16S against SILVA SSU v138 (RESCRIPt) (Quast et al., 2012) with tax filter at 0.75. For ITS2, taxonomy assignment was made against UNITE Eukaryotes USEARCH/UTAX release v83 (Nilsson et al., 2019) with tax filter at 0.75. Additionally, ITSx (Johan Bengtsson-Palme et al., University of Gothenburg Version: 1.1.3) was used to extract and identify the ITS region. Reads that were not classified as fungi based on ITSx were removed prior to further cleaning detailed in the next section. Cleaning of non-target taxa and contaminants was done on the whole set before splitting into the field experiments. For the 16S data, archaea, mitochondria and chloroplasts were removed. All taxa (n=13) found in NTC and blanks were removed from analysis. The average bacterial read count was 18,355 reads, minimum reads at 7,793, consisting of 1,666 taxa, and with a data sparsity of 35%. For the ITS2 dataset, all phyla other than fungi and taxa found in NTC and blanks (n=214) were removed. On average 14,012 reads remained with a minimum at 7,042 reads of 1,277 taxa with a data sparsity of 75%.

### 2.7 Soil functionality

Activities of seven hydrolytic extracellular enzymes were measured, including four enzymes involved in C-acquisition: α-glucosidase, β-glucosidase, cellobiohydrolase, β-xylanase; two enzymes involved in N-acquisition: leucine aminopeptidase, β-N-acetylglucosaminidase, and one enzyme involved in P-acquisition: phosphatase. Extracellular enzyme activity (EEA) was assessed using standard fluorometric techniques and the activities, represented by nmol g^−1^ dry soil h^−1^, of the C, N, and P acquiring enzymes were grouped to represent the general potential C, N, and P acquisition activity in soil samples (Bell et al., 2013).

Chemotrophic, respiratory soil CO2 evolution upon addition of different substrates was determined with Microresp^TM^ system (Campbell et al., 2003). Absolute respiration rates (μg CO2–C g^−1^ dry soil eq. h^−1^) were calculated as shown in the MicroResp™ technical manual (Cameron, 2007). For each sample, eight technical replicates were measured and obvious outliers removed (coefficient of variance > 20). In addition, reference soils were included as standards in each run (Creamer et al., 2016). Soils were pre-incubated for one week at the later assay temperature and soil basal respiration was measured as soil samples supplemented with deionized water, and multiple substrate-induced respiration (MSIR) as the total CO2 flux from all substrates (keto-glutaric acid, oxalic acid, xylan, glucose, N-acetyl- glucoseamine, alanine, amino-butyric acid). Microbial biomass (SIR-MBC) was estimated from the glucose (ca. 59 mg g^-1^ soil dry weight eq.) induced average respiration rate volume after 5 h of incubation and using the formula from Anderson and Domsch (1978). The metabolic quotient (qCO2) was calculated as basal respiration over SIR-MBC (Wardle and Ghani, 1995, Anderson and Domsch, 2010).

### 2.8 Wheat harvest analysis

At maturity, grains were harvested using a plot harvester. Grain yield was assessed after cleaning and air drying the grains. Thousand-kernel weight was measured with an Opto-Agri (Optomachine, Riom, France). For mineral nutrient analyses in wheat grains, dried and milled samples were mineralized by microwave (UltraClave V Fa. MLS Leutkirch, Germany) digestion with HNO3. (VDLUFA, 2021). In samples where silicon was to be analyzed, a second microwave digestion was performed and subsequently 0.5 ml HF were added to inhibit polymerization and precipitation of the dissolved silica. Grain nutrient analysis of C, N, S was determined via Elemental-Analyzer (Elementar Vario EL, Elementar Analysensysteme GmbH, Langenselbold, Germany; VDLUFA, 2012), Cl on Ion Chromatography (Integrion, Thermo Fisher Scientific, Waltham, MA, USA; VDLUFA, 2019), Ca, Cu, Fe, K, Mg, Mn, Na, P and Si on inductively coupled plasma optical emission spectroscopy (ICP-OES, Agilent 5110, Agilent Technologies Inc, USA), Co, Mo, Ni, and Se on ICP-MS (Perkin Elmer NexION 300 X, PerkinElmer, Waltham, MA, USA), and I with ammoniacal extraction on ICP-MS (Perkin Elmer NexION 300 X, PerkinElmer, Waltham, MA, USA). All analyses were conducted at Core Facility University of Hohenheim, Analytical Chemistry Unit. All Na and I was below the detection limit at 20 mg kg-^1^ or 0.5 mg kg^-1^, respectively. Most values of Se were below detection limit at 0.025 mg kg^-1^. For Si and Co some values were below detection limit and were replaced with half detection limit value at 150 mg kg^-1^and 0.025 mg kg^-1^, respectively.

Analysis of seven grain B vitamins was performed as reported previously (Cao et al., 2024). Briefly, around 50 mg of homogenized wheat seeds were extracted with 1 mL of 50 mM phosphate buffer containing internal standards for each type of B vitamin. The extract was treated with multi-enzyme incubation to release protein-bound forms and associated bioactive derivatives and analyzed with liquid chromatography coupled with tandem mass spectrometry (UHPLC-MS/MS).

Technological properties of wheat grains included the distribution of starch size classes distinguishing between A-type (>10 µm), B-type (2-10 µm) and C-type (<2 µm) starch and grain glutenins and gliadins contents. The size distribution of the purified starch granules was estimated with a Mastersizer 2000E laser particle size analyzer and the Hydro 2000SM small volume wet sample dispersion unit (Malvern Panalytical Ltd, Palaiseau, France) following the protocol reported by Edwards et al (2008). Protein composition was determined after sequential extraction. Briefly, 66.6 mg fresh weight of complete flour were mixed with 1 mL of 50 mM phosphate buffer (pH 7.8) containing 100 mM NaCl (30 min, 800 rpm, 4 °C). After centrifugation (10 min, 18,000 g, 4 °C), the pellet was mixed with 1mL of the same 50 mM phosphate buffer (10 min, 800 rpm, 4 °C) and centrifuged in the same condition. This operation was repeated once. To extract gliadins the pellet was mixed with 1 mL 70% ethanol (30 min, 1100 rpm, 4 °C). After centrifugation (10 min, 18,000 g, 4 °C), extraction was done with the same 70% ethanol buffer (10 min, 1100 rpm, 4 °C) and centrifuged. This operation was repeated twice. After each centrifugation the supernatant containing gliadins was collected and pooled. Finally, to extract glutenins the resulting pellet was mixed with 0.5 mL of 25 mM borate buffer (pH 9.8) containing 50 % propanol-2 and 1% DTT (30 min, 1200 rpm, 50 °C) and centrifuged (10 min, 18,000 g, 18 °C). This operation was repeated once. After each centrifugation the supernatant containing glutenins was collected and pooled. For stability, glutenin fraction was alkylated with 4.6% of 4-Vinylpyrridine (15 min, 60 °C). The two glutenin subunits (High-Molecular Weight Glutenin Subunit, HMW-GS; Low-Molecular Weight Glutenin Subunit, LMW-GS) and four gliadin classes (α/β-, γ-, ω1,2-, ω5-gliadin) were quantified by reverse phase high performance liquid chromatography (RP-HPLC) using an Agilent 1290 Infinity LC system (Agilent Technologies, California, USA) as described by Dai et al. (2015). Briefly, gliadin and glutenin extracts were filtered through regenerated cellulose syringe filters (0.45-µm pore diameter, UptiDisc, Interchim, Montluçon, France), then 4 µl of each protein extract were injected into a C8 reversed-phase ZORBAX 300 Stable Bound column (2.1 × 100 mm, 3.5 µm, 300 Å; Agilent Technologies) maintained at 50°C. Proteins were separated at a flow rate of 1 ml/min using linear solvent gradients from 24 to 50% acetonitrile containing 0.1% (v/v) trifluoroacetic acid over 13 min for gliadins, and from 23 to 42% over 25 min for glutenins. Proteins were detected by UV absorbance at 214 nm. Chromatograms were processed with ChemStation 10.1 software (Agilent Technologies) and the HPLC peaks corresponding to each of the four gliadin classes and the two glutenin subunits were identified following the observations of Wieser et al. (1998; Figure S4). All quantities were corrected to take into account the extraction yield which was estimated to be 93% for all fractions. In addition, the glutenin quantities were then corrected to take into account the extraction yield without SDS which was estimated to be 70% (Nicolas et al 1998). Protein quantities were determined with a calibration curve based on the Dumas combustion method using a FlashSmart N Analyzer (ThermoScientific, Villebon-sur-Yvette, France).

### 2.9 Statistical analyses

For data handling, descriptive analysis, and plotting *R* version 4.3 (R Core Team, 2023) was used with tidyverse, grid and gridExtra packages (Wickham et al. 2019, RCore team 2023, Auguie, 2017). Hulls were drawn with ggConvexhull (Martin, 2017) and labels with ggRepel (Slowikowski, 2023). Treatment effects on univariate data (soil chemical, wheat traits, organism abundance, Shannon diversity, and functional activity) were tested by ANOVA with linear model of block and management as fixed factors when from one/composite depth. The models for ratio of eukaryotic to prokaryotic qPCR values and root to shoot ratio were tested as ANCOVA with nominator as response and denominator as covariate. Values were Box-Cox transformed when multiple DHARMa diagnostics (Hartig, 2022) indicated violations. Models that still failed in DHARMa diagnostics are reported in brackets. Fixed factors were tested with type III ANOVA. With data from two depths, (Generalised) Linear Mixed effect Models (GLMM, GLM, or LMM) were used as appropriate for type of response variable with block, management, depth, and management:depth interaction as fixed factors and plot as random factor. The metabolic quotient as the ratio of basal respiration over biomass was modelled as LMM in glm function with denominator as B-spline in splines package (R core team, 2023). For acari count data, GLMM with negative binomial family was used after comparing Akaikès information criterion with Poisson family model. For LMM models, fixed factors were tested with type III ANOVA, Kenward-Roger F-test and Satterthwaite’s degrees of freedom. For GLM and GLMM models, fixed factors were tested with type II ANOVA and Likelihood Ratio test or Wald Chi square test, respectively. For all models, values reported are estimated marginal means from linear combination of model coefficients and were back-transformed for reporting. Brackets report the 95% confidence interval (CI). R packages used were base stats, spline, lme4, glmmTMB, lmtest, DHARMa, emmeans, car & effects, and multcomp (RCore team 2023, Bates et al., 2009, Brooks et al., 2017, Zeileis and Hothorn, 2002, Hartig, 2022, Lenth, 2023, Fox and Weisberg, 2019, Hothorn et al., 2008).

For amplicon sequencing data, rarefaction was made to the minimum sequencing depth sample (Schloss, 2024) with vegan’s rrarefy (Oksanen et al., 2022). Rarefaction with 999 repeats was used for repeatedly calculating alpha and beta diversity indices for averaging over sub-sampling results (Cameron et al., 2021). Beta diversity (site dissimilarity) for bacteria, fungi, acari and nematode data was calculated with Bray-Curtis index (Bray and Curtis, 1957). Alternatives such as e.g. centered-log ratio transforms and Aitchison’s distance to address compositionality (Gloor et al., 2017, Zhou et al., 2022), were also checked (not shown), but were argued to be unnecessary to control type I error of community-wide tests (Zhou et al., 2022). Homogeneity of multivariate group dispersions were tested with betadisper test (Anderson, 2006) in vegan at 5% significance level. Clustering with unweighted pair-group method arithmetic average (UPGMA) and ordination of sample to sample dissimilarities were done with Principal Coordinate Analysis (PCoA) in base R cmdscale function, respectively. Significance of tillage and depth factors (incl. interactions) on community dissimilarity were determined with the permutational multivariate ANOVA (perMANOVA) (Anderson, 2001) in vegan with 9999 permutations and restricting permutations to block when block was found to be significant at p <0.05.

Differential abundance of bacterial and fungal taxa was determined with LinDA (Zhou et al., 2022) at 5% FDR on the rarefied dataset and using centered log-ratio transform (Yang and Chen, 2023, 2022). Combinations of depth and tillage, and block were considered as fixed factors and plot as random factor. Contrasts with significantly differential abundant taxa were retrieved from model coefficients in LinDA Wald test with multiple test correction by the Benjamini-Hochberg method. In this dataset, especially for the sparse fungal set, the LinDA method reported conservative positive detections among other tools employed, such as for example DESeq2 (Love et al., 2014) or per taxon application of glmmTMB (Brooks et al., 2017) (not shown). For cluster analysis, significant taxa were gathered and their log2 transformed counts (+1 pseudo-count) subtracted from their global mean, Euclidean distances calculated, and hierarchically clustered with Ward’s rule (Love et al., 2014). Heatmaps were drawn in pheatmap package (Kolde, 2019) and the number of informative “change” clusters were identified with Gap statistic (b=999 folds) and the Tibshirani et al. (2001) criterium in cluster package (Maechler et al., 2022, Akalin, 2020). The silhouette (similarity) value (Rousseeuw, 1987) of clusters and taxa to their peers was calculated (Akalin, 2020).

## 3. Results

### 3.1 Effects of tillage intensity on soil properties

Soil chemical and physical properties were analyzed to assess the effect of tillage intensity on soil properties after running for almost 30 years. Results showed that total C and N in both depths sampled (0-10 and 10-20 cm) were twice as large in NT as compared to ST and MT (p<0.05, Table 1). Similarly, significantly highest concentrations of total soil P (PAR) and plant available K (KCAL) were found in NT (p<0.05, Table 1). Concentrations of other macro- and micronutrients were similar across tillage treatments (Table 1), thus data for plant available Cu, Fe, Mn, P, and Zn measured using Calcium chloride/DTPA (CAT) extraction are not shown, but are available on Zenodo (link will be added during revision process). Soil moisture contents were low in all treatments ranging between 2.6 to 3.2% with significantly lowest values in NT compared to MT and ST (p<0.05).

**Table 1:** Soil chemical properties standard tillage (ST), minimum tillage (MT) and no-tillage (NT) systems, in order of decreasing tillage intensity, collected from 0-10 cm, 0-20 cm, or individually from 0-10 cm and 10-20 cm as indicated. Extraction matrices used are abbreviated as follows: Calcium Chloride (CaCl2), Calcium-Acetate-Lactate (CAL), nitric acid and hydrochloric acid (AR, aqua regia). Values presented are estimated marginal means with 95% confidence intervals given in brackets. Model outputs are indicated with p ≤ 0.05 *, 0.01 **, 0.001 ***. Statistical analyses that failed ANOVA assumptions are reported in brackets. Different letters indicate significant differences between groups (α=0.05).

| Soil property | Depth | Tillage | ST | MT | NT |
| --- | --- | --- | --- | --- | --- |
| Moisture content (%) | 0-20 cm | $F_{2,6} = 8.7^*$ | 3.2 [3.0, 3.5] a | 3.1 [2.9, 3.4] a | 2.6 [2.4, 2.9] b |
| Soil pH<sub>CaCl2</sub> | 0-20 cm | $F_{2,6} = 0.3$ | 7.4 [7.0, 7.7] | 7.3 [7.0, 7.6] | 7.2 [6.9, 7.5] |
| Total C (%) | 0-10 cm | ($F_{2,8} = 9.6^{**}$) | 0.77 | 0.80 | 1.32 |
| Total C (%) | 10-20 cm | $F_{2,8} = 10.0^{**}$ | 0.73 [0.65, 0.84] b | 0.79 [0.70, 0.95] b | 1.24 [0.94, 1.34] a |
| Total N (%) | 0-10 cm | $F_{2,8} = 29.1^{***}$ | 0.06 [0.05, 0.08] b | 0.07 [0.06, 0.08] b | 0.12 [0.11, 0.13] a |
| Total N (%) | 10-20 cm | $F_{2,8} = 16.2^{**}$ | 0.06 [0.04, 0.07] b | 0.07 [0.05, 0.08] b | 0.11 [0.09, 0.12] a |
| C:N ratio | 0-10 cm | ($F_{2,8} = 0.6$) | 12.1 | 10.9 | 11.4 |
| C:N ratio | 10-20 cm | ($F_{2,8} = 0.2$) | 12.2 | 11.3 | 11.7 |
| P<sub>CAL</sub> (mg kg<sup>-1</sup>) | 0-20 cm | $F_{2,6} = 4.1$ | 692 [499, 884] | 677 [484, 869] | 958 [766, 1151] |
| P<sub>AR</sub> (mg kg<sup>-1</sup>) | 0-20 cm | $F_{2,6} = 68.1^{***}$ | 324 [310, 340] b | 335 [319, 352] b | 451 [425, 480] a |
| K<sub>CAL</sub> (mg kg<sup>-1</sup>) | 0-20 cm | $F_{2,6} = 42.9^{***}$ | 1592 [1255, 1960] b | 1497 [1168, 1857] b | 3477 [3008, 3973] a |
| K<sub>AR</sub> (mg kg<sup>-1</sup>) | 0-20 cm | ($F_{2,4} = 0.1$) | 5290 | 4624 | 4963 |
| Mg<sub>CaCl2</sub> (mg kg<sup>-1</sup>) | 0-20 cm | $F_{2,6} = 0.3$ | 1602 [966, 2238] | 1458 [822, 2093] | 1742 [1106, 2378] |
| Mg<sub>AR</sub> (mg kg<sup>-1</sup>) | 0-20 cm | $F_{2,6} = 3.0$ | 5688 [5061, 6315] | 4848 [4221, 5475] | 5501 [4874, 6128] |
| Ca<sub>AR</sub> (mg kg<sup>-1</sup>) | 0-20 cm | ($F_{2,6} = 0.6$) | 5896 | 4554 | 5822 |
| Fe<sub>AR</sub> (mg kg<sup>-1</sup>) | 0-20 cm | $F_{2,6} = 2.1$ | 18446 [16812, 20080] | 16743 [15109, 18377] | 16792 [15158, 18426] |
| Mn<sub>AR</sub> (mg kg<sup>-1</sup>) | 0-20 cm | $F_{2,6} = 0.3$ | 310 [268, 352] | 295 [253, 338] | 292 [249, 334] |
| Cu<sub>AR</sub> (mg kg<sup>-1</sup>) | 0-20 cm | $F_{2,6} = 0.8$ | 13 [12, 14] | 12 [11, 13] | 12 [11, 14] |
| Zn<sub>AR</sub> (mg kg<sup>-1</sup>) | 0-20 cm | $F_{2,6} = 0.5$ | 46 [41, 51] | 43 [37, 48] | 44 [39, 50] |

### 3.2 Impact of tillage intensity on soil biodiversity

To assess the impact of tillage intensity on soil biodiversity, a range of biological indicators based on molecular and classical morphological assessments were analyzed. Reducing tillage intensity (MT and NT) significantly increased the abundances of gene copy numbers of prokaryotes and eukaryotes as compared to ST (p<0.05, Table 2). The ratio of eukaryotic to prokaryotic gene copies in topsoil increased on average with reduction of tillage intensity from 0.12 in ST, over 0.13 in MT, to 0.15 in NT. Acari were found to be on average most abundant in NT. In particular, predatory acari were significantly most abundant under NT (Table 2). Of note was that total and fungivore acari were significantly more abundant in ST than in MT. Nematode abundance did not differ significantly across tillage treatments. On average, total abundance of nematodes, including their trophic groups such as bacterivores, fungivores, and herbivores increased consistently from ST over MT to NT (Table 2). This trend parallels the increase in prokaryotic and eukaryotic gene copies (Table 2). Regarding community richness and evenness of the respective communities, as assessed by the Shannon index, the bacterial diversity was found to be significantly reduced in NT (p<0.05), while the diversity of other communities did not differ significantly with tillage intensities (Table 2).

**Table 2:**
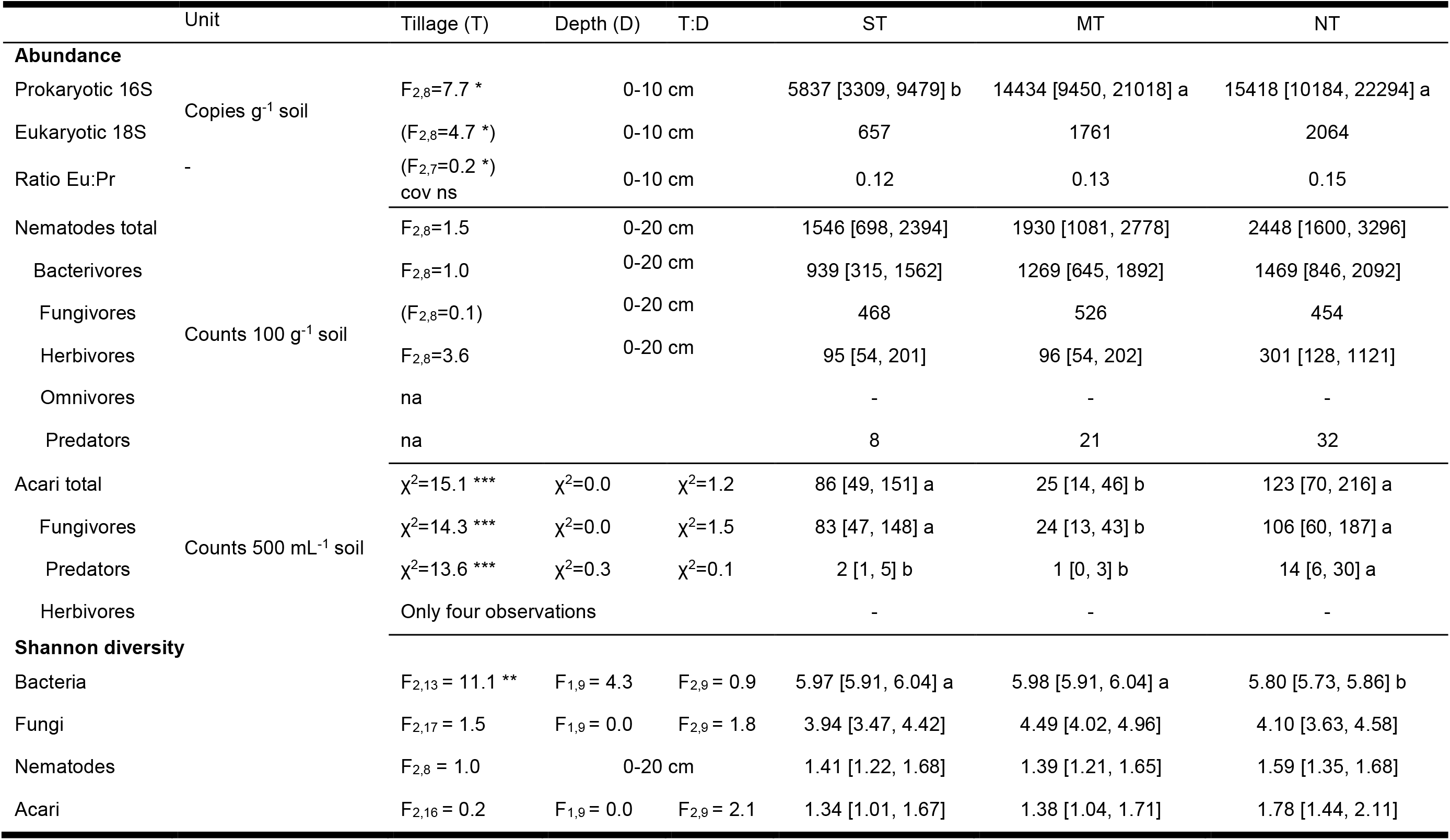
Abundances and diversity of soil bacteria, fungi, acari, and nematodes under standard tillage (ST), minimum tillage (MT) and no-tillage (NT) collected from 0-10 cm, 0-20 cm, or individually from 0-10 cm and 10-20 cm as indicated. Values presented are estimated marginal means with 95% confidence interval given in brackets. Model outputs are indicated with p ≤ 0.05 *, 0.01 **, 0.001 ***. Statistical analyses that failed ANOVA assumptions are reported in brackets. Different letters indicate significant differences between groups (α=0.05).

Multi-taxa community composition dissimilarities as calculated using the Bray-Curtis index were ordinated by Principal Coordinate Analysis (PCoA) for bacteria, fungi, acari and nematodes. This showed in 2D space a gradient along with tillage intensity (Figure 1 A-D). Permutational ANOVA supported this finding by indicating significant effects of tillage intensity (p<0.05) on bacterial, fungal and mite communities, but not on nematodes (Table A1). Soil depth was not found to be a significant factor in community composition.

Taking a closer look at the structure of bacterial communities across all treatments, Planctomycetota constituted the relatively most abundant phylum at 39.0% followed by Acidobacteria (15.8%), Actinobacteriota (14.4%), Proteobacteria (8.6%), Chloroflexi (8.5%), and Verrucomicrobiota (7.9%) (Figure A1). Besides Cyanobacteria, which showed the most pronounced tillage-induced shifts in relative abundance increasing from 0.5% under ST and MT to 3.8% under NT, several other taxa also changed (E-supplementary B1-B9).

Differential abundance analysis showed that 136 bacterial taxa were significantly associated with tillage and only one with depth (LinDA, FDR<5%). These candidates represented the maximum of 23% (NT, 0-10 cm) and minimum of 17% (MT, 10-20 cm) of 16S rRNA gene counts. The change profiles of these bacteria clustered according to Gap statistic and Tibshirani criterion into seven informative groups (Figure 2A). Clusters 7, 6 and 3, in order of decreasing mean log-fold change, were characterized by relative increases of bacteria under NT. A taxon from the genus *Tychonema* (order Cyanobacteriales) in cluster 7 was particularly prominent in terms of high relative abundance at 3.5% at 0-10 cm depth and 3.3% at 10-20 cm depth under NT and its high relative abundance when compared to ST and MT (Figure 2A cluster 7, Figure A37). The cluster 3 contained particularly relatively abundant taxa. These were taxa such as a Rhizobium from the genus *Microvirga* (family *Beijerinckiaceae*), Actinobacteria from the family *Micromonosporaceae* and *Blastocatellaceae*, and Planctomycetes from the Tepidisphaerales, and Isosphaerales.

Bacteria with the highest relative abundance under ST, and/or MT were placed in clusters 1, 2, 4, and 5 (Figure 2A, Figure A37). Particularly interesting in the context of plant responses was cluster 4 with significant enrichment of taxa from *Bacillus*, *Cohnella*, and *Streptomyces* under ST.

Taking a look at fungal community structure derived from PCR amplified ITS2 barcoding, the most relatively abundant phyla in descending order were Ascomycota (44%), Basidiomycota (25%), Chytridiomycota (5%), Mortierellomycota (4%), Glomeromycota (3%), Mucoromycota (1%), and Monoblepharomycota (0.3%) (Figure A38). Further, more basal fungal clades were also detected in lower relative abundance such as Aphelidiomycota, Basidiobolomycota, Olpidiomycota, Rozellomycota, and Zoopagomycota (E-supplementary F1-F9).

Differential abundance analysis indicated that 51 fungal taxa were significantly associated with tillage and only one with depth (FDR<5%). They represented a maximum of 32% (NT, 0-10 cm) and a minimum of 8% (MT, 0-10 cm) of the ITS2 sequences. The change profiles of fungi clustered into 6 informative groups (Figure 2B, Figure A63). Cluster 5, 1, and 2, in order of decreasing mean log-fold change, were associated with relative increases of taxa under NT (Figure 2B, Figure A63). In particular, *Coprinellus curtus* (order Agaricales, phyl. Basidiomycota) in cluster 5 exhibited a strong enrichment in topsoil, increasing from 0.004% under ST to 20.6% under NT conditions. Similarly, other Agaricales in cluster 1 and 2 including *Psathyrella* (fam. *Psathyrellaceae*), *Crepidotus* (fam. *Crepidotaceae*), and *Conocybe* (fam. *Bolbitiaceae*) increased under NT. Other taxa in cluster 5 were from the phylum Ascomycota and the classes Dothideomycetes (fam. *Didymosphaeriaceae* and *Phaeosphaeriacea*) and Leotiomycetes (fam. *Sclerotiniaceae*), amongst others (Figure A63). Clusters 3, 4, and 6 were characterized by strong relative decreases of fungi under NT (Figure 2B). Particularly cluster 3 contained relatively abundant fungal taxa, such as *Zymoseptoria triticii* (Ascomycota), *Aspergillus thesauricus* (Ascomycota), *Ascobolus* (Ascomycota), *Udeniomyces* (Basidiomycota) and several unassigned Ascomycotan families. Cluster 4 was represented by taxa with strongest relative increases under MT, containing a taxon from the genus *Fusarium* (Ascomycota), and further assigned taxa from the same class such as *Achaetomium*, and *Metacordyceps*.

### 3.3 Impact of tillage on soil functionality

Soil functionality was assessed by community level substrate utilization profiling (CLPP), potential enzymatic activity (EEA) of C, N, and P compounds, and calculations of C flow through the nematode community (Table 3). Tillage treatment significantly affected CLPP and EEA levels (p<0.01). Values for CLPP were significantly higher in NT and MT vs. ST (p<0.05), and for the former treatments it was also found that in top soil (0-10 cm) significantly higher values were measured vs. lower depth (10-20 cm). EEA of C, N, and P cycles was significantly higher in NT vs. MT and ST (p<0.05). On average, NT exhibited in all soil functionality measures the highest values, except for the fungal mediated OM decomposition derived from nematode composition.

**Table 3:**
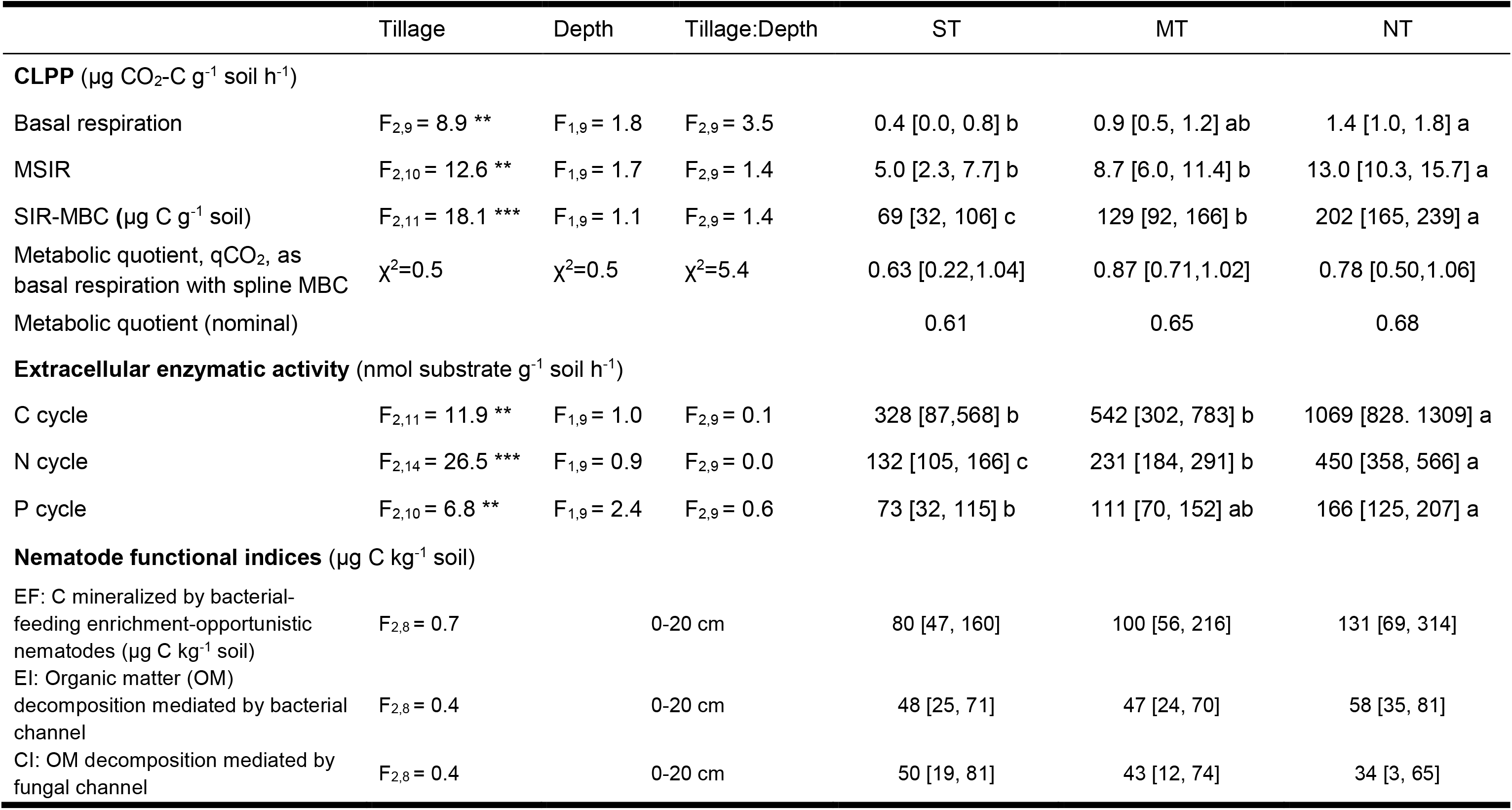
Soil functionality assessed under standard tillage (ST), minimum tillage (MT) and no-tillage (NT) analyzed in 0-20 cm depth or individually in 0-10 cm and 10-20 cm. Soil functionality consists of indices for community level physiological profile (CLPP with basal respiration and cumulated multiple substrate induced respiration (MSIR)), extracellular enzymatic activity (EEA) involved in carbon (C, beta-glucosidase, cello-biohydrolase, xylosidase), nitrogen (N, N-acetyl-glucosidase, L-aminopeptides), and phosphorous (P, alkaline and neutral phosphatases) cycling, and nematode functional indices (enrichment metabolic footprint (EF), enrichment Index (EI), and channel Index (CI)). Values reported are estimated marginal means with 95% confidence interval given in brackets. Model outputs are indicated with p ≤ 0.05 *, 0.01 **, 0.001 ***. Different letters indicate significant differences between groups (α=0.05).

|  | Tillage | Depth | Tillage:Depth | ST | MT | NT |
| --- | --- | --- | --- | --- | --- | --- |
| **CLPP ($\mu\text{g CO}_2\text{-C g}^{-1} \text{ soil h}^{-1}$)** |  |  |  |  |  |  |
| Basal respiration | $F_{2,9} = 8.9^{**}$ | $F_{1,9} = 1.8$ | $F_{2,9} = 3.5$ | 0.4 [0.0, 0.8] b | 0.9 [0.5, 1.2] ab | 1.4 [1.0, 1.8] a |
| MSIR | $F_{2,10} = 12.6^{**}$ | $F_{1,9} = 1.7$ | $F_{2,9} = 1.4$ | 5.0 [2.3, 7.7] b | 8.7 [6.0, 11.4] b | 13.0 [10.3, 15.7] a |
| SIR-MBC ($\mu\text{g C g}^{-1} \text{ soil}$) | $F_{2,11} = 18.1^{***}$ | $F_{1,9} = 1.1$ | $F_{2,9} = 1.4$ | 69 [32, 106] c | 129 [92, 166] b | 202 [165, 239] a |
| Metabolic quotient, $q\text{CO}_2$, as<br>basal respiration with spline MBC | $\chi^2=0.5$ | $\chi^2=0.5$ | $\chi^2=5.4$ | 0.63 [0.22,1.04] | 0.87 [0.71,1.02] | 0.78 [0.50,1.06] |
| Metabolic quotient (nominal) |  |  |  | 0.61 | 0.65 | 0.68 |
| **Extracellular enzymatic activity (nmol substrate $\text{g}^{-1} \text{ soil h}^{-1}$)** |  |  |  |  |  |  |
| C cycle | $F_{2,11} = 11.9^{**}$ | $F_{1,9} = 1.0$ | $F_{2,9} = 0.1$ | 328 [87,568] b | 542 [302, 783] b | 1069 [828, 1309] a |
| N cycle | $F_{2,14} = 26.5^{***}$ | $F_{1,9} = 0.9$ | $F_{2,9} = 0.0$ | 132 [105, 166] c | 231 [184, 291] b | 450 [358, 566] a |
| P cycle | $F_{2,10} = 6.8^{**}$ | $F_{1,9} = 2.4$ | $F_{2,9} = 0.6$ | 73 [32, 115] b | 111 [70, 152] ab | 166 [125, 207] a |
| **Nematode functional indices ($\mu\text{g C kg}^{-1} \text{ soil}$)** |  |  |  |  |  |  |
| EF: C mineralized by bacterial-<br>feeding enrichment-opportunistic<br>nematodes ($\mu\text{g C kg}^{-1} \text{ soil}$) | $F_{2,8} = 0.7$ | | 0-20 cm | 80 [47, 160] | 100 [56, 216] | 131 [69, 314] |
| EI: Organic matter (OM)<br>decomposition mediated by bacterial<br>channel | $F_{2,8} = 0.4$ | | 0-20 cm | 48 [25, 71] | 47 [24, 70] | 58 [35, 81] |
| CI: OM decomposition mediated by<br>fungal channel | $F_{2,8} = 0.4$ | | 0-20 cm | 50 [19, 81] | 43 [12, 74] | 34 [3, 65] |

### 3.4 Impact of tillage on wheat performance

At flowering stage, total aboveground biomass of individually weighed wheat plants was significantly affected by tillage (p<0.05), decreasing from ST, over MT to NT by more than 2- fold (Table 4). This trend was seen alike when resolving for the compartments head, leaves and stem. The root biomass was not significantly different between tillage treatments at this growth stage (p>0.05), however, on average, wheat under ST exhibited higher root biomass, followed by NT and lowest under MT (Table 4). The root:shoot ratio was significantly higher under NT compared to MT and ST (p<0.05). The wheat root-to-soil interface was resolved further by characterizing root traits including number of root tips, branch points, total length/area/volume, diameter of roots and root orientation (Table 4). Only root median diameter was significantly higher under NT compared to ST and MT (p<0.05). Not significant, but on average higher under NT were also root average diameter, root maximum diameter and large (>0.2 mm) root diameter length. For wheat roots under ST, the number of root tips, root surface area and total root volume were on average highest. And for wheat roots under MT, number of root branch points, branching frequency, total root length, the proportion of root length in fine (<0.1 mm) and medium (0.1-0.2 mm) range diameter classes were on average highest, while root angle orientation was lowest (Table 4).

**Table 4:** Wheat growth performance at flowering stage in standard tillage (ST), minimum tillage (MT) and no-tillage (NT) systems. The aboveground parameters refer to individual plants collected from a circular area 9 cm in diameter and the corresponding root parameters from a depth of 20 cm. Values are estimated marginal with 95% confidence interval given in brackets. Model output are indicated with p ≤ 0.05 *, 0.01 **, 0.001 ***. Statistical analyses that failed ANOVA assumptions are reported in brackets. Different letters indicate significant differences between groups (α=0.05).

| Wheat growth parameter | Tillage | ST | MT | NT |
| --- | --- | --- | --- | --- |
| **Plant biomass (per cylinder area)** |  |  |  |  |
| Aboveground biomass (g) | ($F_{2,6} = 5.1^*$) | 27 | 15 | 11 |
| Head (spike) biomass (g) | $F_{2,6} = 3.2$ | 9.0 [5.4, 18.7] | 5.1 [3.5, 8.2] | 4.6 [3.2, 7.3] |
| Leaf biomass (g) | $F_{2,6} = 6.4^*$ | 4.7 [2.8, 7.8] a | 2.6 [1.5, 4.3] ab | 1.6 [1.0, 2.7] b |
| Stem biomass (g) | ($F_{2,6} = 6.5^*$) | 13.7 | 7.6 | 4.9 |
| Root biomass (g) | $F_{2,6} = 0.3$ | 0.86 [0.50, 1.21] | 0.70 [0.35, 1.06] | 0.80 [0.44, 1.15] |
| Root to shoot ratio | $F_{2,6} = 5.9^*$ | 0.03 [0.01, 0.05] b | 0.04 [0.02, 0.06] b | 0.07 [0.05, 0.09] a |
| **Root system characteristics (per cylinder volume)** |  |  |  |  |
| Root tips (counts) | $F_{2,6} = 0.1$ | 51928 [20091, 123600] | 42006 [15908, 101778] | 44599 [16994, 107515] |
| Root branch points (counts) | ($F_{2,6} = 0.9$) | 70995 | 79941 | 56659 |
| Root branching frequency (freq. mm<sup>-1</sup>) | ($F_{2,6} = 1.5$) | 1.10 | 1.18 | 1.00 |
| Root network area (mm<sup>2</sup>) | $F_{2,6} = 0.0$ | 12906 [8345, 17466] | 12792 [8231, 17352] | 12247 [7686, 16808] |
| Total root volume (mm<sup>3</sup>) | $F_{2,6} = 0.1$ | 12272 [6303, 18240] | 10814 [4845, 16783] | 10530 [4561, 16499] |
| Total root surface area (mm<sup>2</sup>) | $F_{2,6} = 0.1$ | 69525 [42125, 96925] | 67592 [40192, 94992] | 62697 [35298, 90097] |
| Total root length (mm) | ($F_{2,6} = 0.6$) | 65102 | 71601 | 56486 |
| Root average diameter (mm) | ($F_{2,6} = 11.5^{**}$) | 0.34 | 0.29 | 0.35 |
| Root median diameter (mm) | $F_{2,6} = 12.3^{**}$ | 0.22 [0.21, 0.24] b | 0.20 [0.18, 0.22] b | 0.25 [0.24, 0.27] a |
| Root maximum diameter (mm) | $F_{2,6} = 0.8$ | 3.40 [2.74, 3.98] | 3.42 [2.76, 4.00] | 3.78 [3.17, 4.33] |
| Root length diameter range <0.1 mm (mm) | $F_{2,6} = 1.2$ | 9114 [4922, 18404] | 11331 [5964, 23665] | 6401 [3597, 12281] |
| Root length diameter range 0.1-0.2 mm (mm) | ($F_{2,6} = 2.1$) | 21762 | 24884 | 16159 |
| Root length diameter range >0.2 mm (mm) | $F_{2,6} = 0.0$ | 33207 | 33127 | 33299 |
| **Root crown analysis** |  |  |  |  |
| Average root orientation (°, deg.) | ($F_{2,6} = 1.6$) | 46.0° | 43.9° | 45.2° |
| Shallow angle frequency 0-30° (%) | $F_{2,6} = 1.6$ | 31.2 [27.1, 35.3] | 35.4 [31.3, 39.5] | 33.6 [29.5, 37.7] |
| Medium angle frequency 30-60° (%) | ($F_{2,6} = 0.2$) | 32.6 | 32.3 | 31.9 |
| Steep angle frequency 60-90° (%) | ($F_{2,6} = 1.5$) | 35.1 | 32.0 | 34.2 |

At harvest, wheat yields and thousand kernel weight were similar across treatments (Figure 5). In contrast, grain N concentration (%) were significantly higher under ST compared to those of NT (F2,6=5.6, p<0.05, Table 5). In correspondence to the highest N content in grains grown under ST, the grain B vitamin content for Thiamine (B1) and Riboflavin (B2) were found to be significantly highest under ST compared to NT and MT. Interestingly, grain Ca and Si contents were significantly highest in wheat grown under MT, while other macronutrients such as C, P, K, S, Mg and micronutrients did not differ between tillage treatments. Regarding technological grain properties, also the distribution of starch size classes significantly differed between treatments with a higher abundance of A-type starches (>10 µm) in ST compared to NT (p< 0.01) and a higher abundance of B-type starches (2<x<10 µm) in NT compared to ST (p< 0.01) (Table 5). In contrast, the abundance of gliadins and glutenins was similar across treatments (Table 5).

**Table 5:** Wheat grain yield, nutritional and technological quality in standard tillage (ST), minimum tillage (MT) and no-tillage (NT) systems. Values presented are estimated marginal means with 95% confidence interval given in brackets. Model output are indicated with p ≤ 0.05 *, 0.01 **, 0.001 ***. Statistical analyses that failed DHARMa diagnostics are reported in brackets. Different letters indicate significant differences between groups (α=0.05).

|  | Tillage | ST | MT | NT |
| --- | --- | --- | --- | --- |
| **Harvest parameters** |  |  |  |  |
| Grain yield (dt ha<sup>-1</sup>) | F<sub>2,6</sub> = 0.8 | 7.3 [4.8, 9.7] | 5.5 [3.0, 7.9] | 6.0 [3.6, 8.5] |
| Thousand kernel weight (g) | F<sub>2,6</sub> = 1.8 | 17.1 [15.1, 19.1] | 17.7 [15.7, 19.7] | 19.3 [17.3, 21.3] |
| **Nutrients** |  |  |  |  |
| C (g kg<sup>-1</sup>) | F<sub>2,6</sub> = 2.0 | 430 [427, 433] | 429 [426, 432] | 427 [424, 430] |
| N (g kg<sup>-1</sup>) | F<sub>2,6</sub> = 5.6 * | 31 [28, 34] a | 28 [25, 31] ab | 25 [22, 28] b |
| S (g kg<sup>-1</sup>) | F<sub>2,6</sub> = 0.8 | 1.8 [1.7, 2.0] | 1.7 [1.5, 1.9] | 1.7 [1.6, 1.9] |
| P (g kg<sup>-1</sup>) | (F<sub>2,6</sub> = 0.8) | 4.06 | 3.88 | 3.92 |
| K (g kg<sup>-1</sup>) | (F<sub>2,6</sub> = 4.9) | 5.43 | 5.72 | 5.19 |
| Mg (g kg<sup>-1</sup>) | F<sub>2,6</sub> = 1.2 | 1.44 [1.33, 1.55] | 1.40 [1.29, 1.51] | 1.34 [1.23, 1.45] |
| Ca (mg kg<sup>-1</sup>) | (F<sub>2,6</sub> = 12.9 **) | 493 | 923 | 410 |
| Cl (mg kg<sup>-1</sup>) | F<sub>2,6</sub> = 4.4 | 724 [613, 835] | 906 [795, 1017] | 768 [658, 879] |
| Si (mg kg<sup>-1</sup>) | F<sub>2,6</sub> = 6.7 * | 191 [53, 329] b | 409 [271, 547] a | 131 [-7, 268] b |
| Fe (mg kg<sup>-1</sup>) | (F<sub>2,6</sub> = 2.6) | 37 | 30 | 25 |
| Mn (mg kg<sup>-1</sup>) | (F<sub>2,6</sub> = 3.6) | 40 | 39 | 34 |
| Zn (mg kg<sup>-1</sup>) | F<sub>2,6</sub> = 0.5 | 30 [25, 38] | 28 [24, 35] | 27 [23, 33] |
| Cu (mg kg<sup>-1</sup>) | F<sub>2,6</sub> = 0.8 | 5.7 [5.1, 6.6] | 5.3 [4.7, 6.0] | 5.3 [4.7, 6.0] |
| Ni (mg kg<sup>-1</sup>) | F<sub>2,6</sub> = 1.3 | 0.5 [0.4, 1.0] | 0.4 [0.3, 0.6] | 0.5 [0.3, 0.9] |
| Mo (mg kg<sup>-1</sup>) | F<sub>2,6</sub> = 0.9 | 0.2 [0.0, 0.3] | 0.1 [0.0, 0.3] | 0.2 [0.1, 0.4] |
| Co (mg kg<sup>-1</sup>) | F<sub>2,6</sub> = 1.0 | 0.03 [0.02, 0.05] | 0.03 [0.01, 0.04] | 0.04 [0.03, 0.06] |
| **B vitamins** |  |  |  |  |
| Thiamine (B1) (µg 100 g<sup>-1</sup>) | F<sub>2,6</sub> = 12.5 ** | 6.4 [5.6, 7.6] a | 4.7 [4.2, 5.3] b | 4.5 [4.1, 5.1] b |
| Riboflavin (B2) (µg 100 g<sup>-1</sup>) | F<sub>2,6</sub> = 6.3 * | 1.1 [0.9, 1.3] a | 0.8 [0.6, 1.0] b | 0.7 [0.5, 0.9] b |
| Niacine (B3) (µg 100 g<sup>-1</sup>) | (F<sub>2,6</sub> = 2.3) | 14.6 | 12.1 | 11.3 |
| Pantothenic acid (B5) (µg 100 g<sup>-1</sup>) | (F<sub>2,6</sub> = 5.5 *) | 9.1 | 7.6 | 6.3 |
| Pyridoxine (B6) (µg 100 g<sup>-1</sup>) | (F<sub>2,6</sub> = 6.9 *) | 2.9 | 2.2 | 2.1 |
| Folate (B9) (µg 100 g<sup>-1</sup>) | F<sub>2,6</sub> = 4.6 | 0.6 [0.5, 0.7] | 0.4 [0.3, 0.5] | 0.4 [0.3, 0.5] |
| **Technological properties** |  |  |  |  |
| A-type starch (>10 µm) (%) | F<sub>2,6</sub> = 11.2 ** | 93.0 [91.5, 94.5] a | 91.3 [89.9, 92.8] a | 89.0 [87.5, 90.5] b |
| B-type starch (2-10 µm) (%) | (F<sub>2,6</sub> = 11.6 **) | 6.9 | 8.6 | 10.9 |
| C-type starch (< 2 µm) (%) | F<sub>2,6</sub> = 0.4 | 0.0 [0.0, 0.1] | 0.0 [0.0, 0.1] | 0.1 [0.0, 0.1] b |
| Glutenins (%) | F<sub>2,6</sub> = 3.7 | 1.38 [1.19, 1.56] | 1.14 [0.96, 1.33] | 1.11 [0.92, 1.29] |
| Gliadins (%) | F<sub>2,6</sub> = 4.0 | 0.67 [0.55, 0.80] | 0.63 [0.50, 0.76] | 0.47 [0.35, 0.60] |

## 4. Discussion

### 4.1 Reducing tillage intensity enhanced soil chemical and biological properties

The increases in total soil C, N, P, and K under NT are consistent with previously reported results of increased organic matter measures from the same field experiment (Martin-Rueda et al., 2007, Martín-Lammerding et al., 2013, 2011), as well as the results of the meta-study by Haddaway et al. (2017).

Unexpectedly, the soil moisture content in NT was the lowest (Table 1). This was not reported in previous studies from the same site, where NT exhibited higher soil water content at flowering stage. Indeed, it was argued that reduced tillage practices are also water saving practices (Santin-Montanya et al., 2020). Further, a higher organic matter content would in principle enable higher water holding capacity from aggregate formation and pore space. The differing water status under NT in this study could derive from temporal water dynamics in connection with sampling time. Co-occurring plants (e.g. weeds) were found to influence soil water status by complex interactions with seasonal total rainfall, legacy effects over years, and with short-term weather/sampling time (Gandia et al., 2021, Santin-Montanya et al., 2020).

Corresponding to the higher C content in NT, our results showed increased abundances of prokaryotic and eukaryotic gene copies, and on average an increase in acari and nematodes (Table 2) as well as higher SIR microbial biomass (Table 3). In addition, we observed higher basal respiration, substrate utilization rates and enzymatic activities under NT (Table 3), which are indicators for higher microbial biomass. Our results are consistent with the meta-analysis of Chen et al. (2020) who found a positive effect of reduced tillage on the abundance of bacteria and fungi as well as an increase in total C and N that might be linked with the increase of SOC under NT (Lori et al. 2017). Similarly, a positive association of soil biological indicators with decreasing tillage intensity was observed in the meta-analysis of Nunes et al. (2020). However, despite that MT exhibited similar prokaryotic and eukaryotic gene copies to NT, the functional activity levels were significantly lower (Table 3). This could result from significantly different bacterial and fungal microbial communities (Figure 1, Figure 2), that would be expected to result in different functionalities and activity levels. Indeed, a previous study from the same field experiment showed an increase in soil basal respiration, microbial biomass C and β- glucosidase activity under NT (Martín-Lammerding et al., 2015). The observed increased in eukaryotic:prokaryotic qPCR gene copy ratio (Table 2) from ST, over MT, to NT is consistent with findings that reduced tillage practices are resulting in increased total microbial biomass and fungal proportions (Six et al., 2006, Piazza et al. 2019). In contrast, van Groenigen et al., (2010) and Murugan et al., (2014) did not find a change in the proportion of bacteria and fungi under reduced tillage, but an increase in saprotrophic fungi, which is in line with the increase in relative abundance of taxa with saprotrophic/seed nutrition under NT observed in this study (discussed in section 4.3). The studies of van Groenigen et al., (2010) and Murugan et al., (2014) highlighted the importance of differing organic matter degrading capabilities of fungal groups and that this can have an effect on the organic matter dynamics. An increased proportion of fungal decomposers could also be an alternative explanation, apart from stress and disturbance (Wardle and Ghani, 1995, Anderson and Domsch, 2010), for the here observed increase of the metabolic quotient from ST, over MT, to NT (Table 3).

### 4.2 Tillage intensity shapes soil biota community structure

As hypothesized, tillage intensity significantly shaped bacterial, fungal, and acarid communities, while this could not be shown for nematodes (Figure 1, Table A1). Largely this is consistent with previous studies documenting differences in the community structures of microorganisms (Hartman et al., 2018; Kraut-Cohen et al., 2020; Orrù et al., 2021; Wang et al., 2020), nematode (Puissant et al., 2021), and acari (Betancur-Corredor et al., 2022) between ploughed, reduced tillage, and no-tillage systems. A key factor driving these differences may be the increase in SOC with reduced tillage intensity and the associated increase in microbial community size and activity (Lori et al. 2017, Ramírez et al., 2020). Moreover, tillage intensity has been demonstrated to impact various soil properties, including the stability and size of aggregates and bulk density (Li et al., 2019; Nunes et al., 2020; Mondal and Chakraborty, 2022). These properties are also influenced by soil microbes, which in turn might shape the other soil community by bottom-up forces (Hartmann and Six, 2022, Philippot et al., 2024). Since tillage intensity affects numerous relative abundance changes at the microbial level, even contrasting within families and genera, an extensive ecological description of bacterial and fungal clades with changes under tillage intensities can be found in the supplementary (supplementary data 1.1.1 for bacteria and 1.4.1 for fungi). From a biological standpoint, these findings align with ecological theory. Tillage-induced changes in nutrient availability (quality and quantity of organic matter, root traits) and further soil properties lead to shifts in multidimensional niche space. Indeed, the competitive exclusion principle states that if two species use the exact same resources, the one using the resources more efficiently will exclude the other (Gauss 1934). Therefore, coexistence of such a high number of bacterial and fungal taxa (hundreds to thousands) found on a small volume that DNA was extracted from, and with many taxa using similar resources is only possible, if substantial niche differences evolve between closely related taxa or taxa using similar resources (the multidimensional niche: Hutchinson, 1959). Of these niche dimensions a high degree of micro-spatial separation and environmental variability could play a role (Horner-Devine et al., 2004). Indeed, high resolution of soil bacterial translational dynamics have shown that competition within nutritional guilds was strongest and affected by specialist-directed nutritional substances and abundance of competitors (Moyne et al., 2023). For a more in-depth discussion, we will focus on two key aspects: the formation of soil biocrust (explained later) under NT and the dynamics of fungal plant pathogens and fungal antagonists.

Under NT, the most pronounced gene copy increase in bacteria was from a taxon in the *Tychonema* genus (a Cyanobacterium) (Figure 2A, cluster 7). This taxon assignment is monophyletic with *Microcoleus vaginatus* (Strunecky et al., 2023, Zhang et al., 2016), which are well described for their importance to soil biocrust formation (Büdel, 2005, Powell et al., 2015). Given the increase in total 16S gene copies under NT (Table 2), the increase in cyanobacteria may hint at their role in soil C and N accumulation (Table 1). This could contribute to soil biocrust formation, a less studied aspect in arable farming. Biological soil crusts are linkages of mineral soil particles with a community of photo-autotrophic cyanobacteria, algae, lichens, bryophytes, and associated heterotrophic fungi and prokaryotes that often develop in arid regions or on early habitats such as rocks. The organismal composition is structured by local degree of abiotic stresses such as aridity and solar radiation (Büdel, 2005, Grishkan et al., 2019, Hu et al., 2003). In arable farming, where the soil surface is often disturbed and exposed when bare, particularly in semi-arid regions, the pioneering role of biocrusts and microbial photoautotrophs is interesting in regard to their reported contributions to SOM built-up, N2-fixation, and climate resilience (Grishkan et al., 2019, Powell et al., 2015, Maddigan, et al., 2013, Büdel, 2005, Hu et al., 2003, Mazor et al., 1996). Besides *Tychonema*, we also detected other cyanobacteria some associated to lichens or algae (from the order Chaetothyriales such as *Bradymyces*, *Exophiala*, Lücking et al., 2009), as well as micro-fungal and bacterial heterotrophs that were reported to occur in soil biocrusts (supplementary data 1.4). Thus, our results suggest that the development of biocrusts is possible under NT, whereas ST and MT might exhibit a “critical” degree of disturbance (Figure A65).

Focusing on fungal pathogen – antagonist interactions, we observed a relative decrease of two fungal plant pathogens, *Zymoseptoria tricii* (Ascomycota, class Dothideomycetes, Quadvlieg et al., 2011) and a *Fusarium* sp. (Ascomycota, class Sordariomycetes, Lombard et al., 2015) under NT compared to ST (clusters 3 and 4 in Figure 2B). In contrast, other organisms in the class Dothideomycetes relatively increased under NT including *Spegazzinia radermacherae,* a saprotroph living on fallen seed pods (Jayasiri et al., 2019) and *Sclerostagonospora*, a pathogen or saprotroph of various monocotyledons and dicotyledons (Phookamsak et al., 2014) as well as other saprotrophs including *Keissleriella* (*cirsii*) and *Clarireedia bennettii* (class Leotiomycetes), the latter known as pathogen of C3 turfgrasses (Salgado-Salazar et al., 2018). The increase of some fungal plant saprotrophs and seed/pollen attacking taxa under NT might have resulted from an increased frequency of co-occurring weeds, weed richness, and weed seed bank density observed in soil under reduced tillage (MT and NT) (Santín-Montanyá et al., 2020, 2018, 2016). The increase of herbivore nematodes under NT (Table 2) supports this assumption.

On the flipside, the most notable fungal plant pathogen antagonist strongly increased under NT was a *Coprinellus curtus* (Cluster 5, Figure 2, Figure 64). As a saprotroph, *C. curtus* could be expected to grow well at higher OM levels under NT. In addition, a relevant agricultural isolate of *C. curtus* showed antagonistic behavior against several plant pathogens including *Fusarium oxysporum* and *Rhizoctonia solani* (Nakasaki et al., 2007), which might explain the observed decrease of *Fusarium* sp. in the present study. A multifunctionality of other OM- degrading Basidiomycetes that relatively increased under NT, such as *Psathyrella*, *Crepidotus*, and *Conocybe,* may be their plant pathogen suppressive effect, either by direct interference or by indirect competition. It is speculated here that higher OM levels under NT may allow these fungi to thrive.

### 4.3 Impaired wheat growth and nutritional quality under reduced tillage despite enhanced soil biological indicators

Wheat growth, average wheat yields, and the nutritional quality indicators grain N content and grain vitamin thiamine (B1) and riboflavin (B2) levels, were highest under ST and lowest under NT (Table 5). Our findings are consistent with the meta-analysis by Pittelkow et al. (2015), who reported increased wheat yield and N uptake under more intensive tillage practices. As N is a crucial element in the synthesis of proteins and enzymes involved in the production of B vitamins, it was expected that vitamin B levels would show similar patterns, directly correlating with grain N content. In addition, studies have demonstrated that B vitamins are differentially affected by genotype and environmental factors such as temperature, precipitation, and soil conditions (Shewry et al., 2011; Batifoulier et al., 2006). These factors may further explain the observed differences in B vitamin levels induced by tillage practices. In addition, this is the first study reporting about tillage-induced changes in wheat technological properties as reflected by a shift in A- and B-type starch distribution and on average higher content of Gliadins and Glutenins under ST. As recently reviewed by Guo et al. (2023) the distribution of A- and B-type starch are important technological quality indicators as they strongly influence the quality of dough and final products.

Explanations for wheat performance may trace to root system architecture, here measured at the flowering stage. The roots data (Table 4) showed that the root-to-sprout ratio was highest under NT, indicating resource allocation for soil penetration and root thickening in wheat in compacted soils (Munoz-Romero et al., 2010, Watt et al., 2005, Pandey et al., 2021). Indeed, wheat roots under NT had the significantly highest median root diameter, and on average higher root average diameter, root maximum diameter, and larger root diameter, which were consistent with descriptions of soil compaction effects on wheat (Munoz-Romero et al., 2010, Atwell, 1989, Watt et al., 2005). The better performance of wheat under ST is reflected by the characteristics of its root system with an increase in root area, root volume and the number of root tips (Table 4). These characteristics, in turn, are reported to be linked to higher N mineralization potential through: 1) root exudation associated with number of actively growing root tips, considered as hot spots of microbial abundance and activity in the rhizosphere, 2) root surface or length being indicative of the potential for nutrient exploration and establishment of biological associations, and 3) proportion of fine roots influencing the water uptake capacity and the total root surface (Freschet et al., 2021, Delaplace et al., 2015, Kuzyakov and Blagodatskaya, 2015, Watt et al., 2009, Watt et al., 2008, McCully, 1999). Roots influence microbes and their nutrient mobilization potential by supplying energy in what is summarized as the “Rhizosphere effect” (Kuzyakov and Blagodatskaya, 2015, Nguyen, 2009, Hodge et al., 2000). This could explain the significantly higher proportion of rhizosphere-associated or potentially plant growth promoting taxa under ST of *Bacillus* and *Cohnella* (Firmicutes), and Actinomycetes such as *Streptomyces* and *Luedemanella*.

Wheat grown under MT did not perform as well in regards to grain yield and quality as wheat grown under ST (Table 5). In the former, root traits were characterized by on average highest values for root branch points, branching frequency, and lowest root angle orientation with higher frequency of shallow angled roots (Table 4). Considering that MT only loosens the soil to a depth of about 15 cm, could mean easier soil penetration during the early growth stages of wheat, and negative effects during later growth stages associated with a soil compaction layer. At the same time, the root angle influences the vertical distribution of the roots (and their water uptake potential in deeper soil layers), which is influenced by soil compaction and could affect later wheat growth (Freschet et al., 2021, Delaplace et al., 2015, Watt et al., 2009, Watt et al., 2008, Watt et al., 2005). Sampling throughout the growing season as done in Munoz- Romero et al., (2010) for the comparison of NT versus ST could clarify whether the intermediate growth of wheat under MT is due to early growth promotion by easy soil penetration and later growth inhibition by soil compaction. In addition, legacy effects of long-term NT/MT on root characteristics might differ to conditions with recent/alternating tillage management practices.

Despite higher concentration of available soil nutrients (Table 1), increased abundances of prokaryotes, eukaryotes, nematodes and acari (Table 2), substrate mineralization potential and enzyme cycling activity under NT (i.e. N cycling, Table 3), wheat performance was lower than under ST (Table 4, Table 5). This is striking as the action of microbial grazers as well as the diurnal cycles of water uptake in the rhizosphere should liberate nutrients from the microbial pool for plant uptake (Snapp and Drinkwater, 2007, Clarholm, 1985) enhancing nutrient uptake under NT. Thus, other factors such as nutrient competition by higher weed pressure, microbial immobilization and/or slower SOM mineralization resulting in a reduced N availability or uptake might have limited wheat growth and grain N content under reduced tillage (Soane et al. 2012, Cooper et al. 2016, Hofmeijer et al. 2019, Kuzyakov and Blagodatskaya, 2015, Nguyen, 2009, Hodge et al., 2000). Here we have outlined possible physical and biological reasons for the decrease in wheat performance under reduced soil tillage intensity, however, the exact causes require further research.

## 5. Conclusion

Our study highlights the important role of tillage in shaping soil biodiversity and functionality. Varying tillage intensity is reflected in distinct shifts in both the structural and functional communities within the soil. Our findings provide evidence that the formation of biocrusts under NT may contribute to the accumulation of OM in upper soil layers. The lower growth performance and nutritional quality of wheat grains, despite enhanced soil chemistry and biology, points to underlying root structure problems that might have limited wheat growth and nutrient uptake. Addressing these issues may require the application of additional measures such as the cultivation of deep rooting inter-crops or complementary tillage practices to improve wheat production under no-tillage conditions. Alternatively, research into wheat varieties with root systems adapted to no-tillage conditions may offer promising solutions. In addition, the study showed that tillage intensity has the potential to alter technological grain properties and should be considered in the subsequent use of the wheat for production.

## Acknowledgments

This research is part of the BIOFAIR project (http://www.biofair.uliege.be) funded through the 2019-2020 BiodivERsA joint call for research proposals, under the BiodivClim ERA-Net COFUND programme, and with the funding organisations Agence Nationale de la Recherche (ANR, France; ANR-20-EBI5-0002), Agencia Estatal de Investigación (AEI, Spain; PCI2020- 120713-2), Deutsches Zentrum fuer Luft- und Raumfahrt Projektträger (DLR-PT, Germany), Fonds de la Recherche Scientifique (FNRS, Wallonia, Belgium; R.8001.20), Fonds voor Wetenschappelijk Onderzoek - Vlaanderen (FWO, Flanders, Belgium; FWO ERA-NET G0H7320N) & Schweizerischer Nationalfonds (SNF, Switzerland; 31BD30_193869). Data produced and analyzed in this paper were generated in collaboration with the Genetic Diversity Centre (GDC), ETH Zurich, and Core Facility, University of Hohenheim.

## Data availability

The microbiome data presented in the study are deposited in the NCBI Sequence Read Archive (SRA) repository, accession number xxx. All other data are available on Zenodo (link will be added during revision).

