## Supplementary Figures and Tables for "Tensions in tillage: Reduction in tillage intensity associates with lower wheat growth and nutritional grain quality despite enhanced soil biological indicators"

### 1. Supplementary data and figures

#### 1.1 Taxonomic composition of bacteria

The phylum with the highest average relative abundance of 16S amplicons in extracts of this Spanish soil (calcic Haploxeralf) were Planctomycetota (39.0%). Further phyla in descending relative abundance of 16S amplicons were Acidobacteria (15.8%), Actinobacteriota (14.4%), Proteobacteria (8.6%), Chloroflexi (8.5%), and Verrucomicrobiota (7.9%).

Cyanobacteria exhibited an increase from 0.5% in standard and reduced tillage (ST and MT) to 3.8% in no-tillage (NT).

Firmicutes amplicons were detected in low copy numbers at ca. 0.4%. The low relative abundance of Firmicutes may be often the case in arable Mediterranean soil, as other studies, some of which were also from the wheat rhizosphere/bulk soil, reported this (Lasa et al., 2019, dal Cortivo et al., 2020, Visioli et al., 2020).

Lasa et al., (2019) used the MoBio RNA PowerSoil DNA and RNA co-extraction kit with 15 mL tubes, silica beads, and using a vortex at max setting for 15 minutes, and amplifying the V3-V4-V5 region with primers U341F and U926R (Baker et al., 2003).

Dal Cortivo et al., (2020) used the MP Biomedicals FastDNA spin kit for soil using bead beating tubes, extracting from 0.5 g soil, lysing for 40 s in MP FastPrep homogenizer, and amplifying the 16S V3 region with primers Probio_Uni and Probio_Rev (Milani et al., 2013).

Visioli et al., (2020) extracted DNA and amplified 16S V3 region as in dal Cortivo et al., (2020).

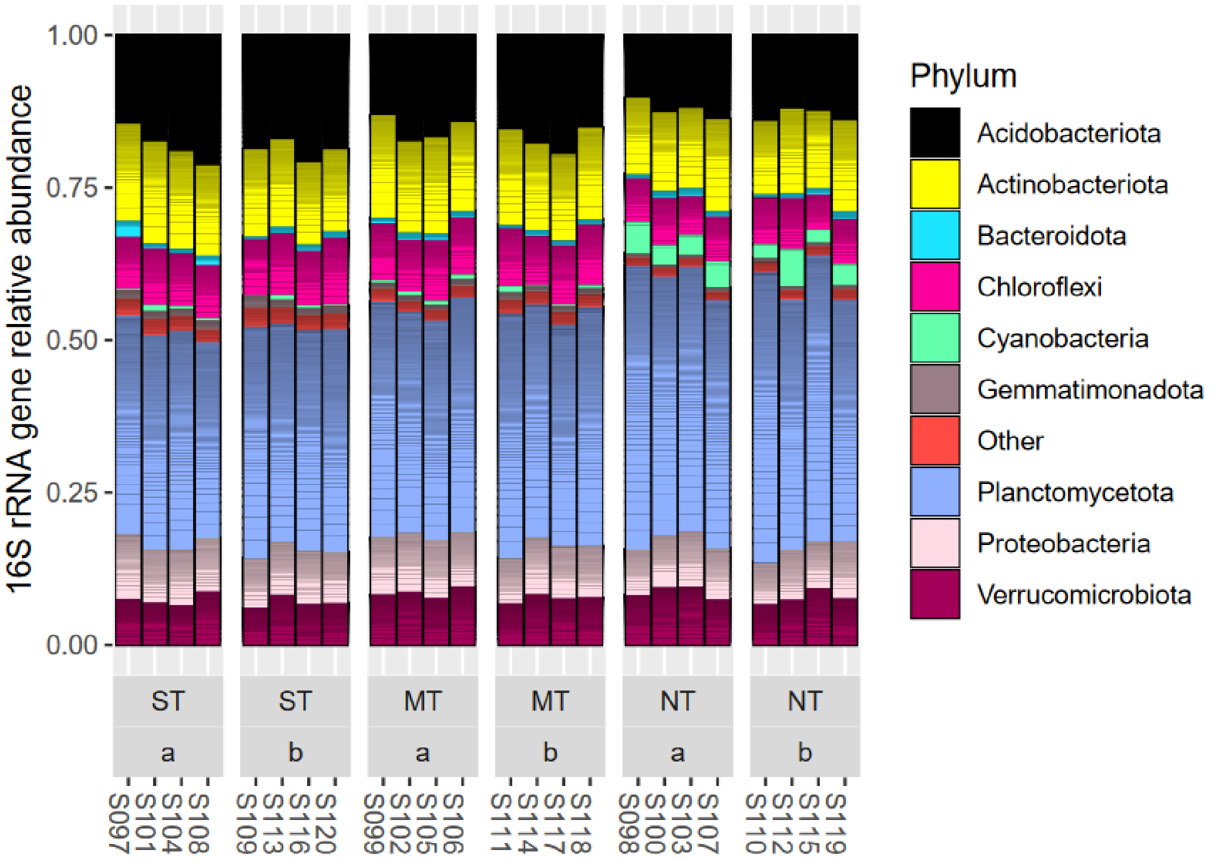


**Figure A1: Bacterial phyla relative abundances. Phyla with less than 1% relative abundance were bulked as other.**

##### I.I.I Proteobacteria – diverse and important biogeochemical cyclers

Shown in the following are the detected alpha and gamma classes of the phylum Proteobacteria. Missing in the dataset are classes, such as beta-proteobacteria, while Myxoccoci in the delta group have been reclassed to their own phylum.

Diverse metabolic functions in the transformation of C, N, P, S, Fe, H- compounds such as methylotrophy, sulfate reduction, nitrogen fixation, sulfur, iron, hydrogen and ammonia oxidation are usually found across Proteobacterial classes, as well as diverse chemoheterotrophic/ organic matter degradation capacities with specific genera (Madigan et al., 2021).

Alpha-Proteobacteria total relative abundance was higher in MT by ca. 1%, while the richness was similar in all tillage treatments.

In this class, 4 taxa with genus assignment exhibited significant associations to the tillage factor:

- Microvirga (family Beijerinckiaceae, order Rhizobiales, NT vs ST in both depths increase, Cluster 3)
- Rhodoplanes (family Xanthobacteraceae, order Rhizobiales, NT vs ST topsoil decrease, Cluster 1)

- Two Sphingomonas (family Sphingomonadaceae, order Sphingomonadales, NT vs ST lower soil decrease, Cluster 1)

The *Microvirga* OTU (family *Beijerinckiaceae*, order Rhizobiales) was also the taxon with the highest relative abundance of all treatments in this grouping. Members of this genus have been isolated from root nodules of Lupine (Ardley et al., 2012), and from rice field and forest soil (Zhang et al., 2009, Zhang et al., 2019).

Another taxon from the order Rhizobiales, genus *Rhodoplanes*, was also found to be significantly associated to the NT factor (decrease vs ST) in top soil layer.

The second and third most relatively abundant taxa in this group were two *Sphingomonas* OTUs (family *Sphingomonadaceae*). The first of these was significantly decreased in depth b of NT when compared to ST (significant LinDA test at FDR 5%), while the other was not. Another taxon from S. with lower relative abundance also was also significantly changing. This genus is characterized by obligately aerobic and nutritionally versatile organisms, that are generally easily cultivated. They notably exhibit the ability to degrade a wide range of organic compounds, especially aromatics such as environmental contaminants (Madigan et al., 2021). *Sphingomonas* have been investigated as control organisms against powdery mildew and Fusarium headblight in winter wheat (Wachowska et al., 2013).

In the Gamma-Proteobacteria, three taxa exhibited significant associations to the tillage management factor. Two were significantly increased in NT vs ST:

- Unassigned genus (“SC-I-84”) from order Burkholderiales (NT vs ST increased depth lower, Cluster 3)
- Unassigned genus from family *Rhodanobacteraceae* (order Xanthomonadales, NT vs ST increased both depths, Cluster 3)

One taxon was significantly decreased in NT vs ST in top layer:

- Unassigned from family *Steroidobacteraceae* (order Steroidobacterales, Cluster 1)

One taxon from the Burkholderiales was found to be significantly increased under NT tillage management in lower depth. The Burkholderiales order contains organisms with wide ranging metabolic and ecological properties, but also contain plant and human pathogens. In contrast to the classification with Silva database into Gamma-Proteobacteria (data here), references assign this order to the Beta-Proteobacteria (Madigan et al., 2021, Parte et al., 2020). Further genera for the order Burkholderiales were found in the Spanish soil such as *Massilia* (family *Oxalobacteraceae*, order Burkholderiales), which was the most relatively abundant, particularly in ST and MT management in the top layer (ns. different between tillage management). For example, a representative from this genus was isolated from soil under previous *Brassica* cultivation (Singh et al., 2015). Further notable taxa, in this order were *Variovorax* and other taxa from its family, the *Comamonadaceae*. Members of the family *Comamonadaceae,* are *Variovorax*, *Rhodoferax* and *Nitrosospira*. The latter, *Nitrosospira*, are typically capable of aerobic ammonia oxidation (first step of nitrification) for energy generation (lithotroph) (Madigan at el., 2021). *Variovorax* is a genus with organisms of aerobic chemoheterotrophic nutrition and notably degrade a wide range of substrates. Some strains are also capably of lithoautotrophic growth with hydrogen as energy source (Willems et al., 1991).

A taxon from the family *Rhodanobacteraceae*, order Xanthomonadales, was found to have significant association to the treatment factor (increase NT versus ST at both depths, $p<5\%$).

The family consists of the genera *Aquimonas, Dokdonella, Dyella, Frateuria, Fulvimonas, Luteibacter, Pseudofulvimonas, Rhodanobacter* (the type genus for this family) and *Rudaea*. These are non-spore forming aerobic chemoorganotrophs (Naushad et al., 2015). The family is a sister group to the *Xanthomonadaceae* (*Lysobacteraceae*). A taxon from the *Lysobacter* (fam. *Xanthomonadaceae)* genus was also found in the soil extracts. *Lysobacter* were reported to degrade polysaccharides (i.e. chitin) and a wide range of proteinaceous compounds, which includes predating on other microorganisms such as bacteria, algae, yeasts, and filamentous fungi (Christensen, 2015).

Another taxon with significant association to NT vs ST (decrease) was from the *Steroidobacteraceae* family. The two genera, *Steroidobacter* and *Povalibacter*, were moved to this novel family following polyphasic analysis. Members exhibit chemoorganotrophic nutrition and some have ability for assimilatory and dissimilatory nitrate reduction ammonification (ANRA / DNRA) (Liu et al., 2019).

Further notable taxa in the dataset, but without significant effects from tillage management, are *Pseudomonas*. This group contains fermenters, well-known plant and human pathogens and exhibits the ability to form biofilms (Madigan et al., 2021). Some of these organisms are also capable of anaerobic dissimilatory nitrate reduction to $N_2O$ (denitrification) in anoxic conditions (for example water logged soil).

The predatory Myxobacteria have been reclassified to their own phylum Myxococcota (presented later).

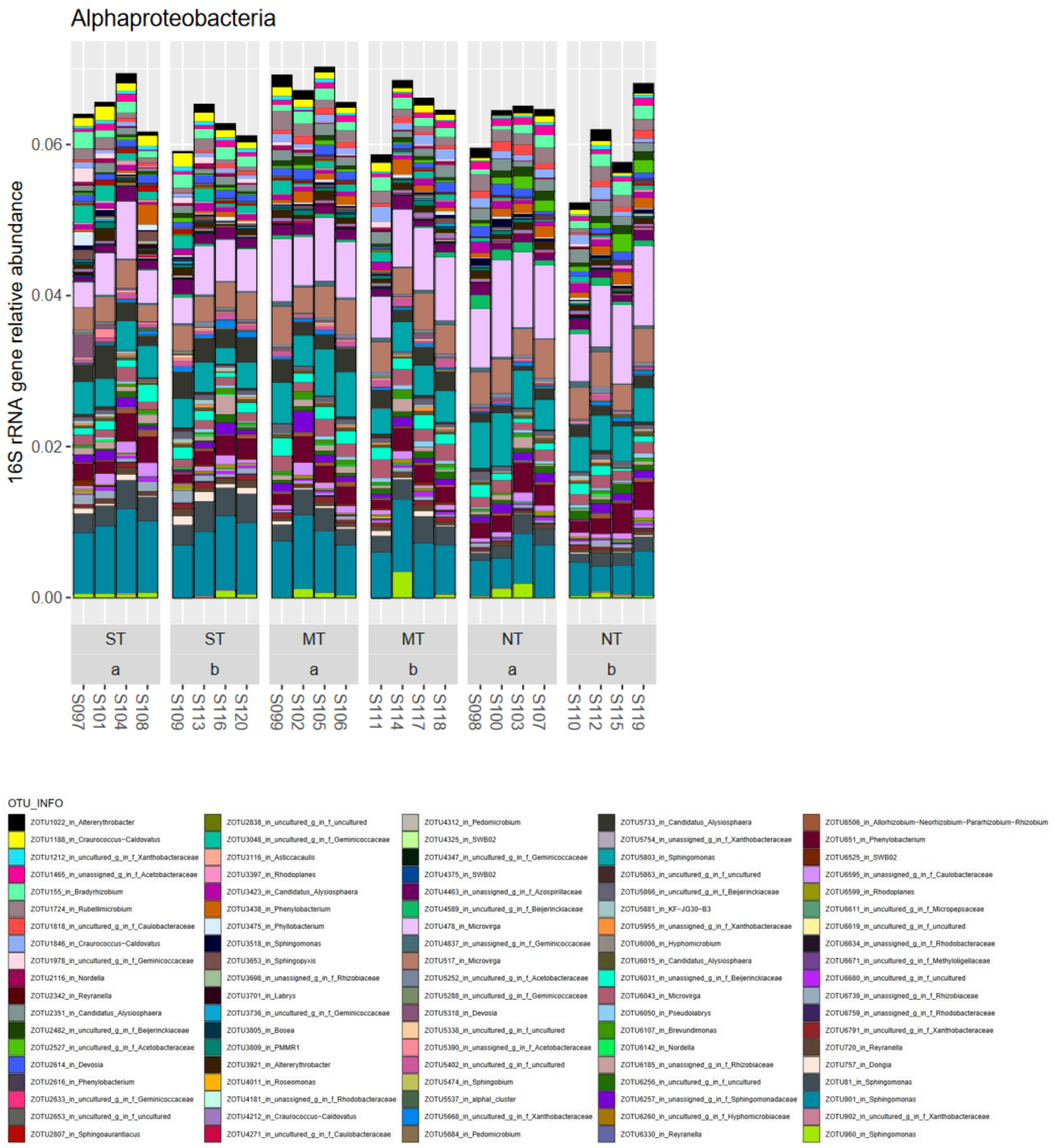


**Figure A2: Alpha-Proteobacteria relative abundances.**

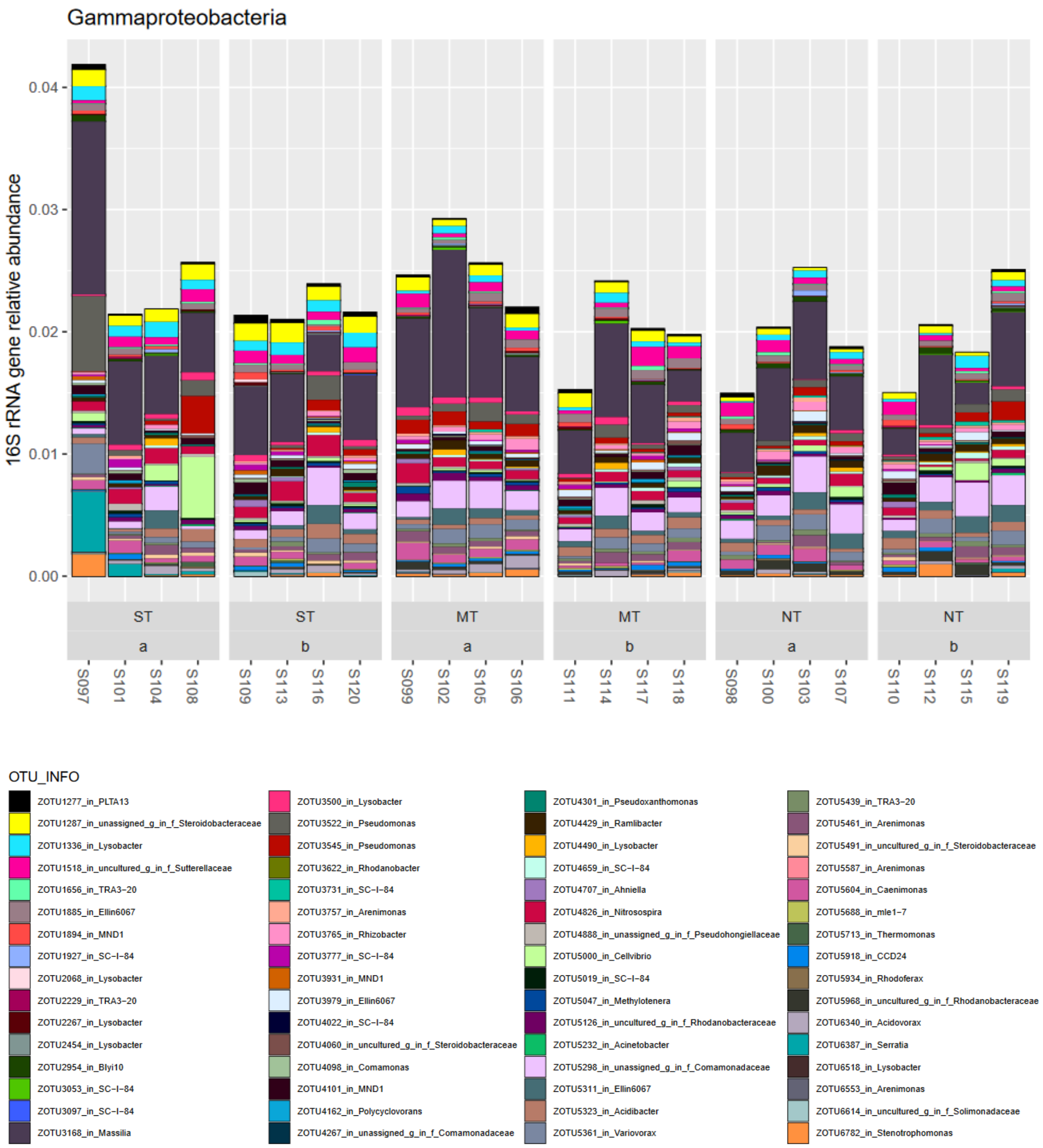


**Figure A3: Gamma-Proteobacteria relative abundances.**

##### 1.1.2 Actinobacteria – able competitors

The phylum Actinobacteriota contained the following classes in descending order of relative abundance: Actinobacteria, Thermoleophilia, Rubrobacteria, Acidimicrobiia, and MB−A2−108.

Actinobacterial taxa with significant associations to tillage treatment factors were found in the families *Micromonosporaceae*, *Micrococcaceae*, *Ilumatobacteraceae*, *Cellulomonadaceae*, *Streptomycetaceae*, and *Gaiellaceae* (LinDA, FDR<5%).

In the class Actinobacteria, four taxa from the family *Micromonosporaceae* (order Micromonosporales) were significantly changing. Of those, three were relatively increasing under NT vs ST (both depths, Clusters 3 and 6; **Cluster 6 has increases for taxa in NT and MT!**). One of them, *Actinoplanes*, (Cluster 6) was assigned on genus level. *Micromonosporaceae* family contains organisms often isolated from soil and specifically with some isolates from the rhizosphere of e.g. lupines and peas. Members of this family are noted for their ability to produce specialized metabolites (i.e. antibiotics), polymeric organic matter degrading enzymes, biocontrol, and plant growth hormones (Carro et al., 2018). The characteristics of this family are that they are aerobic, Gram-positive, mesophilic organisms that form mycelia (Nouinoui et al., 2018). Another taxon with genus level assignment was *Luedemanella.* It was significantly decreased in NT, with MT and ST similar in relative increase (LinDA, FDR<5%). This genus also belongs to the family *Micromonosporaceae* (Ara and Kudo, 2007).

Further taxa with significant effects in the class Actinobacteria were from *Streptomyces* (decrease in NT vs ST both depths, cluster 2) from the order Streptomycetales, and further taxa such as *Cellulomonas* (increase NT vs ST both depths, cluster 3) and one unassigned from the order Micrococcales (families *Cellulomonodaceae* and *Micrococcaceae*). The unassigned genus taxon was a highly relatively abundant taxon from the family *Micrococcaceae* (zOTU122, cluster 1) with significant decreases in NT vs ST in top soil.

*Streptomyces* are well known for their filamentous growth, capacity to degrade polymers such as cellulose and hemicellulose, and antibiotics synthesis (Madigan et al., 2021). Some Streptomyces strains have been investigated as biocontrol agents against the cereal head blight *Fusarium graminearum* (Jung et al., 2013).

Most strains of *Cellulomonas*, the only type genus of the *Cellulomonadaceae* family, are cellulolytic (Nouinoui et al., 2018). The family *Micrococcaceae* contains the type genus *Micrococcus*, and further well-known organic matter degraders such as for examples *Arthrobacter*. Further genera in this family are *Acaricomes*, *Citricoccus*, *Kocuria*, *Nesterenkonia*, *Renibacterium*, and *Rothia* (Zhi et al., 2009).

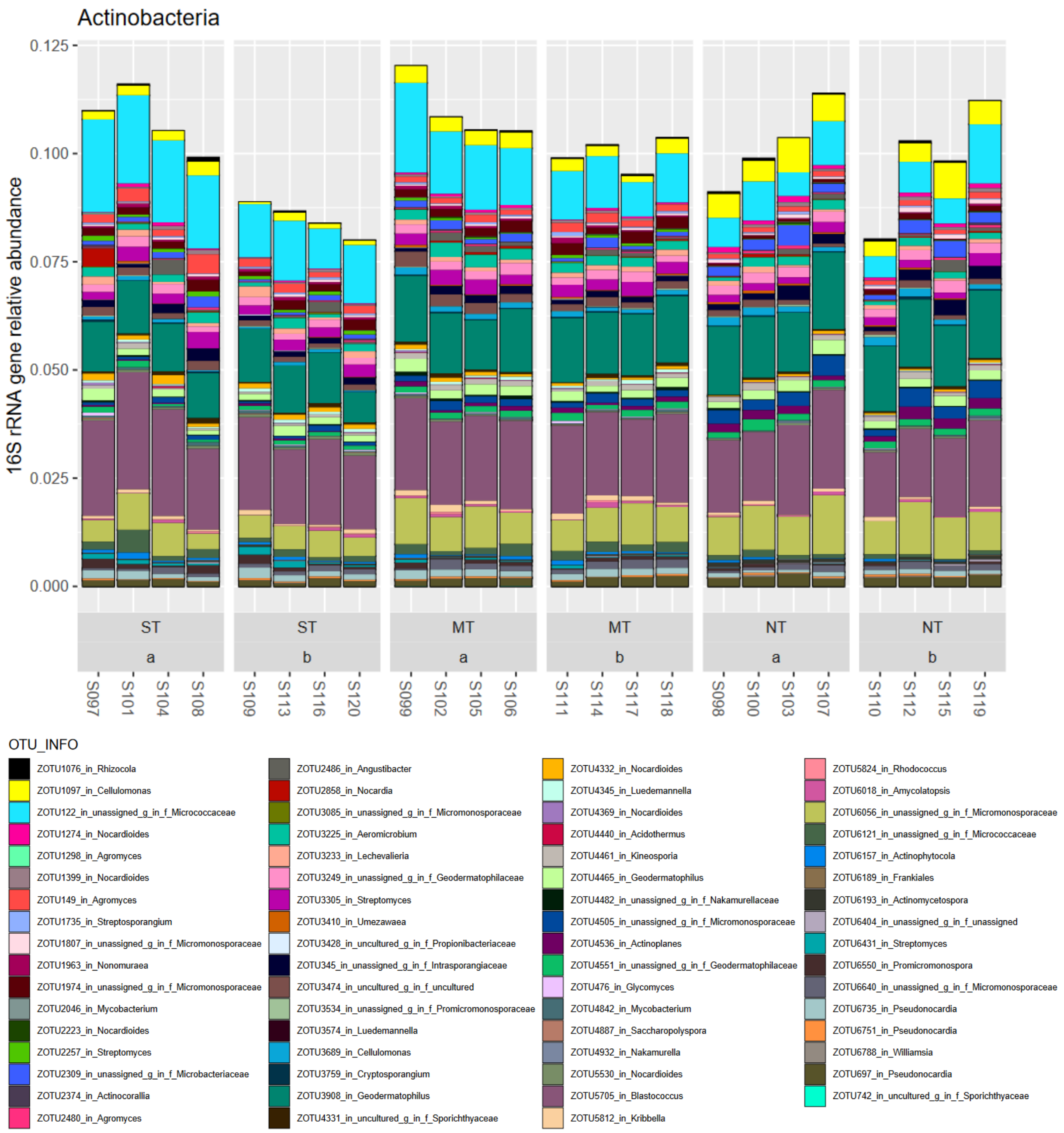


**Figure A4: Class Actinobacteria in the phylum Actinobacteria relative abundances.**

In the class Thermoleophilia, significant changes with tillage management were detected for 6 taxa in the orders Gaiellales and Solirubrobacteriales. All these taxa were found to be significantly decreased in NT vs ST (top soil and/or lower soil, clusters 1, 2, 4).

In the results, the only genus assignment was to *Gaiella* in family *Gaiellaceae*. The only representative and type strain (*Gaiella occulta*) defines the classification up to its own order, Gaiellales, and has been characterized as strictly aerobic, degrading gelatine but not assimilating simple sugars, amino acids and aromatic hydrocarbons, and being able to reduce nitrate to nitrite. Members of the order Gaiellales and the family *Gailleaceae* are Gram-negative, strictly aerobic chemoorganotrophs and non-pigmented. Members of the Solirubrobacterales are Gram-positive and some are pigmented (Albuquerque et al., 2011).

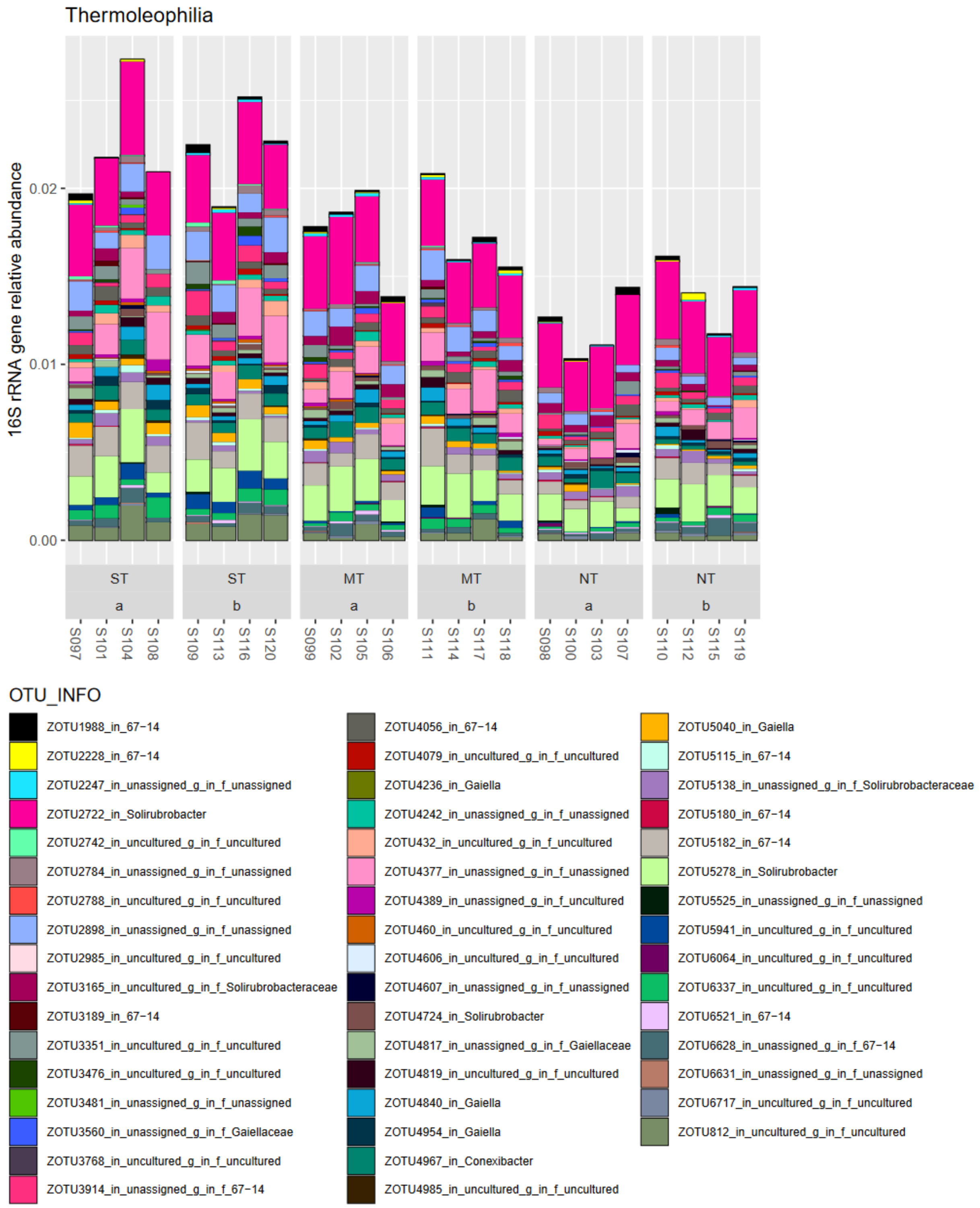


**Figure A5: Class Thermoleophilia in the phylum Actinobacteria relative abundance.**

In the class Rubrobacteria, no significant changes were detected with tillage management factor. The strains in this genus are aerobic organotrophs, with ability to reduce nitrate to nitrite. Some strains exhibit desiccation and/or radiation tolerance (Albuquerque et al., 2014).

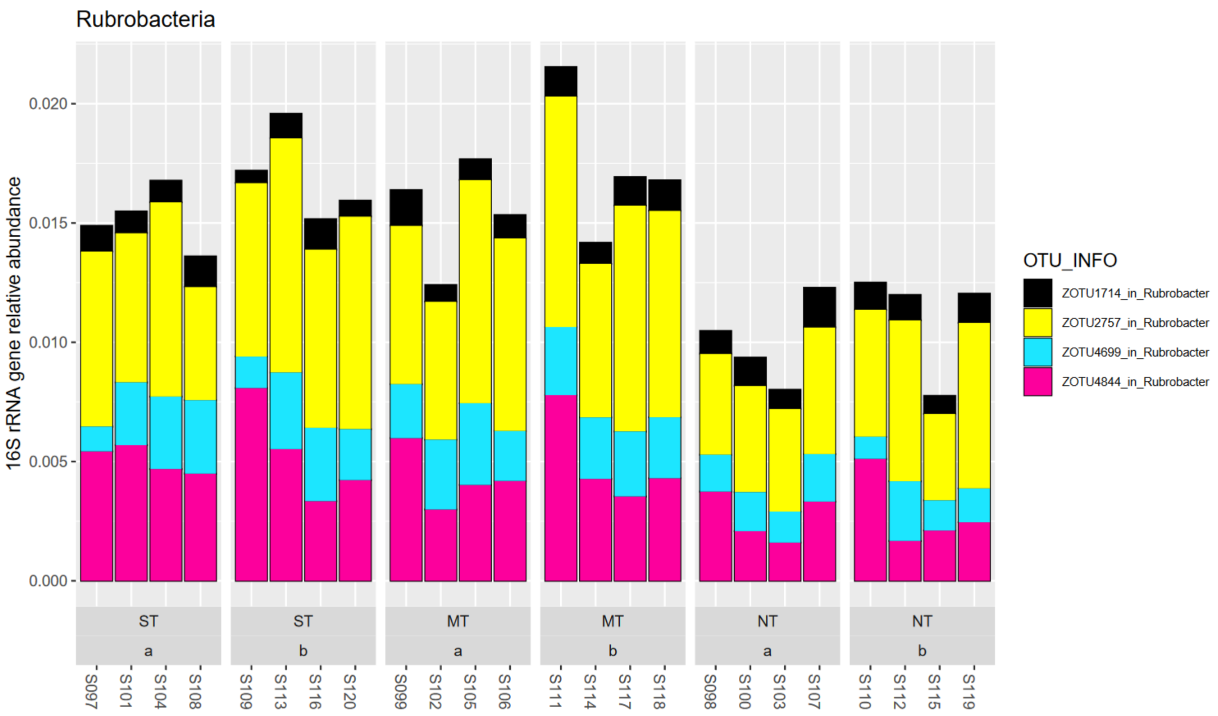


**Figure A6: Order Rubrobacteria in phylum Actinobacteria relative abundance.**

In the class Acidimicrobiia, a taxon from the genus *Illumatobacter* (family *Ilumatobacteraceae*) was significantly increased under NT versus ST in lower soil depth (10-20 cm). Several strains of this genus have been isolated from sediment. They were Gram-positive and aerobic growing, and exhibited alkaline and acid phosphatase, esterase, lipase, leucine arylamidase, and naphtol-AS-BI-phosphohydrolase but not several activities on simple sugars such as for example a-glucosidase (Matsumoto et al., 2013, 2009).

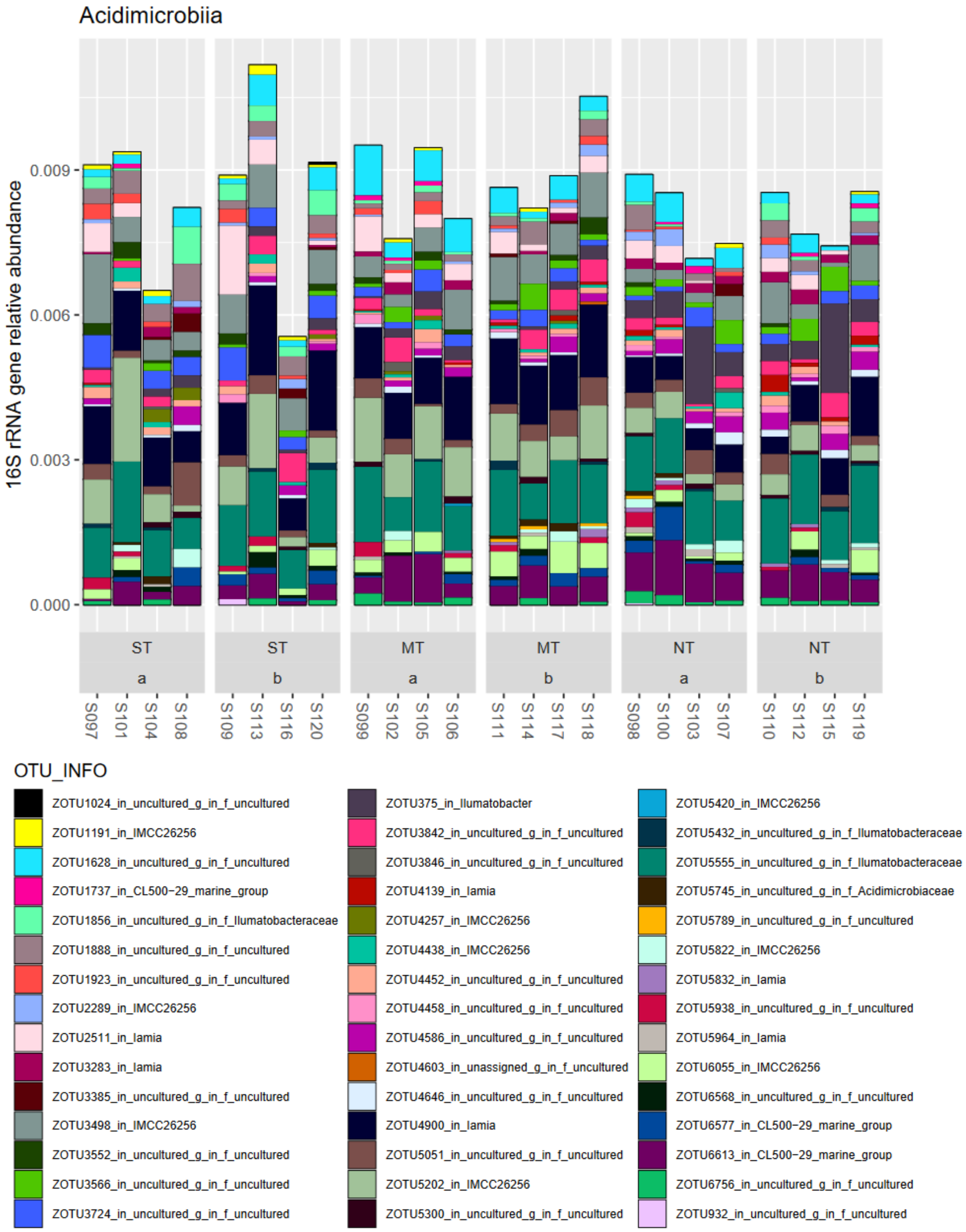


**Figure A7: Class Acidimicrobiia in phylum Actinobacteria**

Four taxa in the class MB-A2-108 exhibited significant decreases with NT management (Clusters 1, 2, 5). None have organisms with order, family, or genus level assignments (no isolate so far?). Particularly cluster 5 was characterized by taxa with the strongest decreases in NT management.

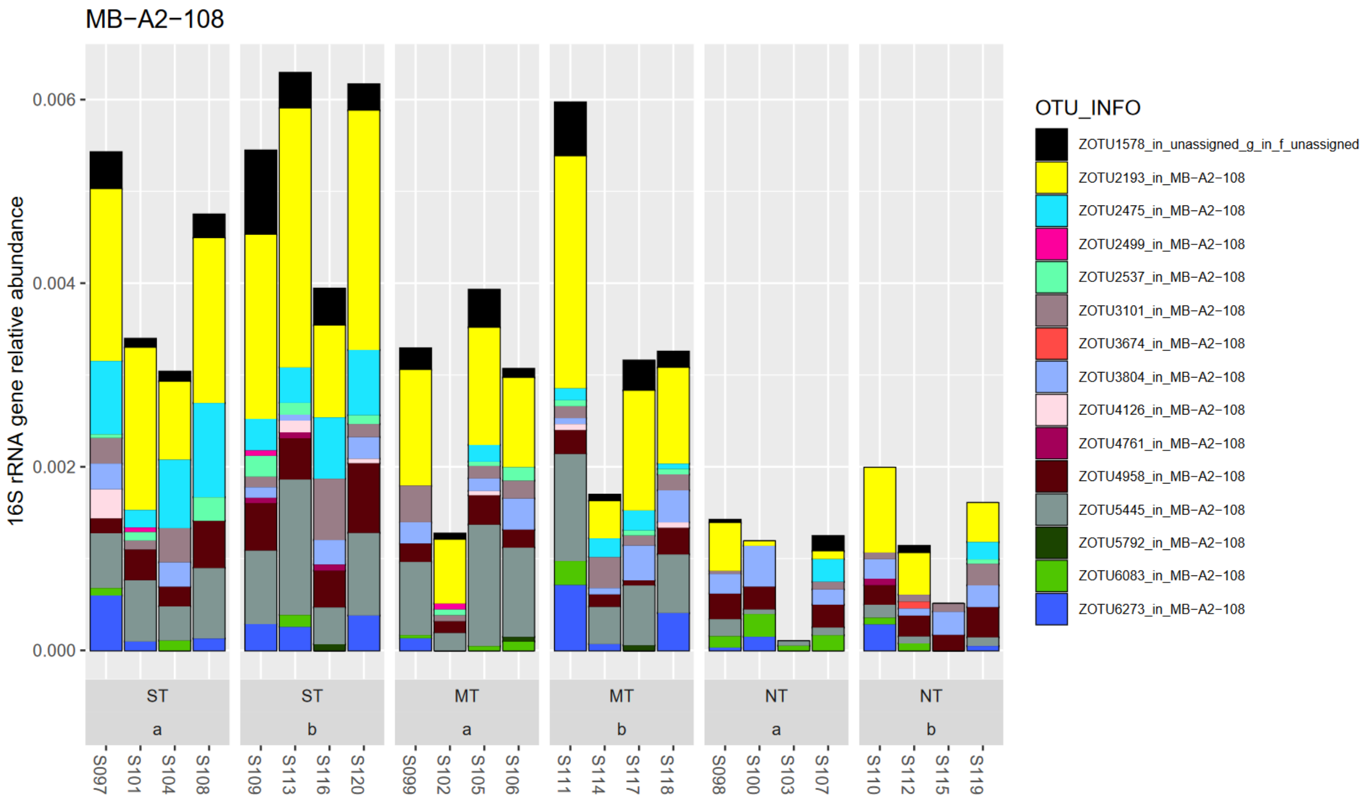


**Figure A8: Class MB-A2-108 in the phylum Actinobacteria**

##### 1.1.3 Firmicutes – some well-known taxa such as *Bacillus*

Two taxa with significant increases in ST management vs NT and MT (Cluster 4) were detected in the phylum Firmicutes, genera *Bacillus* (OTU 5532) and *Cohnella* (OTU 4886). These two taxa were of comparably lower relative abundance in the dataset of Firmicutes. Generally, the total relative abundance of Firmicutes to the total bacterial community was ca. 0.4% and the richness consisted of only 10 taxa.

Bacilli are usually a rich phylum of organisms with well-known relevance for soil-plant-fauna interactions in the Bacillales order such as *Bacilli, Paenibacillus, Sporosarcina,* and *Staphylococcus* (Madigan et al., 2021). The genus *Bacillus* with over 280 reported species is considered polyphyletic with reclassifications suggesting that only those belonging to *B.cereus* and *B.subtilis* clade remain and others be assigned to 17 novel genera of *Bacillaceae* (Gupta et al., 2020). *Cohnella* organisms were isolated from diverse environments such as hot substrates, water, root nodules and soils of forest, wetland and dryland. The organisms are either aerobic or facultatively anaerobic and stain Gram positive or negative, with the type strain growing at high temperatures/thermotolerant. A most recent isolate from soil exhibited xylanolytic activity (Khianngam et al., 2010).

In this dataset, the relative abundance of 16S gene amplicons from Firmicutes was mostly below 0.8%. The low relative abundance on DNA level of soil extracts was also found in other studies in Mediterranean soil (Dal Cortivo et al., 2020, Illescas et al., 2020, Visioli et al., 2020).

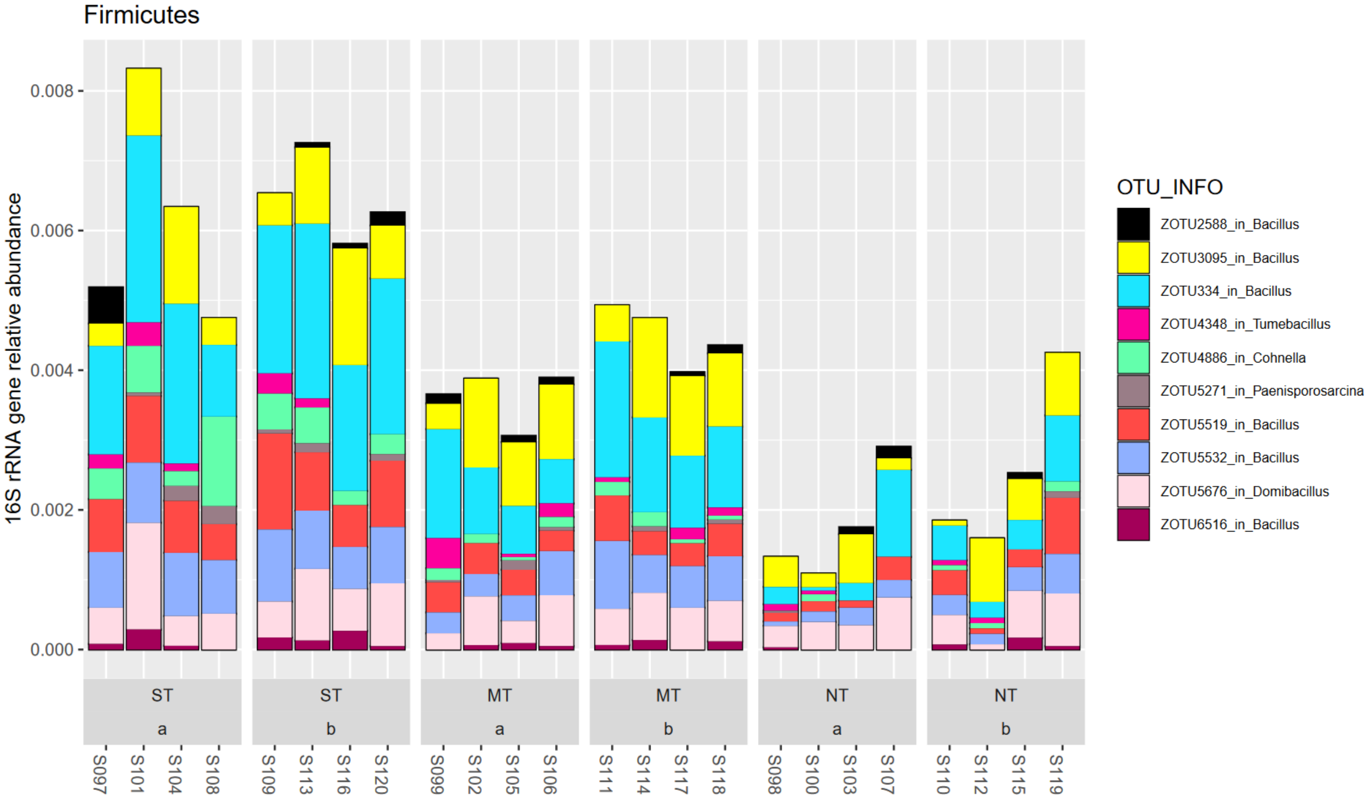


**Figure A9: Phylum Firmicutes. Comparably low relative abundance and richness of genera found in this dataset.**

##### 1.1.4 Acidobacteria – increasingly recognized but difficult to culture

In the dataset, the Acidobacteria constituted the second most relatively abundant phylum of 16S rRNA gene copies (15.8% of total of 16S amplicon set). In the dataset, the most relatively abundant and richest classes were constituted by the Vicinamibacteria, followed by Blastocatellia, and Acidobacteriae. Further classes are also shown below. With respect to the tillage treatment factor, 25 taxa were significantly associated (LinDA, FDR<5%).

While the Acidobacteria are widespread in the environment, few have been cultured to date, with few information on nutrition and lifestyle (Madigan et al., 2021). Increasingly, strains are successfully isolated and described, and circa 60 species in 26 classes have been described (Huber and Overmann, 2018).

The *Vicinamibacteria* class contained 13 taxa with significant changes to tillage treatments (all in cluster 1,2,4,5 which are characterized by decreases in NT vs ST – or termed *vice versa* increases in ST vs NT). Of those, 6 taxa were from the *Vicinamibacteraceae* family. The family is characterized by strains with aerobic chemo-organo-heterotrophic nutrition on simple sugars, and complex proteinaceous compounds and with enzyme activities for alkaline and acid phosphatase, naphtol-AS-BI-phosphohydrolase, α-chymotrypsin, trypsin and esterase C4 (Huber and Overmann, 2018).

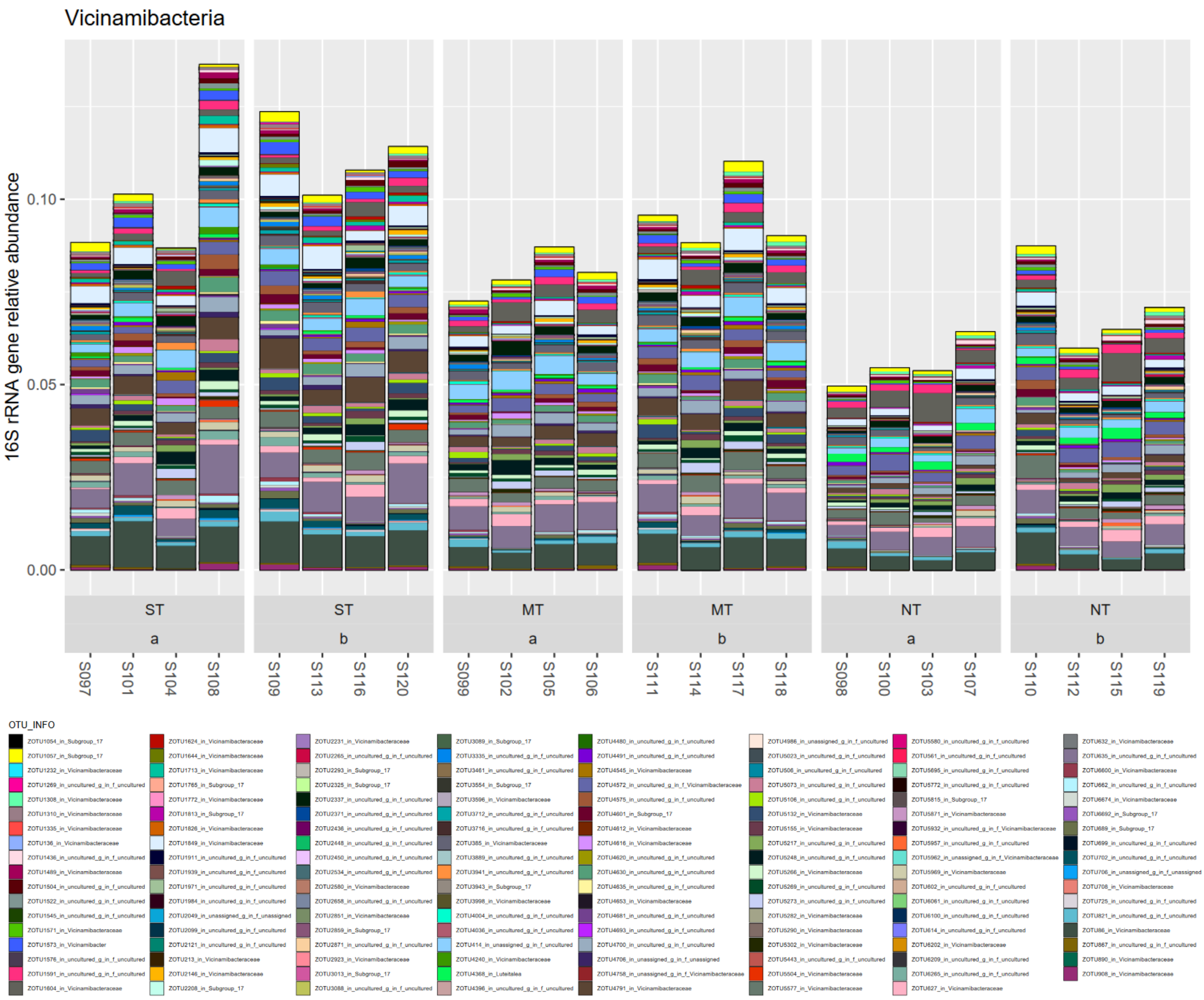


**Figure A10: Class Vicinamibacteria in phylum Acidobacteria.**

The class Blastocatellia exhibited 9 taxa with significant associations to the tillage treatment (of those 4 were in clusters with increases in NT vs ST, the rest in decrease clusters), of which 4 are from the *Blastocatellaceae* family (all these increased under NT). Interestingly, some taxa from the family *Pyrinomonadaceae* (taxa 6258, 72, 3751, 1717) and with the genus code RB41 were indicated to be significantly affected by tillage management (decrease under NT), but taxa with much higher relative abundance in this genus were not significantly changing.

The class contains on the one hand moderately thermophilic, and slightly alkaliphilic aerobic chemoheterotrophic organisms and on the other hand microaerophilic and anoxygenic photoheterotrophs (Pascual et al., 2015).

Members of the *Blastocatellaceae* family have been characterized as K-strategists/oligotrophs/nutrient poor soils with a preference for complex proteinaceous substrates (Pascual et al., 2015).

Organisms from the *Pyrinomonadaceae*, are characterized by aerobic chemo-organo-heterotrophic but limited range of substrate utilization/nutrition. A preference for proteinaceous substrates and incidences of polymer hydrolyzation capacity were described (Wüst et al., 2016).

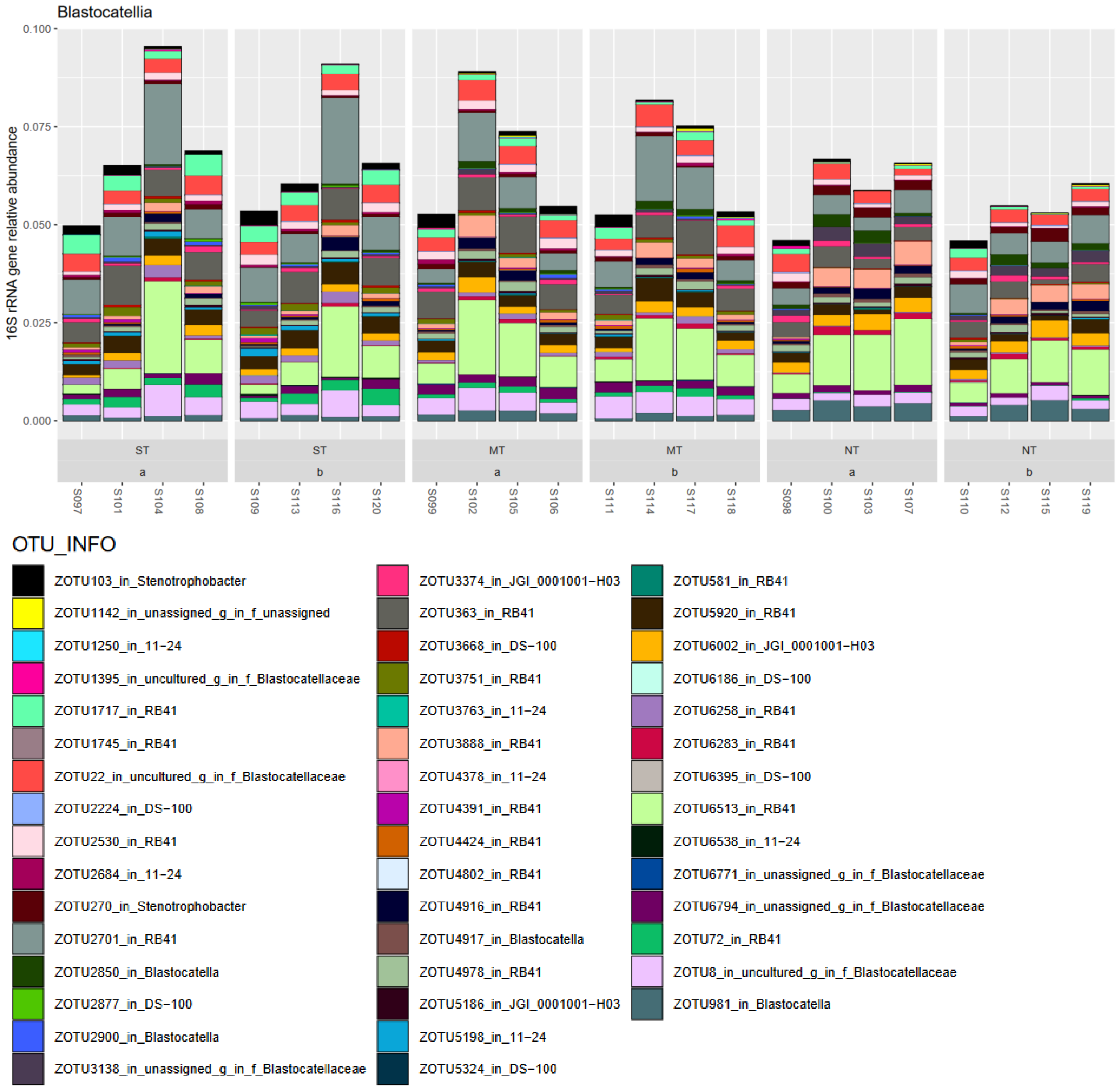


**Figure A11: Class Blastocatellia in phylum Acidobacteria.**

No members from the class Acidobacteriae were significantly associated with tillage treatment factors. A taxon from the genus *Bryobacter* was relatively prominent in all treatments. This genus is characterised by strictly aerobic chemo-organotrophic nutrition on sugars. The type strain was isolated from peat moss but does not utilise most organic acids, nitrate nor urea and is unable to hydrolyse polymers cellulose, xylan, cellulose, and chitin, among others (Kulichevskaya et al., 2010).

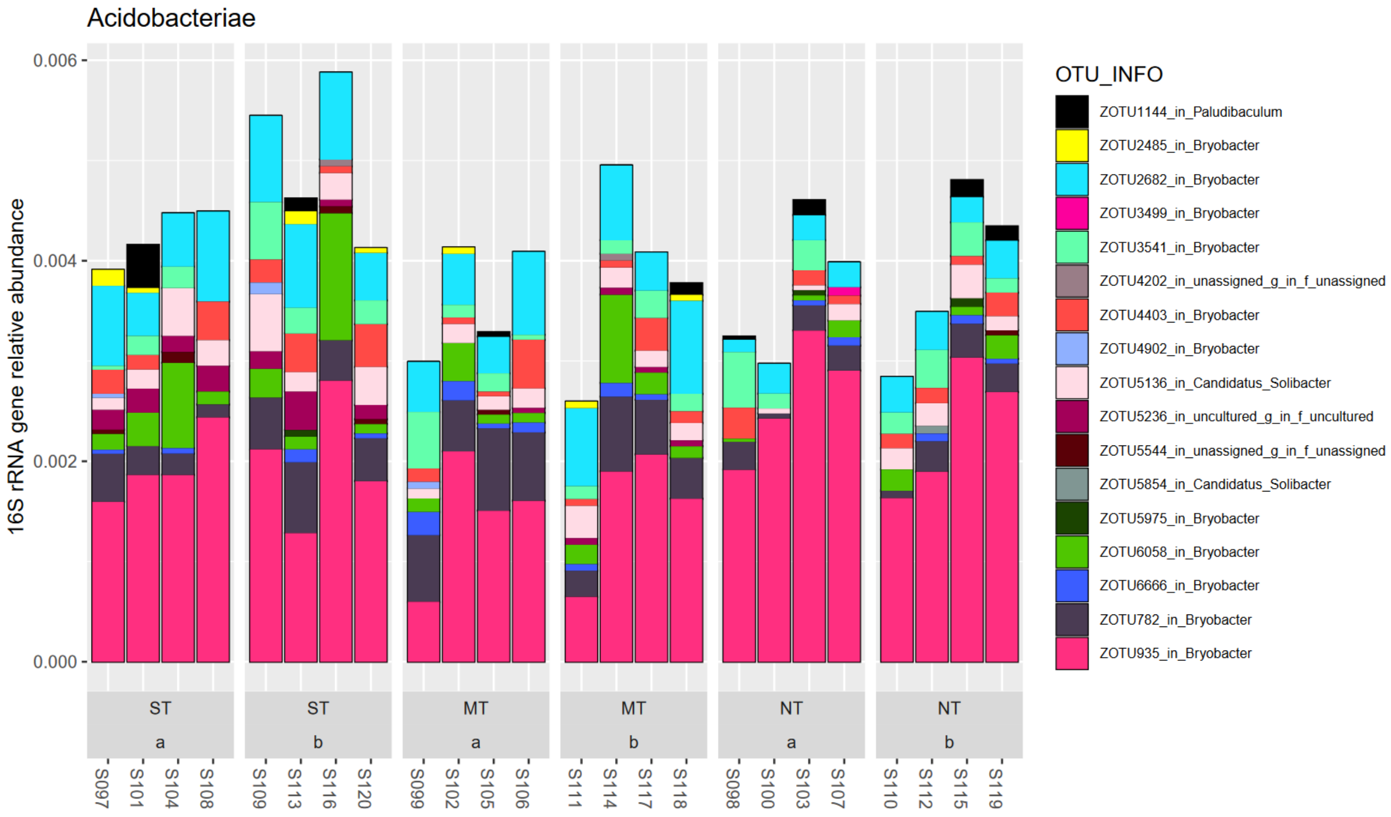


**Figure A12: Class Acidobacteriae in phylum Acidobacteria.**

The class Thermoanaerobaculia exhibited only one taxon with significant changes to the tillage treatment and with only low relative abundance (zOTU4085). Generally, this class is formed by one cultured representative and several environmental clone sequences from aquatic habitats (thermal hot springs in marine and terrestrial) (Dedysh and Yilmaz, 2018).

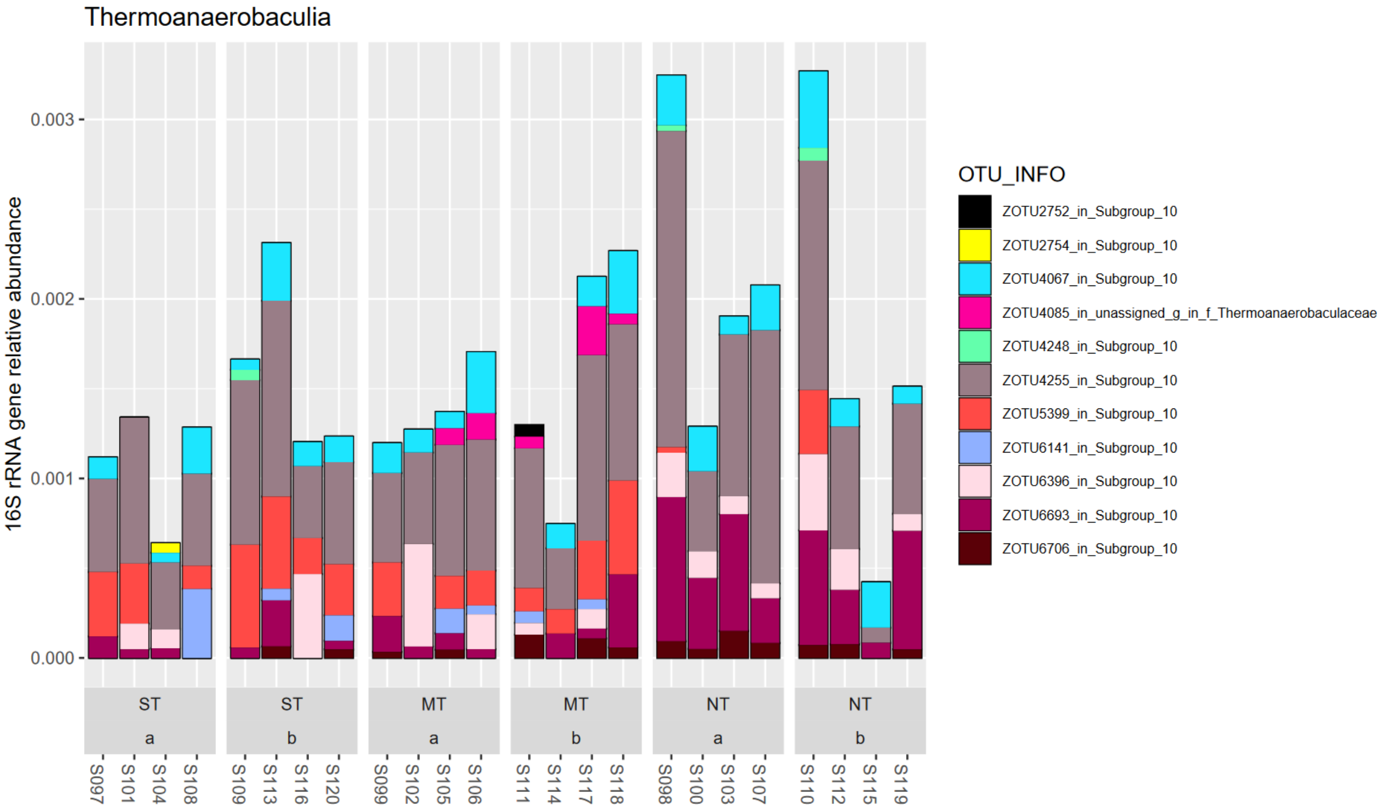


**Figure A13: Class Thermoanaerobaculia in phylum Acidobacteria.**

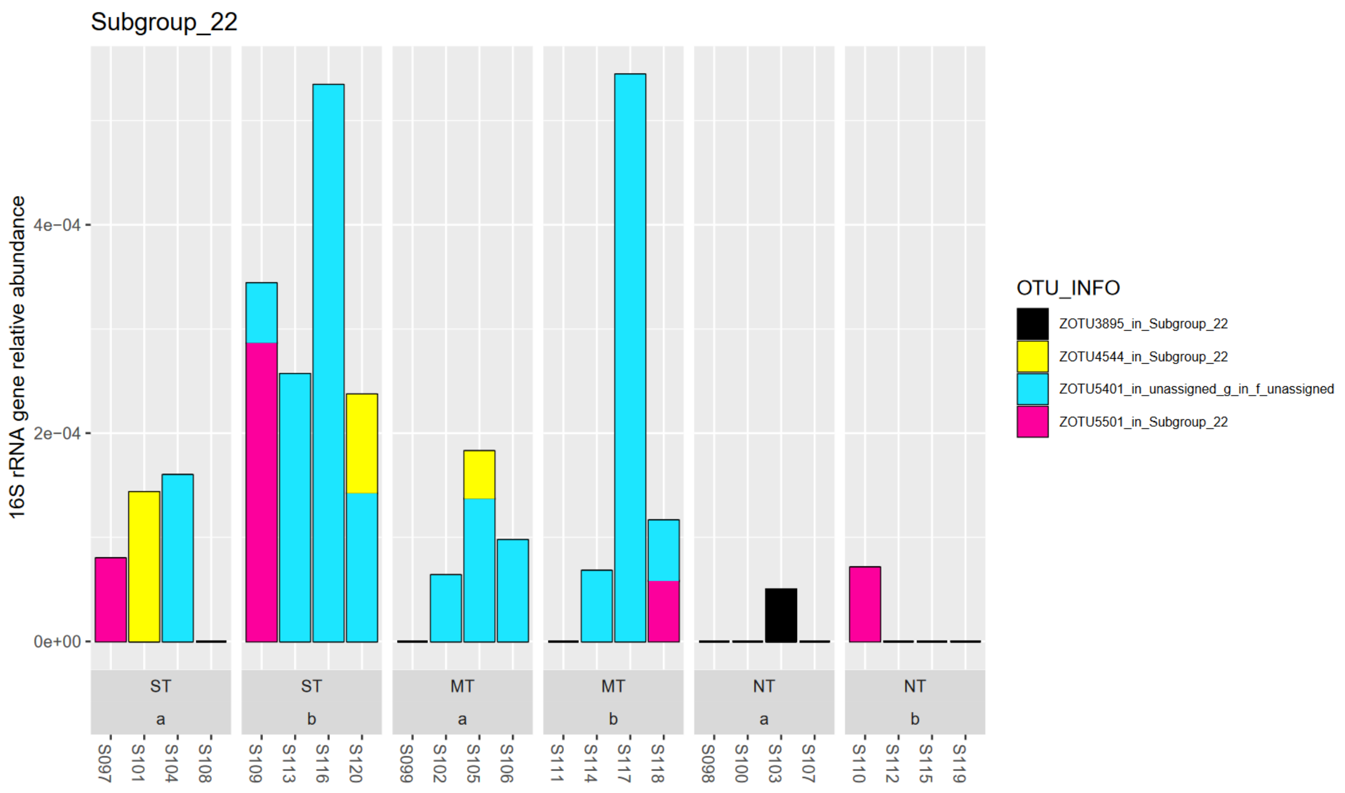


**Figure A14: Subgroup 22 in phylum Acidobacteria**

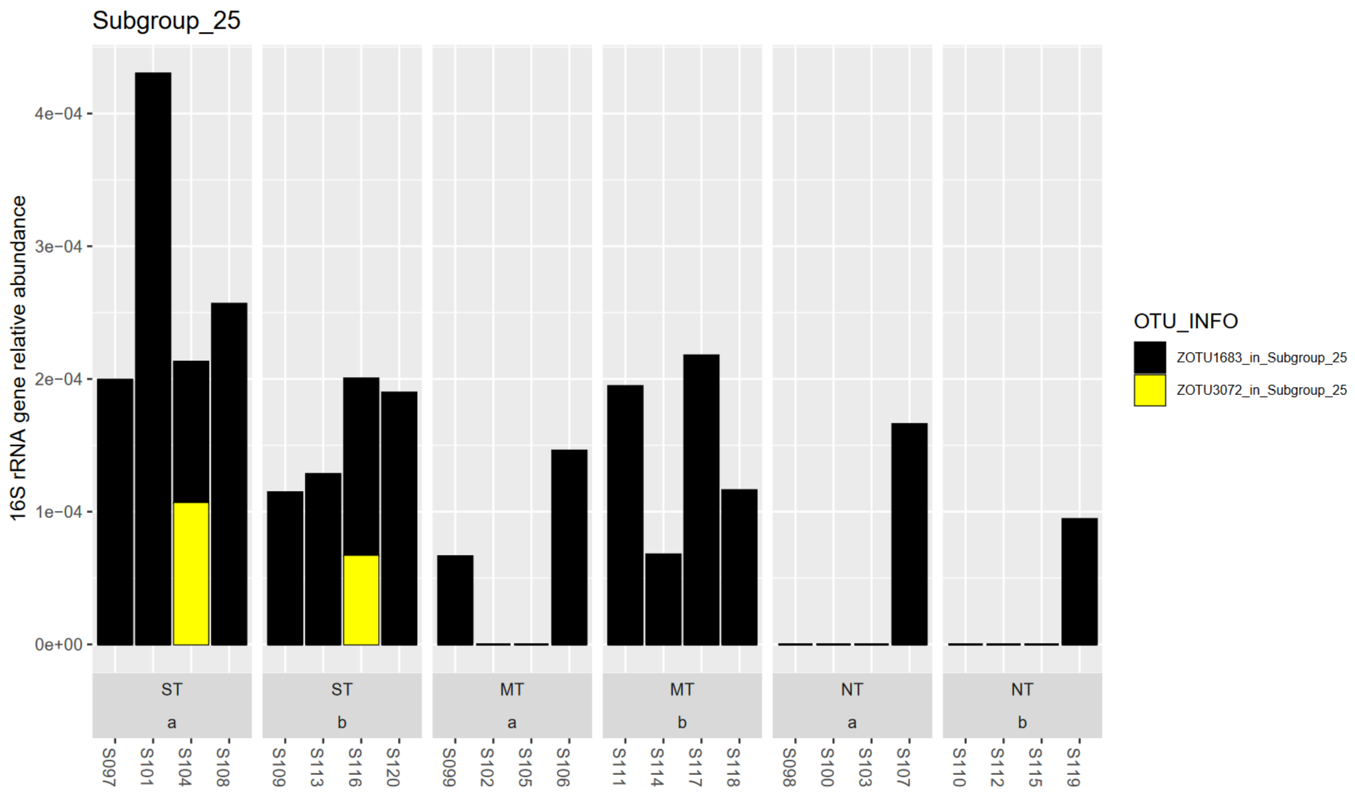


**Figure A15: Subgroup 25 in phylum Acidobacteria**

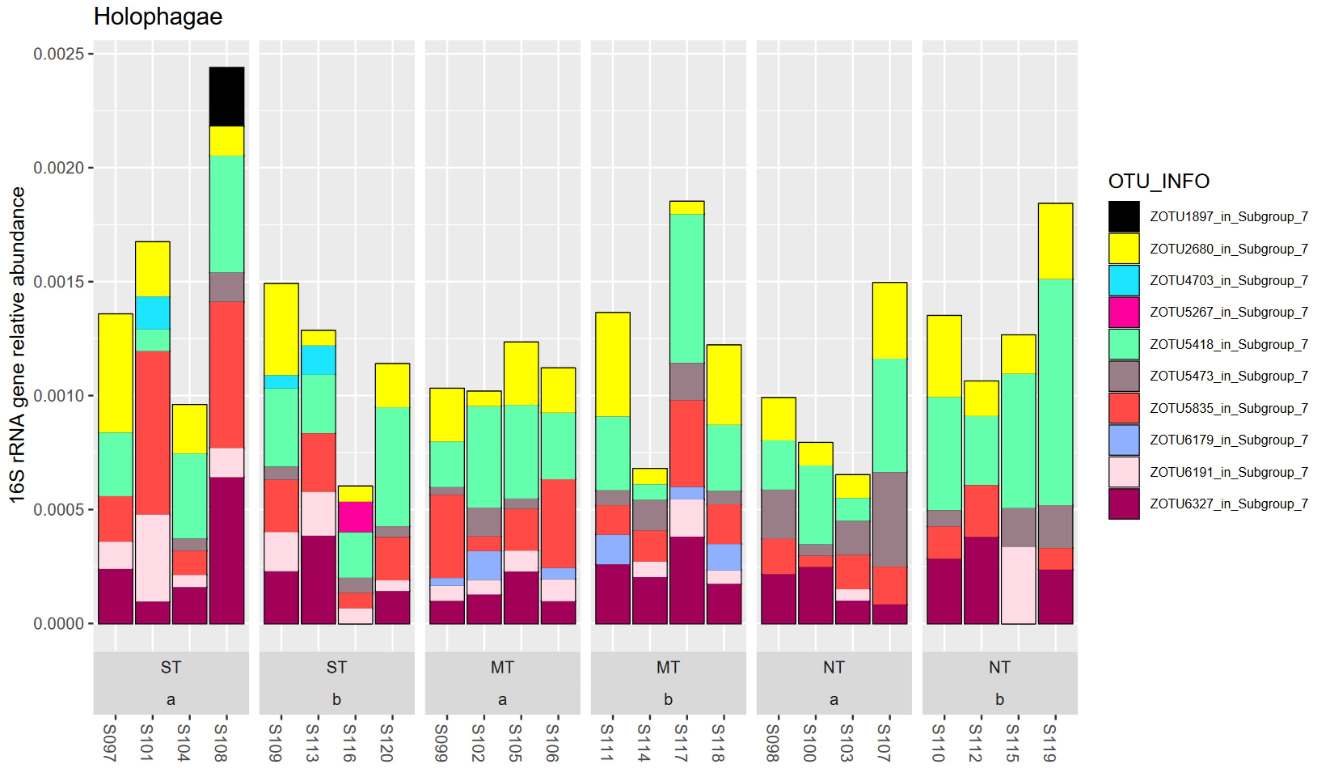


**Figure A16: Holophagaceae in phylum Acidobacteria**

##### 1.1.5 Predatory bacteria – important for regulation of prey populations

Typical predatory clades were found in the dataset from the Myxococcota and Bdellovibrionota, though predators are also found in the phyla Proteobacteria, Bacteroidetes, Chloroflexi and Melainabacteria (Madigan et al., 2021). Previously, Myxococcales were classed under the Delta-Proteobacteria, but polyphasic investigations have warranted a reclassification into four new phyla, Bdellovibrionota, Myxococcota, Desulfobacterota, and "SAR324", which is also consistent with different hunting strategy of their members (Waite et al., 2020). In the case of Bdellovibrionota, predators operate by infiltrating the periplasm of other bacteria, "periplasmic predators" and in the case of Myxococcota, organisms operate as a group swarming to search for prey, "social predators" (Madigan et al., 2021). Predators can in principle enable higher diversity and co-existence of species that would otherwise compete strongly for the same niche, viz. they can reduce abundant/exclude species that would otherwise outcompete other more inferior competitors (Putman and Wratten, 1988).

In the Spanish tillage trial, Bdellovibrionota exhibited no significant associations to the tillage factors (LinDA, FDR>5%).

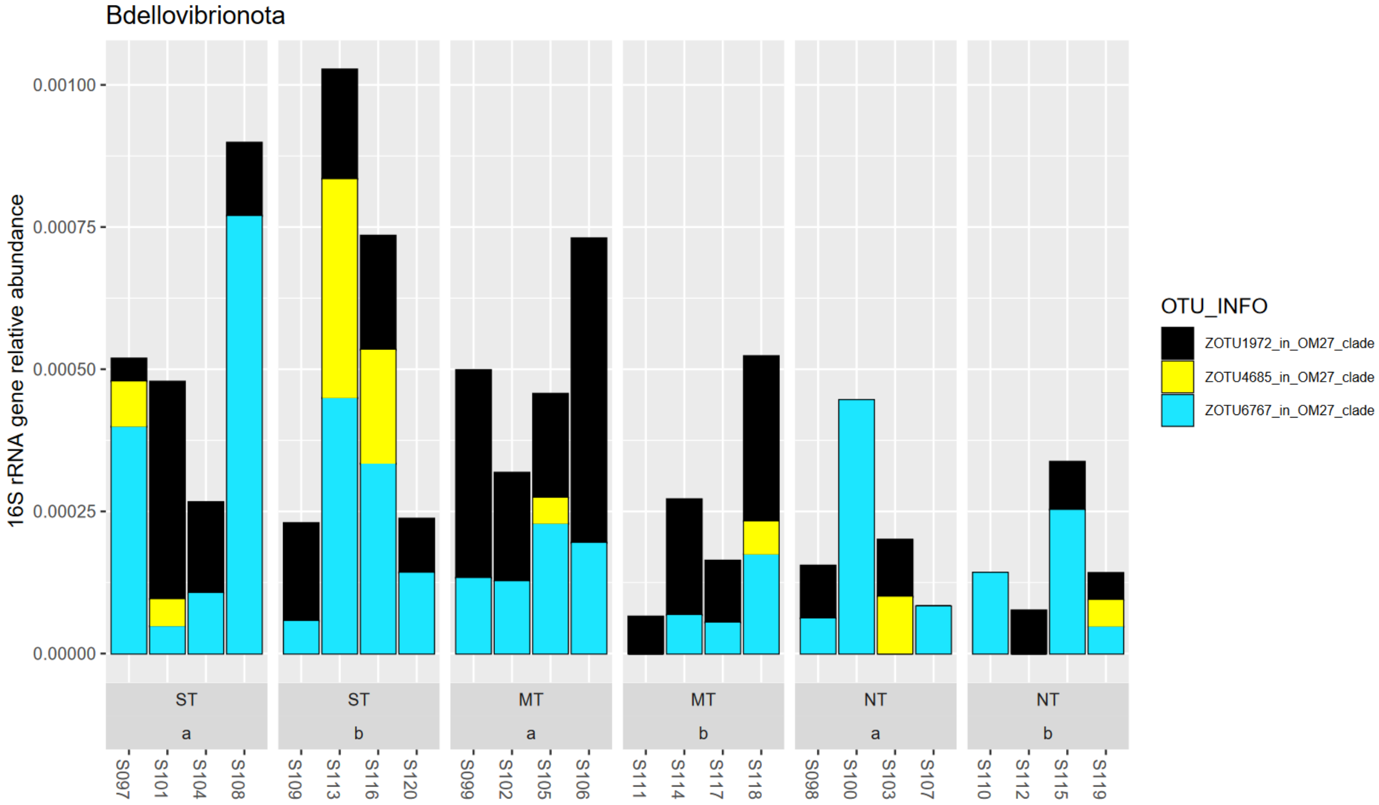


**Figure A17: Phylum Bdellovibrionota**

The Myxococcota constituted up to 1% relative abundance of the 16S amplicons. Most relatively abundant in this phylum was a taxon from the family *Myxococcaceae*, and in second place a taxon from the genus *Nannocystis (*family *Nannocystaceae)*. Only one taxon in this phylum, zOTU 3764 from the order Polyangiales, family *Blri41*, was found to be significantly increased in NT vs ST in lower soil depth, but it was of comparably lower relative abundance in this class. The order Polyangiales contains the families *Polyangiaceae* and *Sandaracinaceae*. The family *Polyangiaceae* contains the genera *Chondromyces, Sorangium, Minicystis, Labilithrix, Pajaroellobacter*, and *Phaselicistis*. The family *Sandaracinaceae* contains the genus *Sandaracinus* (Waite et al., 2020).

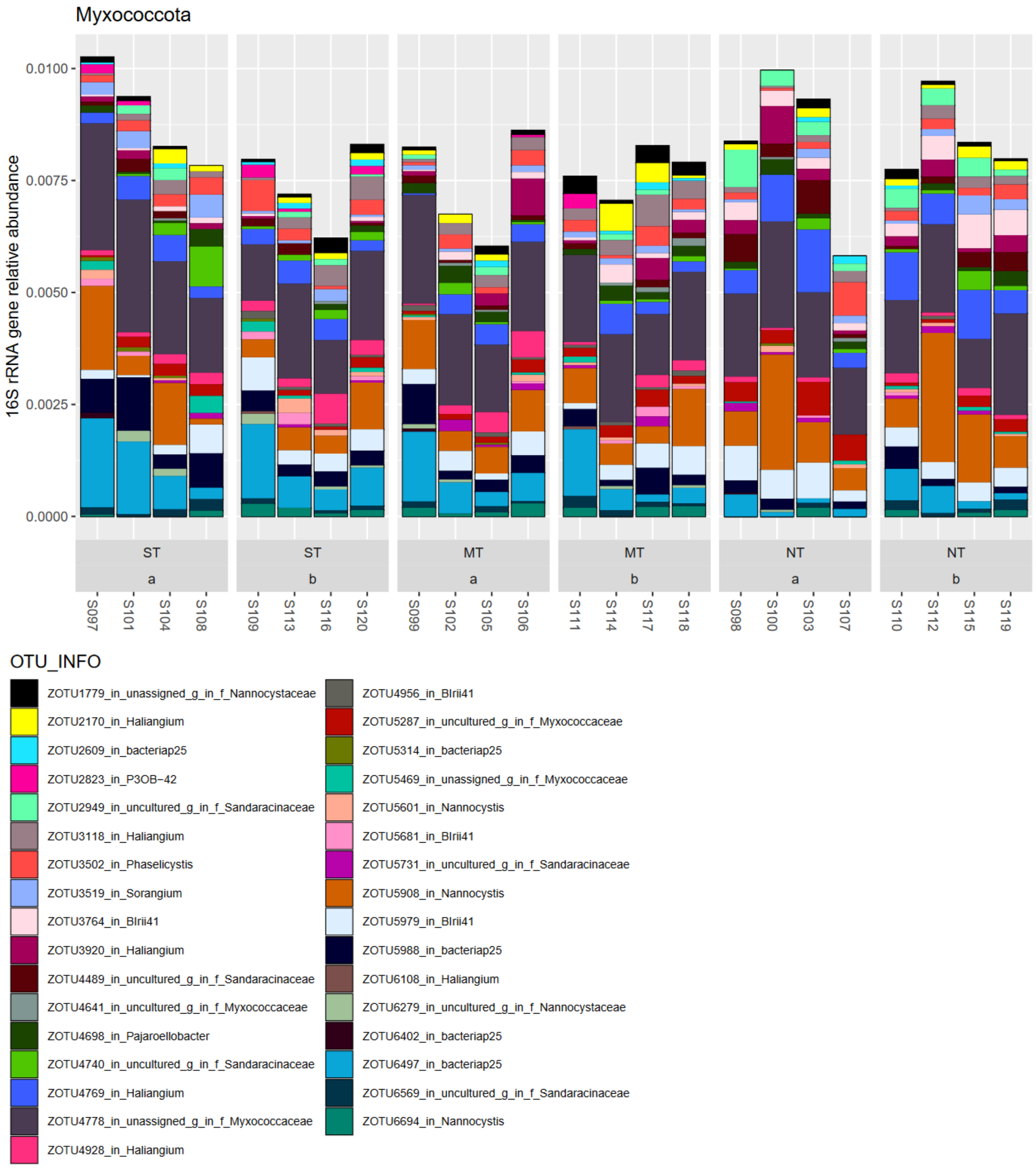


**Figure A18: Phylum Myxococcota.**

##### 1.1.6 Chloroflexi – not all are photosynthetic

In this dataset, Chloroflexi 16S amplicons were as abundant as those from the class Proteobacteria (8.6% vs. 8.5%). The classes in descending relative abundance were Chloroflexia (*sic*), KD4-96, Anaerolineae, Gitt-GS-136, TK10, Dehalococcoidia, JG30-KF-CM66, Ktedonobacteria, P2-11E, and OLB14.

The classes that belong to the phylum Chloroflexi *sensu stricto* are Chloroflexi (*sic*) and Thermomicrobia (in this dataset with Silva taxonomy, the order Thermomicrobiales are in class Chloroflexia, *sic*), while the classes Dehalococcoidetes, Anaerolineae, Caldilineae and Ktedonobacteria are regarded as associated (Gupta et al., 2013).

In the class Chloroflexia, four taxa from the orders Thermomicrobiales (etymologically indicating thermophilic growth) and Kallotenuales were significantly associated with tillage treatment factor (zOTUs 3790 and 2979 with decreases in NT vs ST and 3490 and 3083 with increases under NT).

Kallotenuales are formed by the type species *Kallotenue papyrolyticum*. This is a filamentous thermophile with aerobic chemoheterotrophic nutrition and the special ability to degrade paper and also otherwise grow on a range of polymeric and/or complex substrates (Cole et al., 2013).

Organisms in this class usually exhibit gliding mobility. They lack the genes for lipopolysaccharides and other key outer membrane functions. The order Chloroflexales contains organisms that are obligate or facultative phototrophs (Gupta et al., 2013).

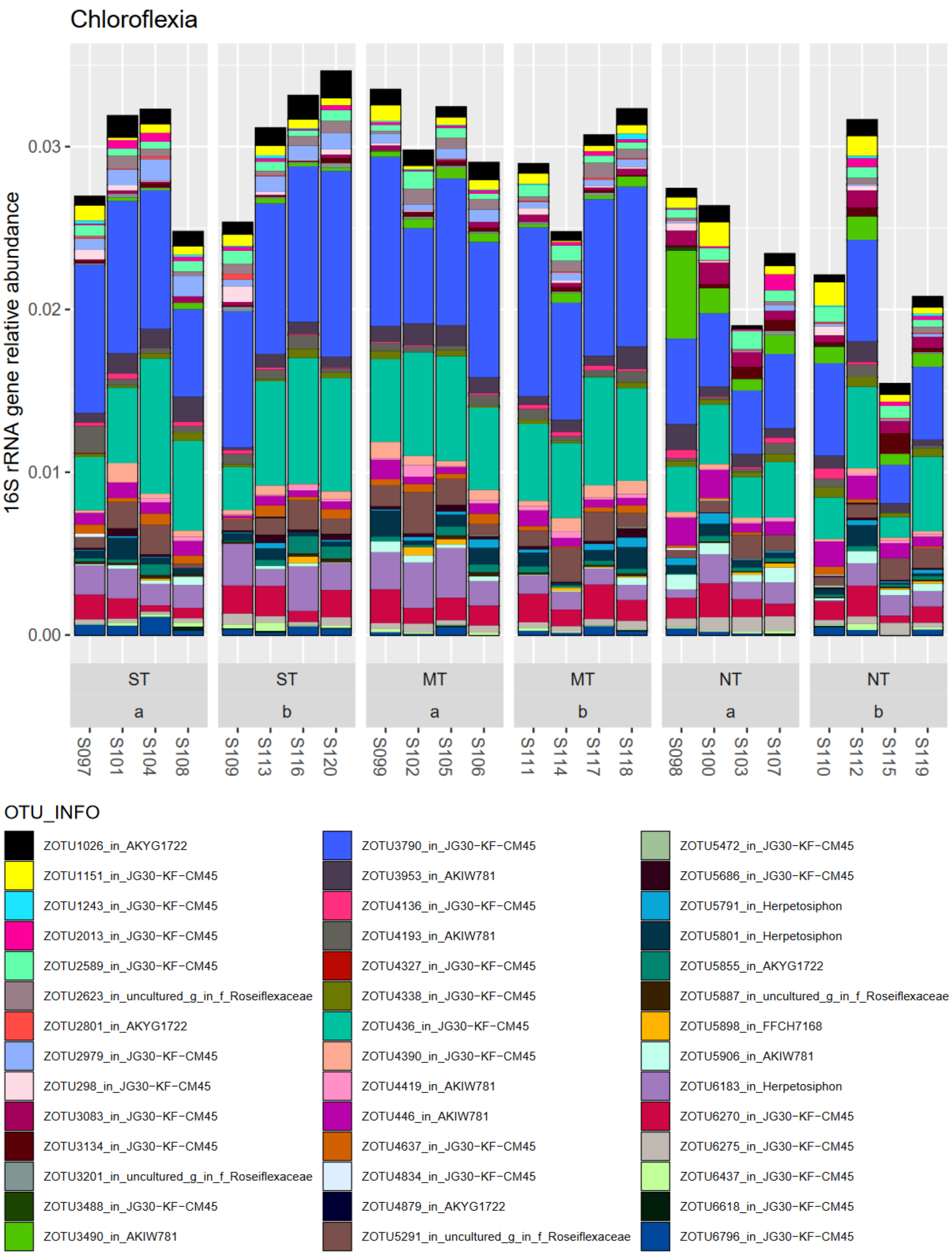


**Figure A19: Class Chloroflexia in phylum Chloroflexia**

The class KD4-96 contained two taxa with significant associations to clusters with relative reduction under NT management (zOTUs 1028, 931). KD4-96 is not a class in LPSN nor GTDB. A taxon from this group was found in another metabarcoding study to be strongly explaining soil health measures (Wilhelm et al., 2022).

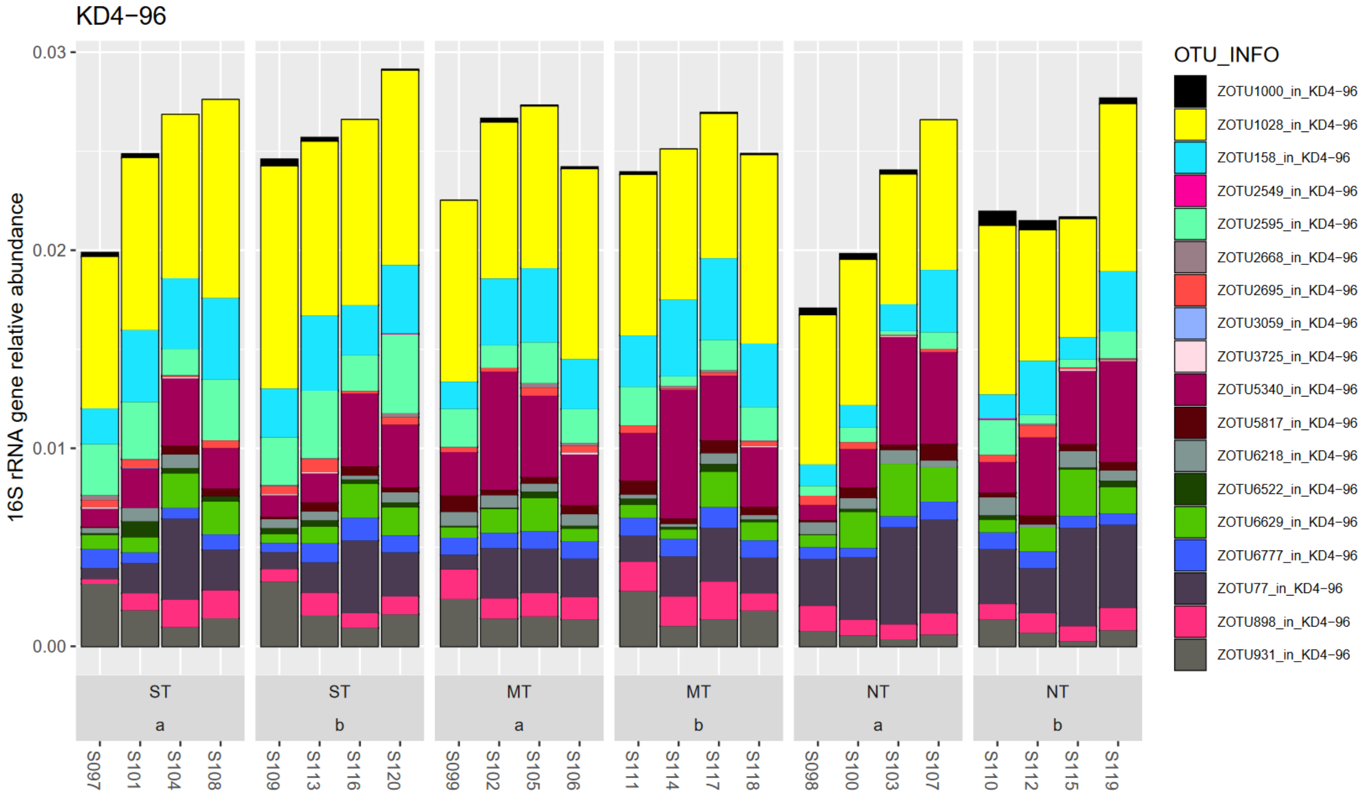


**Figure A20: KD4-96 in phylum Chloroflexi**

The class Anaerolineae contained three taxa with significant increases under NT (vs ST, particularly abundant OTU 5723) and 6 with decreases under NT (particularly abundant OTU 1228).

This class contains isolates that are likely anaerobes with chemoheterotrophic nutrition (Madigan et al., 2021, Yamada et al., 2006).

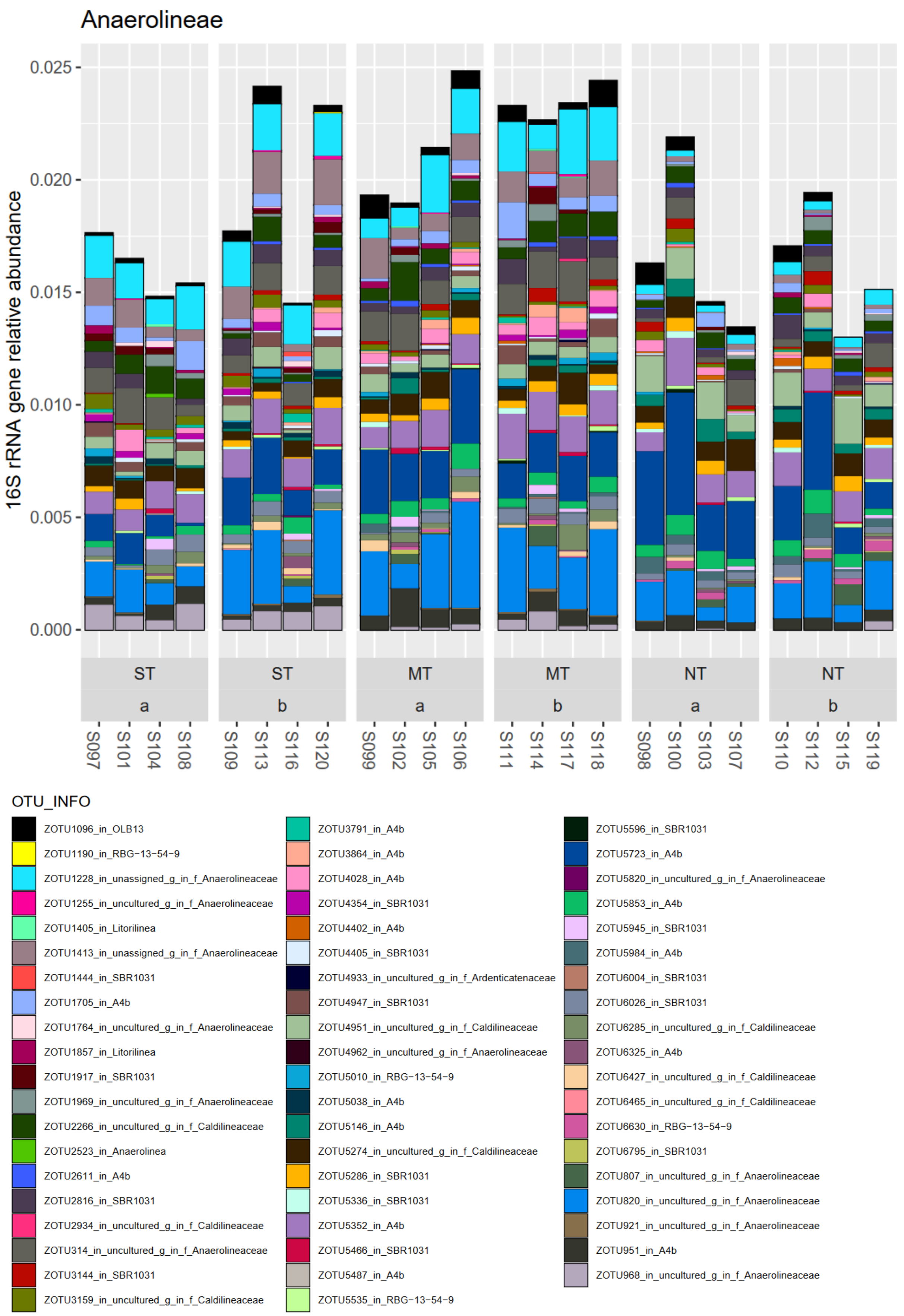


**Figure A21: Class Anaerolineae in phylum Chloroflexia**

This class exhibited one taxon (zOTU 1234) with significant association to tillage treatment, by a decrease in NT vs ST in top soil layer.

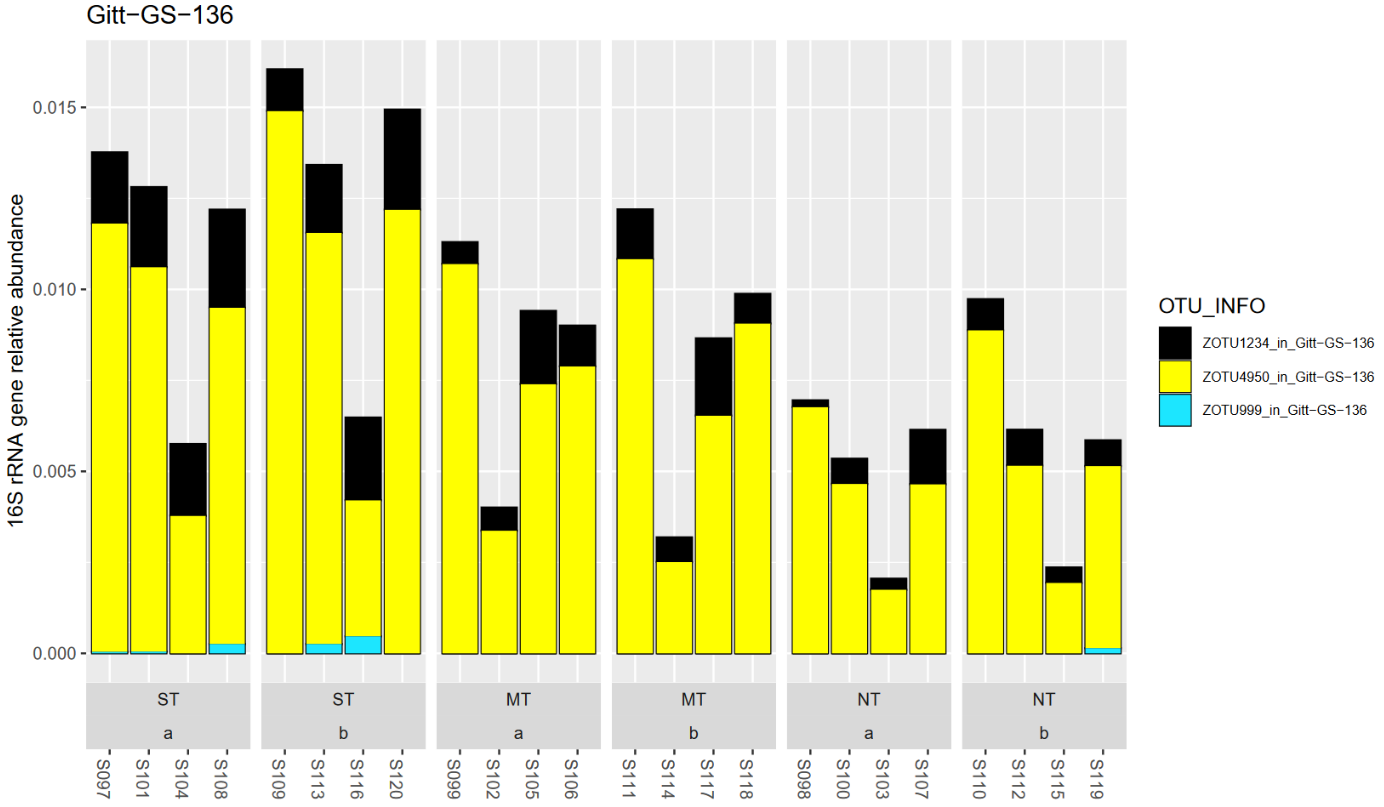


**Figure A22: Class Gitt-GS-136 in phylum Chloroflexia.**

This class did not exhibit significant changes with tillage treatment (LinDA, FDR>5%).

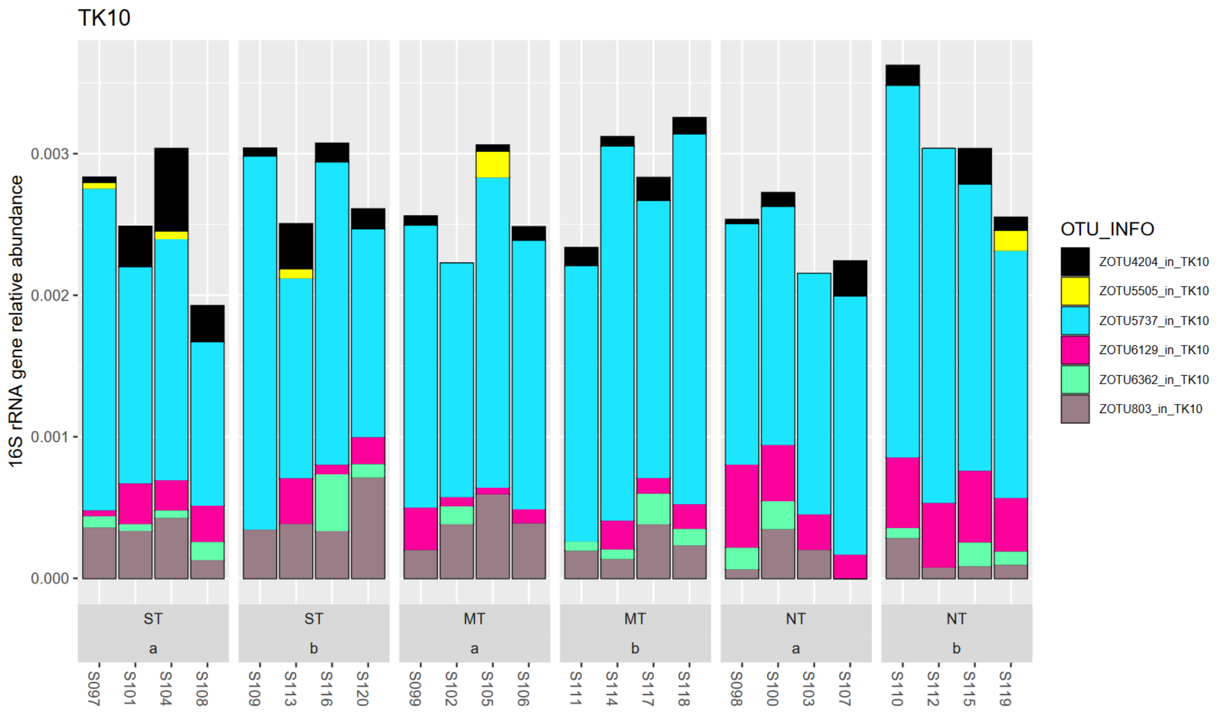


**Figure A23: TK10 in phylum Chloroflexi**

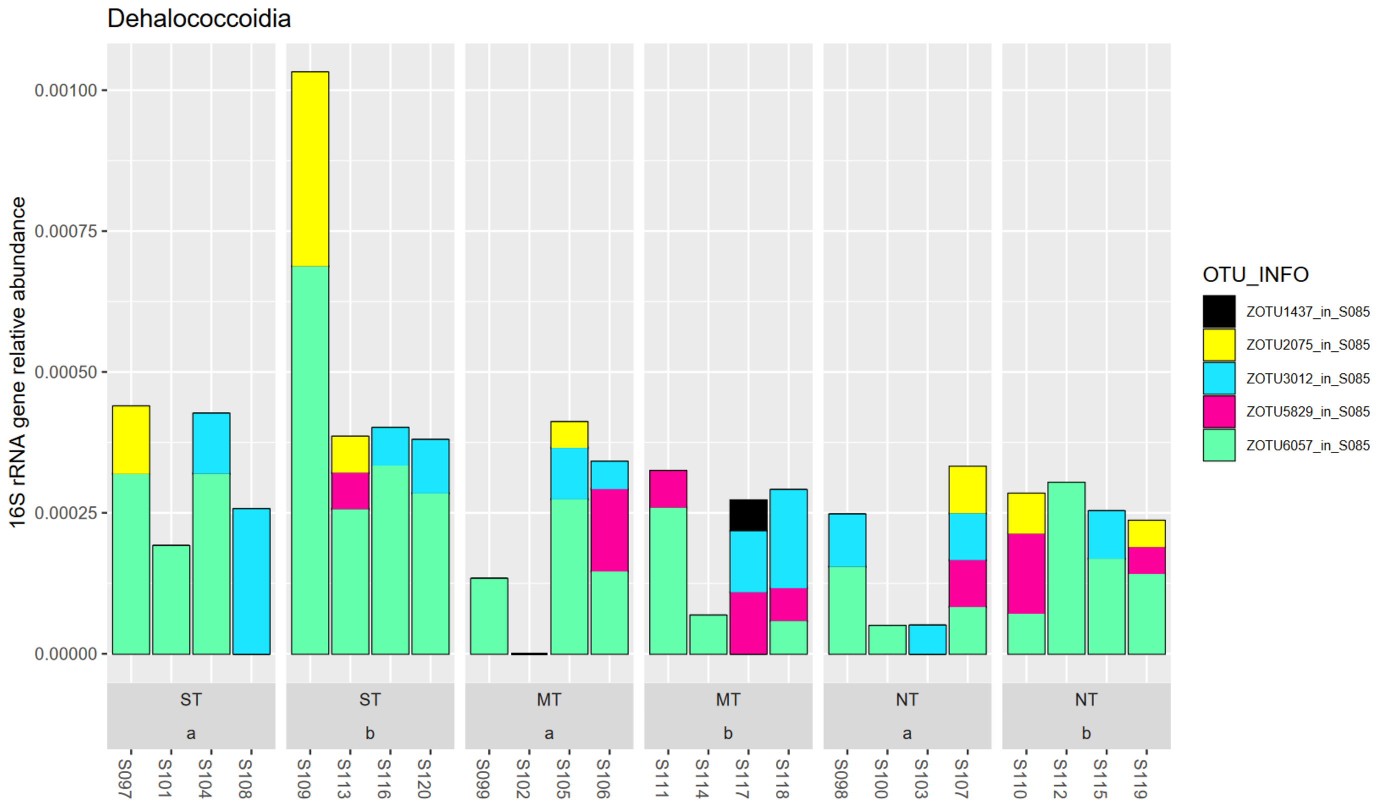


**Figure A24: Class Dehalococcoidia in phylum Chloroflexia**

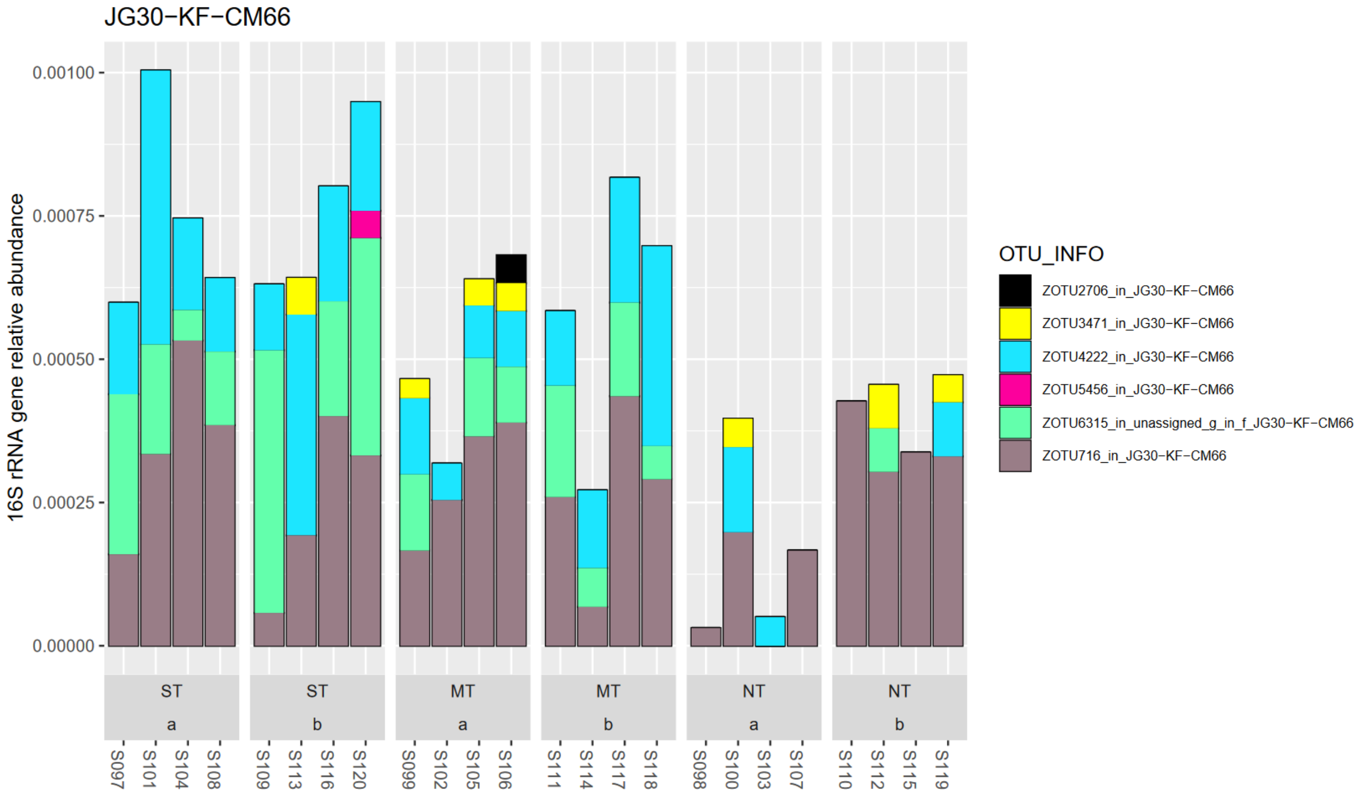


**Figure A25: Class JG30-KF-CM66 in phylum Chloroflexia.**

This class exhibited two taxa with significant decreases to NT vs ST management contrast (zOTU 1585, 4180)

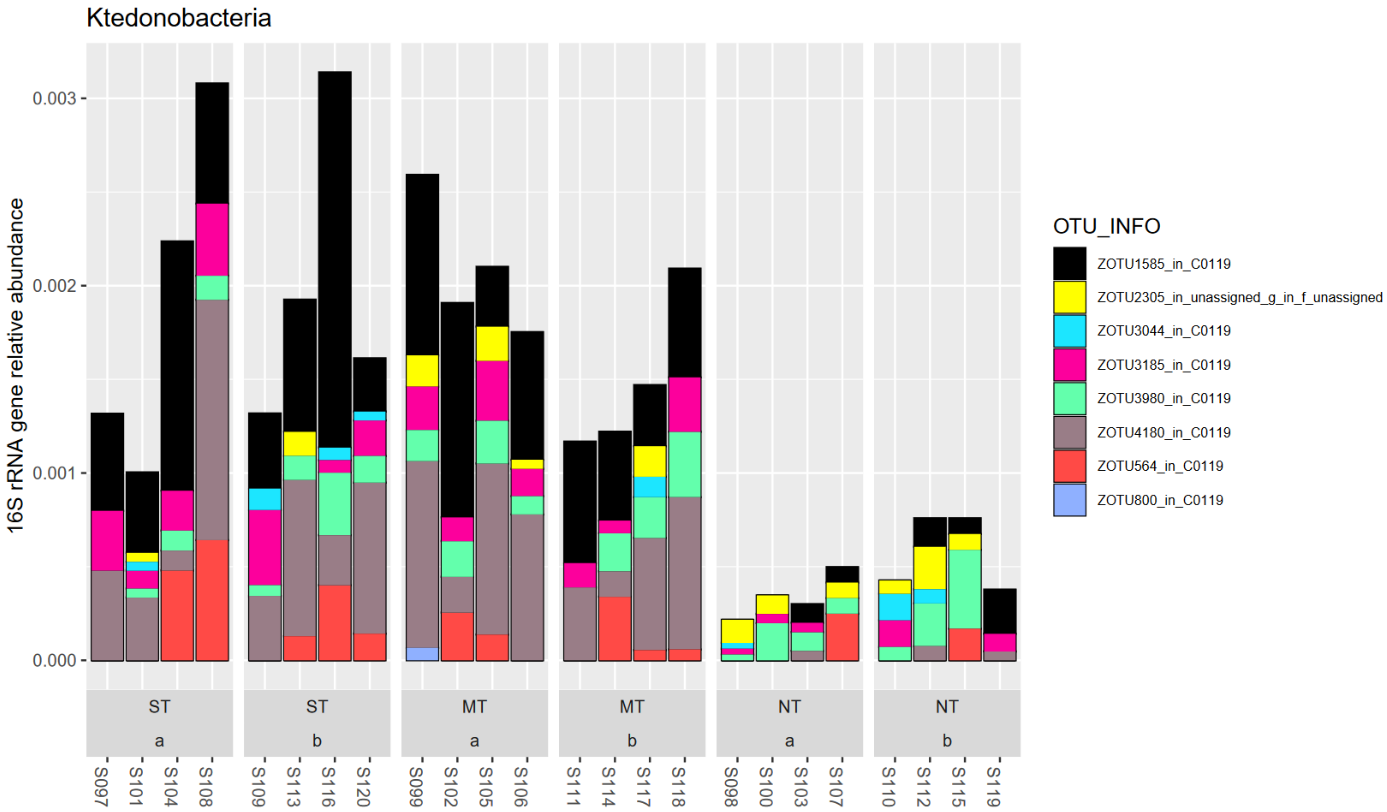


**Figure A26: Class Ktedonobacteria in phylum Chloroflexia.**

In this class, zOTU 403 was found to be significantly changing with tillage treatment factor (decrease NT vs ST or MT).

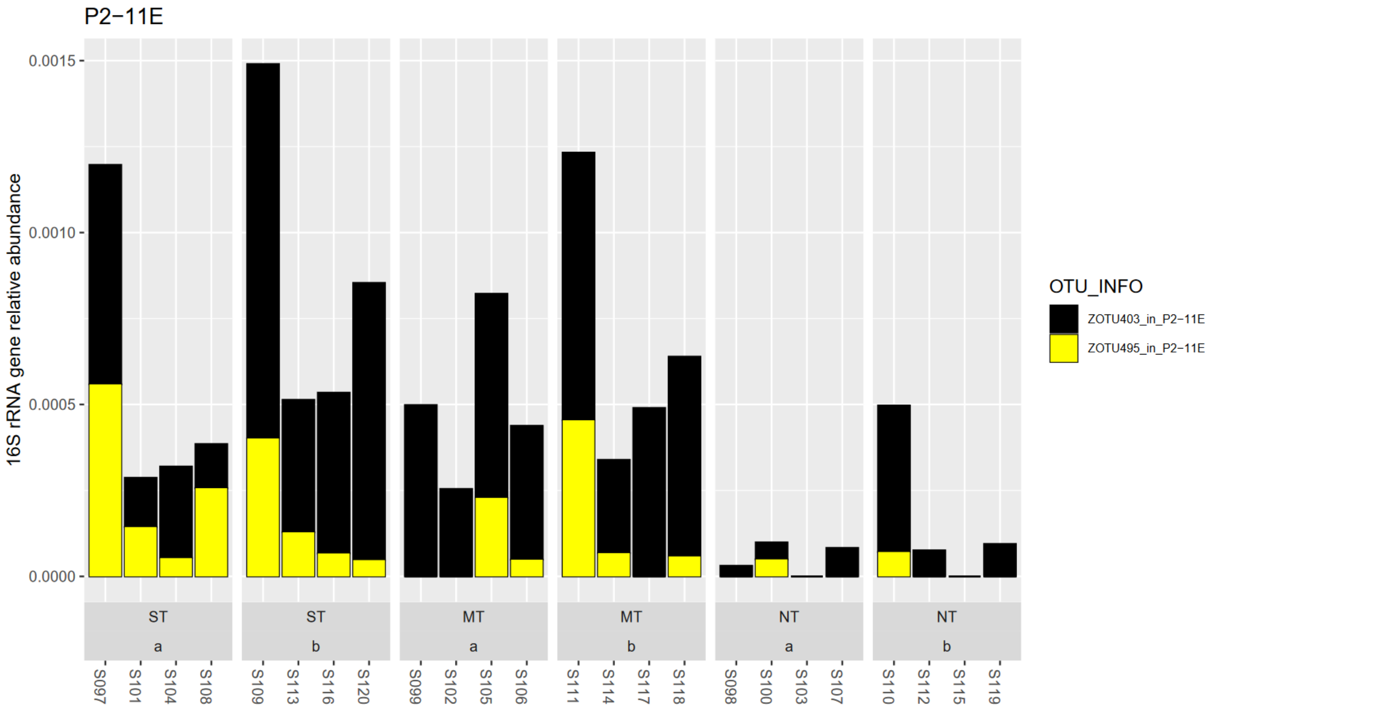


**Figure A27: Class P2-11E in phylum Chloroflexia.**

No significant changes with tillage management were detected in class OLB14 of the Chloroflexi phylum.

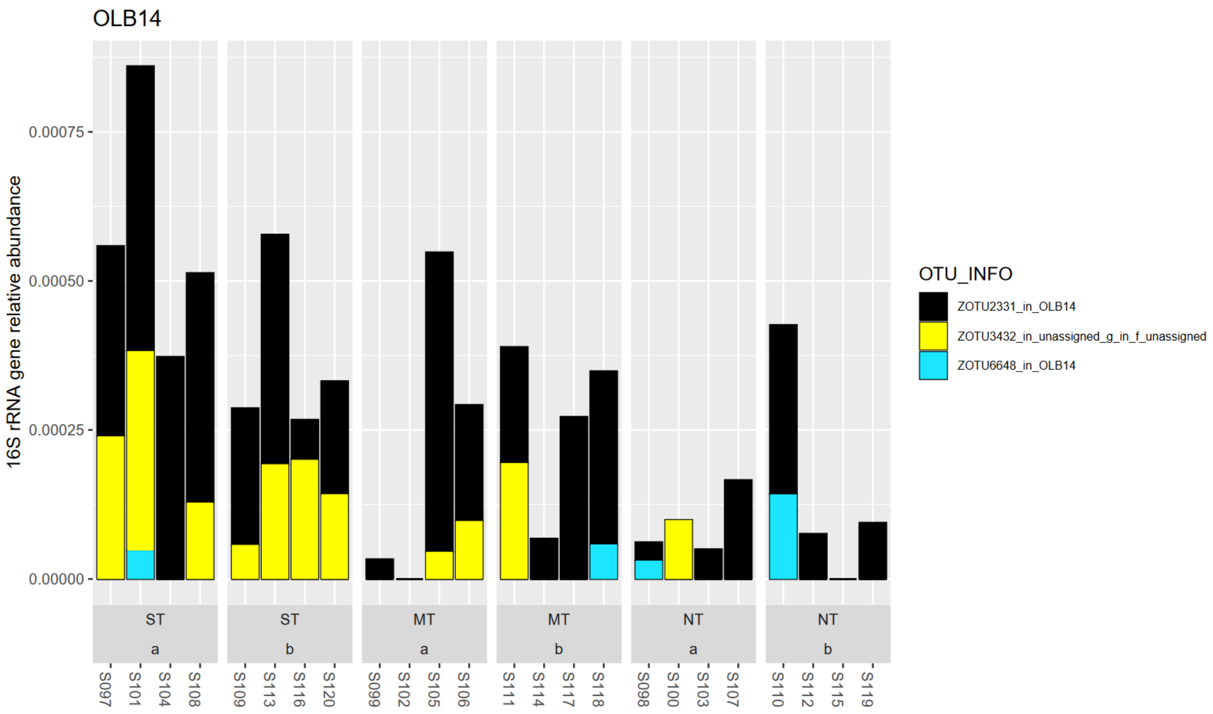


**Figure A28: OLB14 in phylum Chloroflexi**

##### 1.1.7 Nitrospirota – N cycle

In the phylum Nitrospirota, no significantly changing taxa with tillage treatment were detected (LinDA, FDR>5%). Generally, the 16S amplicons of this phylum were not relatively abundant in the dataset (at ca. 0.1%).

The phylum Nitrospirota could be of interest to N cycles due the capacity to perform nitrite oxidation to nitrate (second step of nitrification). These organisms are suspected to be most important in the nitrification fluxes of nitrogen rich environments (soils and waste water). Some organisms have also been reported to perform complete nitrification (comammox) (Madigan et al., 2021).

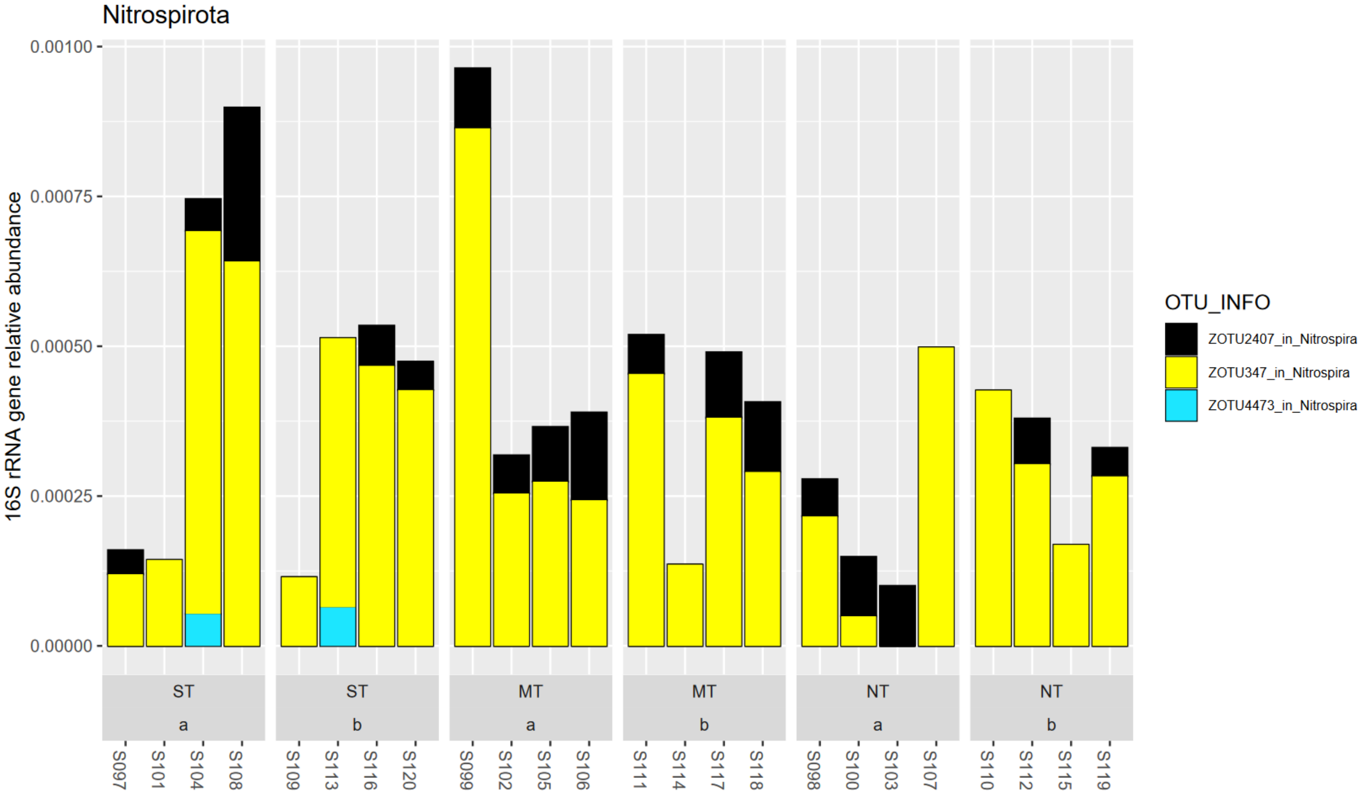


**Figure A29: Phylum Nitrospirota.**

##### I.I.8 Planctomycetes – widespread in soil but poorly understood

In the bacterial amplicon dataset, the Planctomycetes phylum was the most relatively abundant representing 39% of all 16S amplicons.

Such a high relative abundance for this phylum was not observed in other studies in Mediterranean soil (Lasa et al., 2019, dal Cortivo et al., 2020, Visioli et al., 2020).

The tillage treatment factor was significantly affecting 39 taxa (LinDA FDR<5%). The taxa with significant changes were from the classes Planctomycetes, Phycisphaerae, OM190, and Pla4 lineage.

Planctomycetes are gram-negative bacteria that often have stalks, appendages, or rosettes, and generally unusual morphology such crateriform pits. They are widely found in soil but their ecology remains poorly understood. The Brocardiales can carry out anaerobic ammonia oxidation (anamox). In the class Planctomycetes, the genus *Gemmata*, is noted for its intricate three-dimensional structure, that is speculated to play a role in ingestion of macromolecules (Madigan et al., 2021).

The class Planctomycetes contained 16 taxa with significant associations to tillage factor (LinDA, FDR< 5%). Of those, 10 taxa were in clusters with increases under NT management and 6 in clusters with decreases under NT (vs ST). The most relatively abundant taxon in this class was from the family *Isosphaeraceae*. It did not exhibit significant changes with the tillage treatments. A relatively abundant taxon with increases under NT vs ST in top soil (p<5%, cluster 3) was from the genus *Gemmata*. Further taxa from the *Gemmataceae* family (*Gemmata*, *Fimbriiglobus*, unassigned) were also in the same cluster with significant increases under NT as well as taxa from the family *Isosphaeraceae* (*Tundrisphaera*, several unassigned genera). Notably, the family with decreases under NT were often from the *Pirellulaceae* family (some with genus level annotation *Pirellula*)

The family *Gemmataceae* contains the genera *Gemmata, Zavarzinella, Telmatocola and Fimbriiglobus* and are characterized by strict aerobic and chemoheterotrophic metabolism *(*Kulichevskaya et al., 2017*). Gemmata* strains have been usually isolated from aqueous environments such as bogs, fens and other water bodies (Ivanova et al., 2021)*.* The type strain (*G. obscuriglobus*) was hypothesized to attach to other biological forms and utilize simple sugars found on the host surface layer or secreted by other organisms (Franzmann and Skerman, 1984). *G. palustris*, was found to prefer polysaccharides as growth substrate such as pectin, xylan, lichenan, and xanthan gum, among other (Ivanova et al., 2021)*.*

*Fimbriiglobus* genus is characterized in addition to family description by moderately acidophilic and mesophilic growth conditions. The type strain *F.ruber* is able to use a range of simple sugars and polymers as C source, such as laminarin, starch and xylan, but not for example pectin, chitosan, chitin and cellulose (Kulichevskaya et al., 2017).

*Isosphaeraceae* were, as *Gemmataceae*, also isolated from an aqueous environment of peat bog with the same aerobic chemoheterotrophic nutrition and crateriform surface but the delineating feature being the absence of stalks (Kulichevskaya et al., 2016).

Original members of the *Pirellulaceae* family have been isolated from brackish aqueous environments with strict aerobic growth and have composed microbial biofilms of macro-algae and participated in the breakdown of sulfated polysaccharides. The order is characterized by chemoheterotrophic nutrition (Dedysh et al., 2020).

Members of this class have exhibited contrasting relative abundance changes with tillage management while generally being aerobic chemoheterotrophs. Their changes might be explained in fine-scale changes to the environment of their niche hypervolume (e.g. change in OM composition as some degrade specific simple sugars and/or polysaccharides that other taxa do not, or different water regime reflecting that most isolates to date were isolated from different aqueous environments).

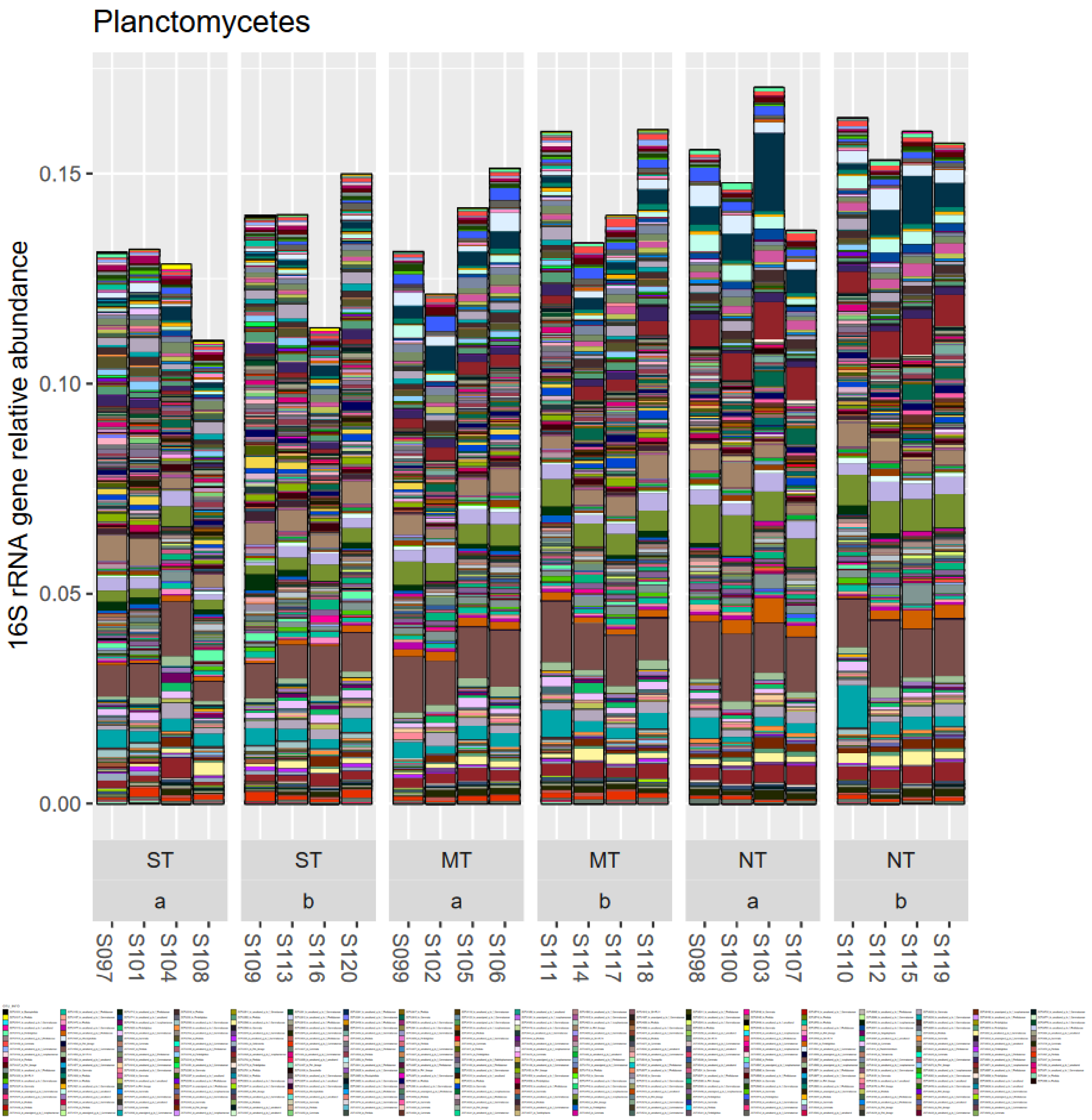


**Figure A30: Class Planctomycetes in phylum Planctomycetes.**

The class Phycisphaerae contained 17 taxa with significant associations to tillage management factor. Of these, 15 taxa were found in clusters with significant increases under NT management vs ST (cluster 3,6,7). All taxa belonged to the order Tepidisphaerales, with only one taxon further annotated to family level, *Tepidisphaeraceae*. All other taxa were assigned on sister family level to the “WD2101_soil_group”.

This class originates back to defining isolates of the type species *Phycisphaera mikurensis* with Gram-negative stain and facultative anaerobic metabolism on diverse sugars and capacity to reduce nitrate to nitrite (Fukunaga et al., 2009).

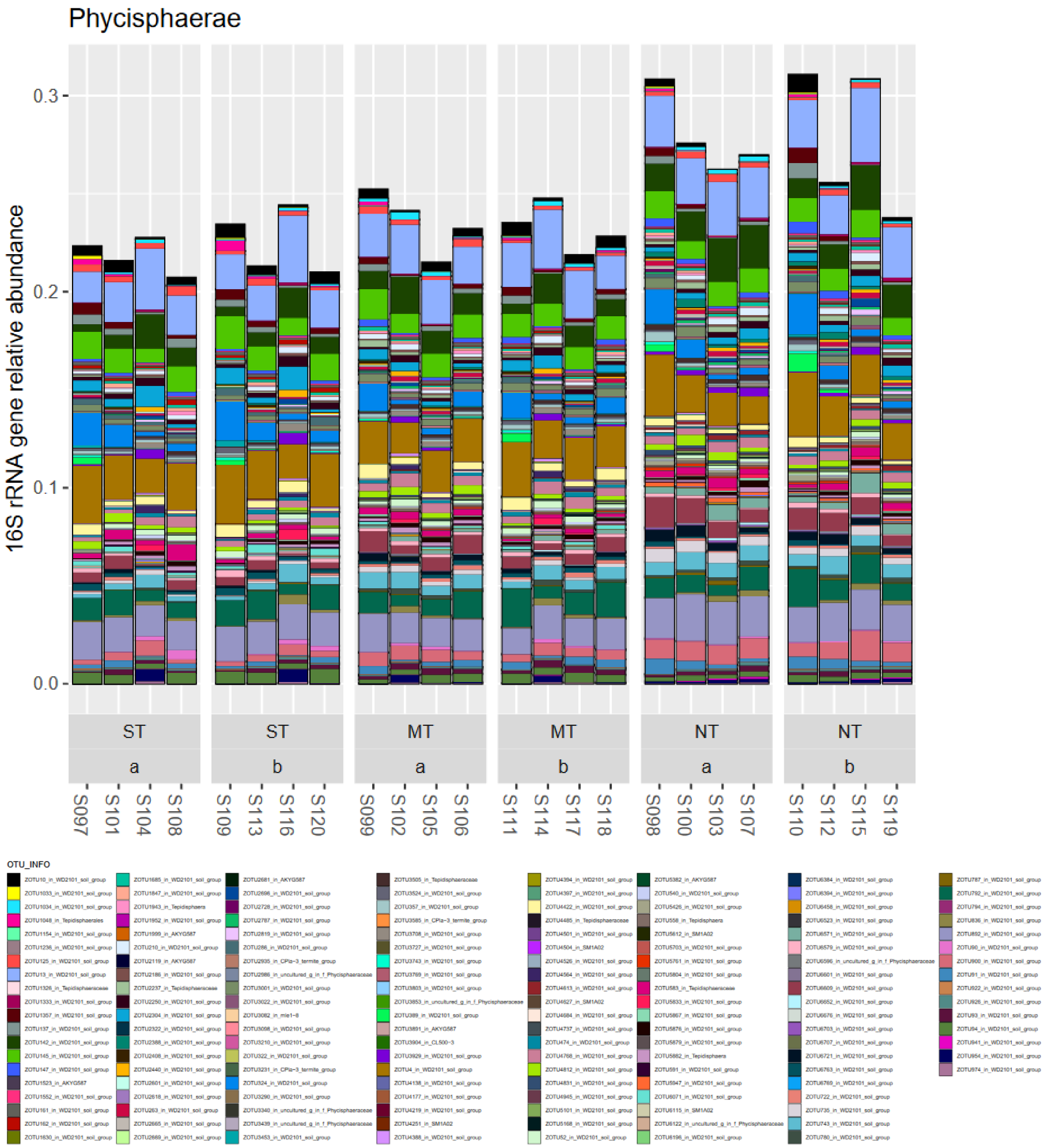


**Figure A31: Class Phycisphaerae in Phylum Planctomycetes.**

The class OM190 contained 4 taxa clusters with significant decreases under NT management (vs ST). This is not a class in LPSN or GTDB databases, where the latter has likely reclassed this grouping.

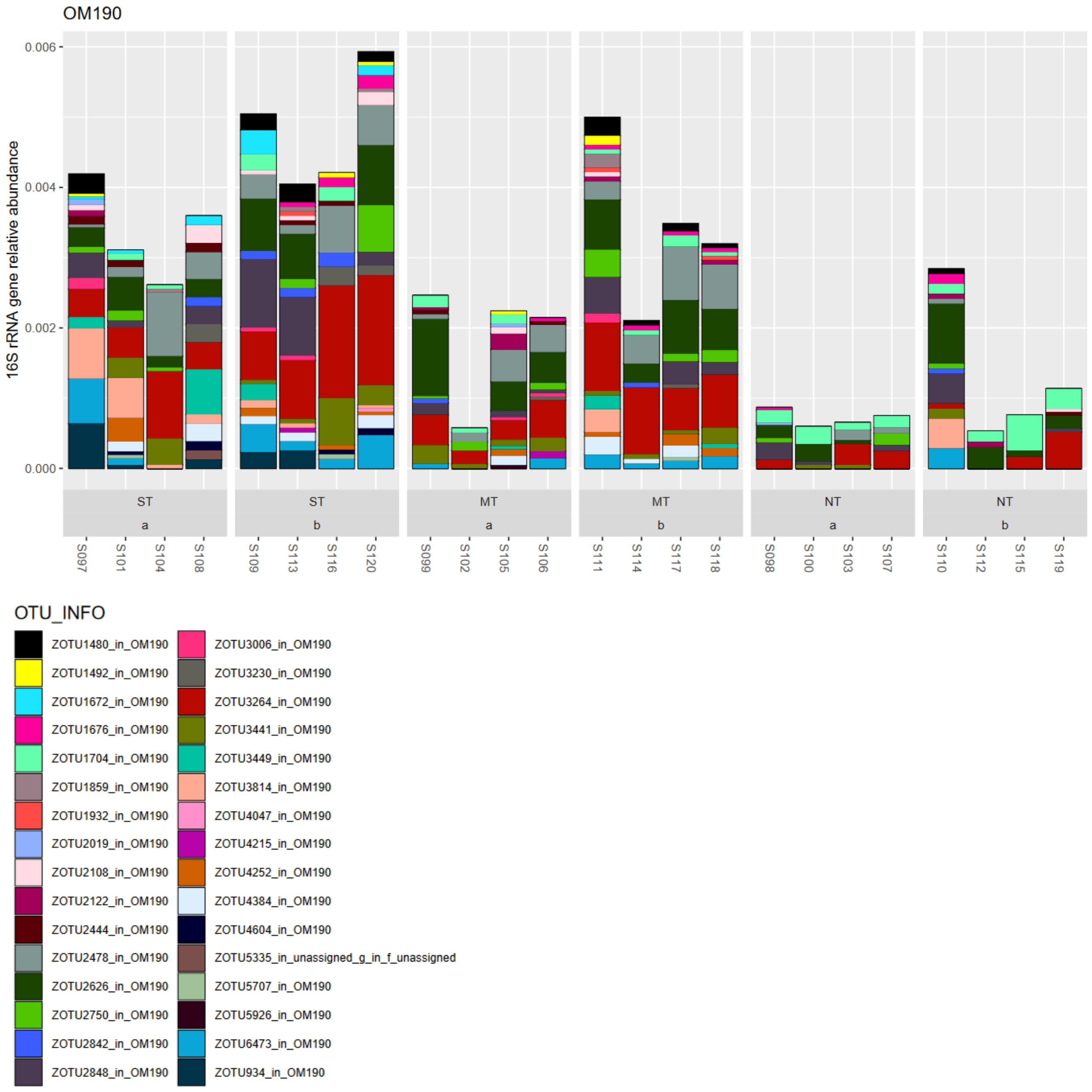


**Figure A32: Class OM190 in phylum Planctomycetes**

The class Pla4 contained 2 taxa in clusters with significant decreases under NT management (vs ST). Pla4 is not a class in LPSN nor GTDB, and has likely been reclassed in the latter as there remains a Pla2 class.

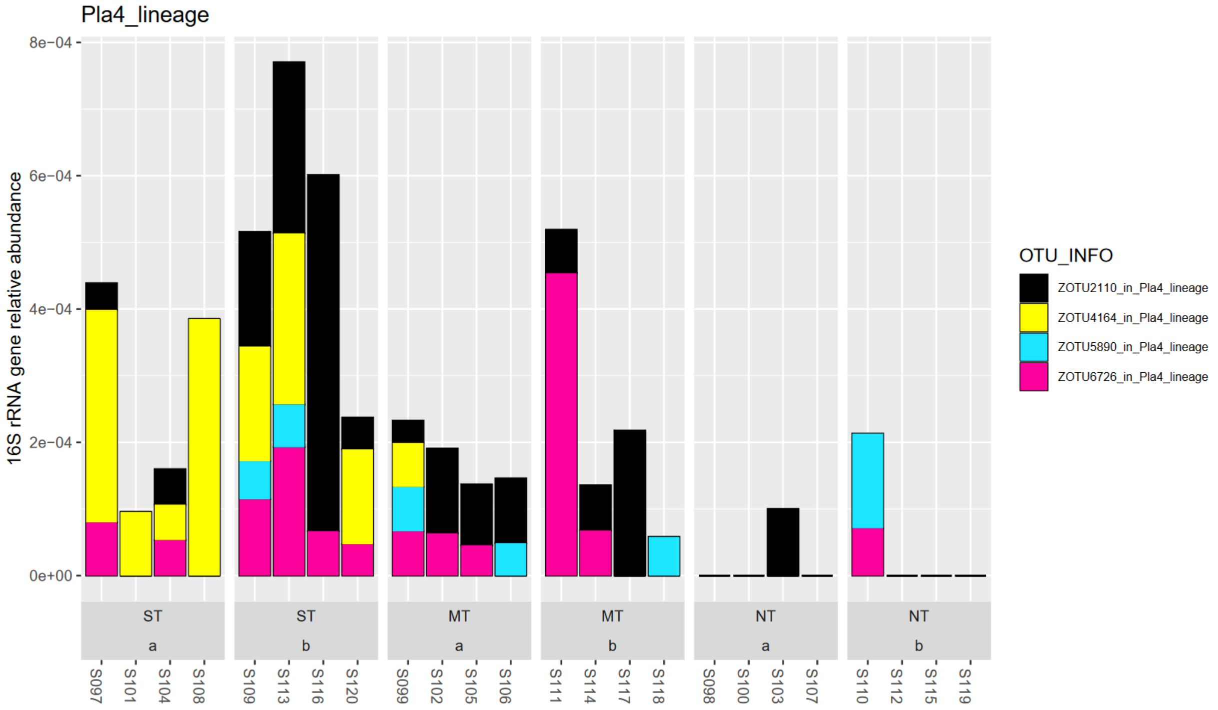


**Figure A33: Class Pla4 lineage in phylum Planctomycetes.**

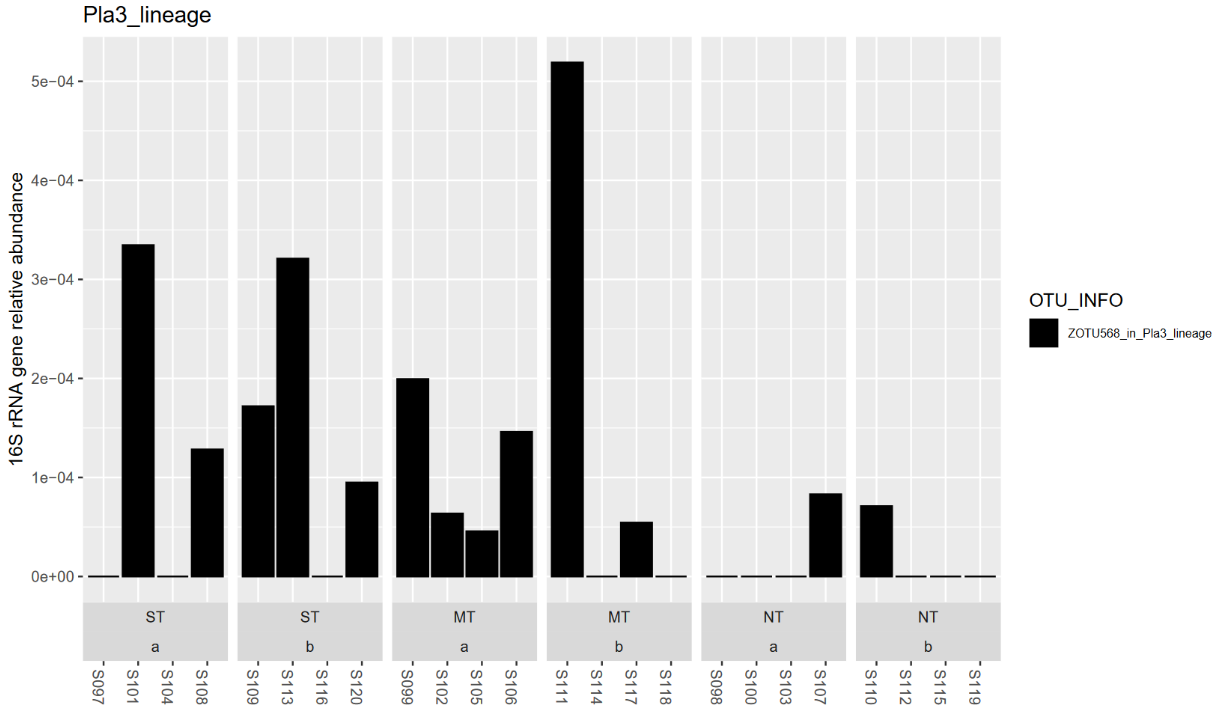


**Figure A34: Class Pla3 lineage in Planctomycetes.**

##### 1.1.9 Verrucomicrobia – high surface area for solute exchange

The phylum Verrucomicrobia exhibited 12 taxa with significant associations to tillage management factor (FDR <5%). All taxa came from the class Verrucomicrobiae. Of those, 7 taxa were associated with increases under NT (vs ST, 5 in cluster 6 with both NT and MT relative increases vs ST). Most taxa with relative increases in reduced tillage treatments, were from the family *Chthoniobacteraceae*. Three genus level annotations were made to *Chthoniobacter*.

The taxa with relative increases under tillage (ST) were from the families *Pedosphaeraceae* and *Verrucomicrobiaceae*.

The type strain for this genus, a *Chthoniobacter* isolate in Sangwan et al., (2004), was isolated from pasture soil and exhibited an aerobic metabolism of pyruvate, simple and polymeric sugars, but did not grow on amino acids, alcohols, or organic acids. It was able to store carbohydrates and P in inclusion bodies (Sangwan et al., 2004).

*Pedosphaeraceae* family is not found in LPSN but in GTDB. The genus *Pedosphaera* is only preliminarily assigned based on a not validly published species *Pedosphaera parvula* (Kant et al., 2011). The strain was isolated with a range of other strains from pasture soils placed in the class Verrucomicrobiae (Kant et al., 2011, Sangwan et al., 2005)). The isolates were able to grow aerobically on VL55 liquid media (Glucose, vitamins, and further nutrients) (Sangwan et al., 2005).

*Verrucomicrobiaceae* (family defining order) are Gram-negative bacteria isolated from aqueous phases of ponds, lakes, human, and marine environments. This family contains the genera *Prosthecobacter*, *Verrucomicrobium*, *Akkermansia*, *Rubritalea*, *Persicirhabdus*, *Roseibacillus* and *Luteolibacter*. Most are characterized as tolerant of oxygen, some genera are further detailed as explicitly strictly aerobic (Yoon et al., 2008).

The out-growing forms and appendages of Verrucomicrobia (etymology for warty) were argued to create a high-surface area for solute exchange (maybe scavenging/filtering solutes for nutrients) (Madigan et al., 2021). However, the prosthecate form was rare in several investigated strains of the class Verrucomicrobiae (Sangwan et al., 2004).

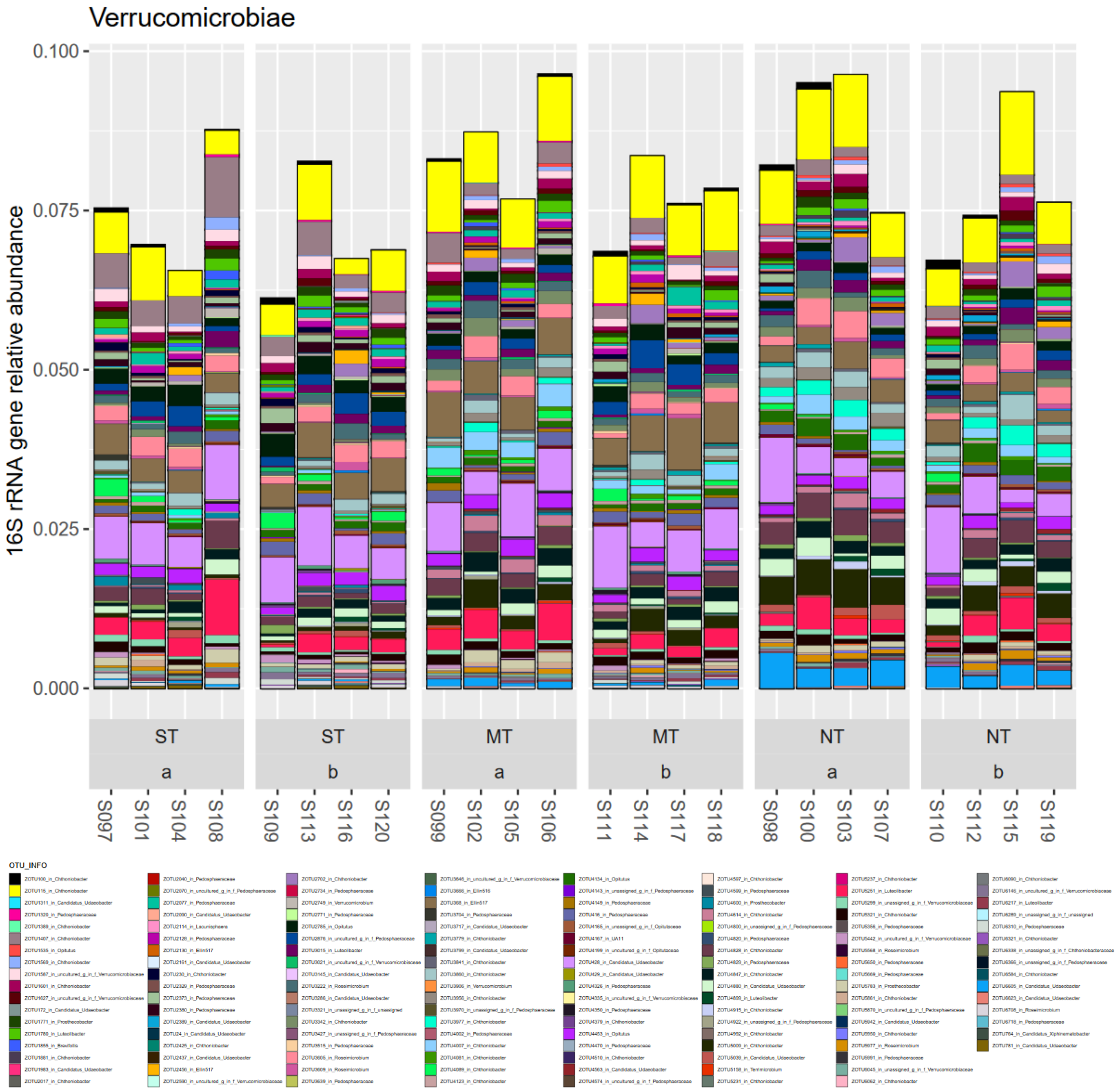


**Figure A35: Phylum Verrucomicrobiota**

##### 1.1.10 Cyanobacteria

The phylum Cyanobacteria consisted in this soil of 4 taxonomic units. Two taxa were associated with cluster 7 that exhibited strong increases under NT management (LinDA, FDR<5%). This was foremost a taxon from the genus *Tychonema* (SILVA taxonomy family *Phormidiaceae*, order Cyanobacteriales). The other taxon was less relatively abundant and was from the genus assignment *Microcoleus* strain PCC 7113 (SILVA family *Coleofasciculaceae*, order Cyanobacteriales). With recent phylogenomic and polyphasic analysis this polyphyletic phylum has been re-arranged (Strunecky et al., 2023, Cheng et al., 2022). The family *Phormidiaceae* was removed and *Tychonema* (*T.bourrelyi* CCAP 1459) was placed in the *Microcoleaceae* family and order Oscillatoriales (Strunecky et al., 2023). The false "*Microcoleus*" strain PCC 7113 was replaced from the family *Coleofasculaceae* to the *Wilmottiaceae* and order Coleofascicucales (Strunecky et al., 2023). The true *Microcoleaceae* are characterized by abundant occurrence of mucilaginous sheats and contains numerous filamentous growing genera such as *Microcoleus*, a relevant taxon in soil crusts. Interestingly as *Tychonema* is monophyletic with *Microcoleus vaginatus* (Strunecky et al., 2023, Zhang et al., 2016), but usually found in plankton, resolving the ecology and phylogenetic relationship is outstanding (Strunecky et al., 2023). Both, are as Cyanobacteria however, phototrophic and found in soil biocrusts (Büdel, 2005). Especially *M.vaginatus* was frequently found to be most abundant in biocrusts (Gundlapally and Garcia-Pichel, 2006). *Microcoleus* are characterized as UV-sensitive that reside mainly beneath the soil surface to avoid intense solar radiation and wind, and glide to the surface only when conditions are moist (Powell et al., 2015).

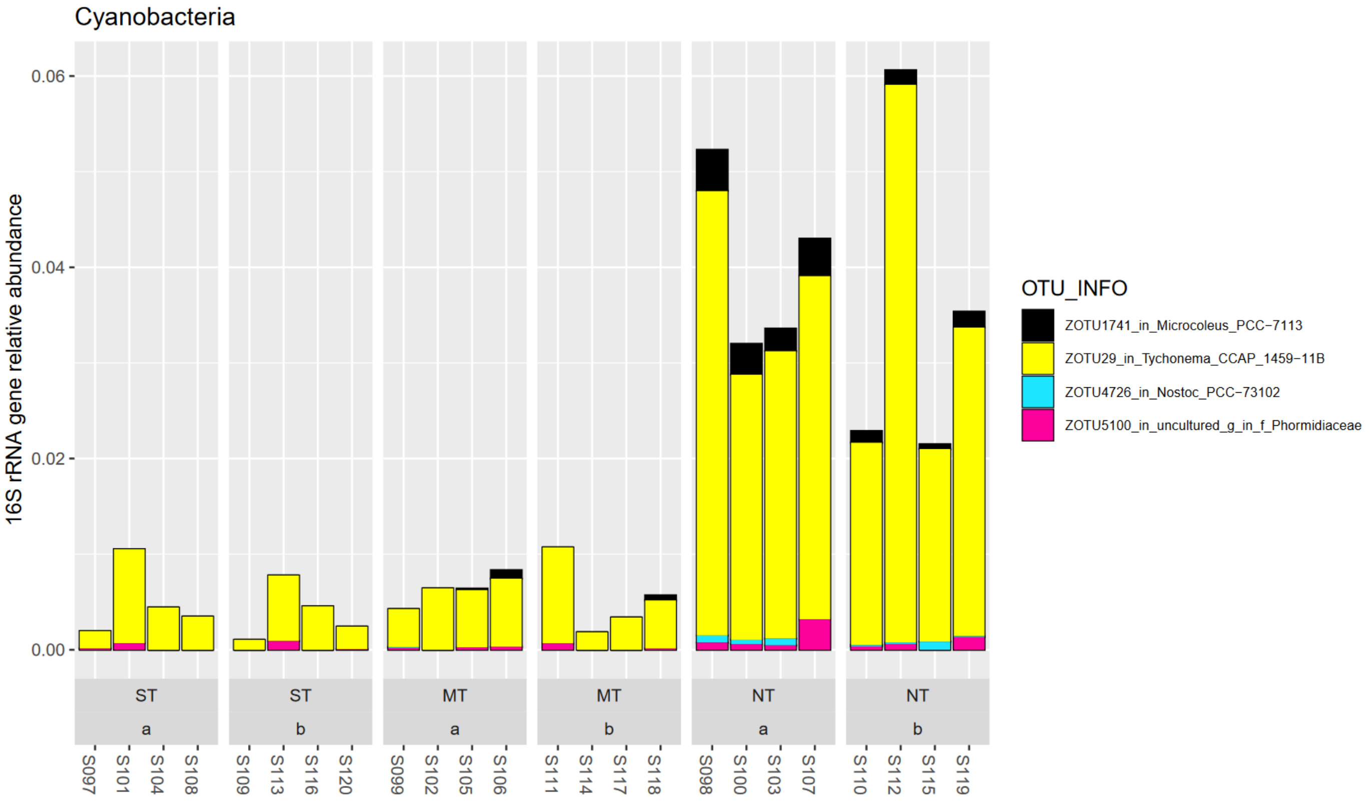


**Figure A36: Phylum Cyanobacteria**

#### 1.2 Differential abundance of bacteria

Results of differential abundance tests were also visualized with interactive Krona plots (Ondov et al., 2011). These can be opened with any current web browser and show the relative abundance of the marker gene, log-fold change in color code on per taxon resolution, and the taxon classifications. Files are made for the contrasts between tillage treatments and for each depth. Taxa with significant association to contrasts as determined by LinDA at 5% FDR are indicated by a suffix (e.g. NTSTa for NT vs. ST in depth a).

Bacteria: B1-B9_Bacteria_Krona.html

Fungi: F1-F9_Fungi_Krona.html

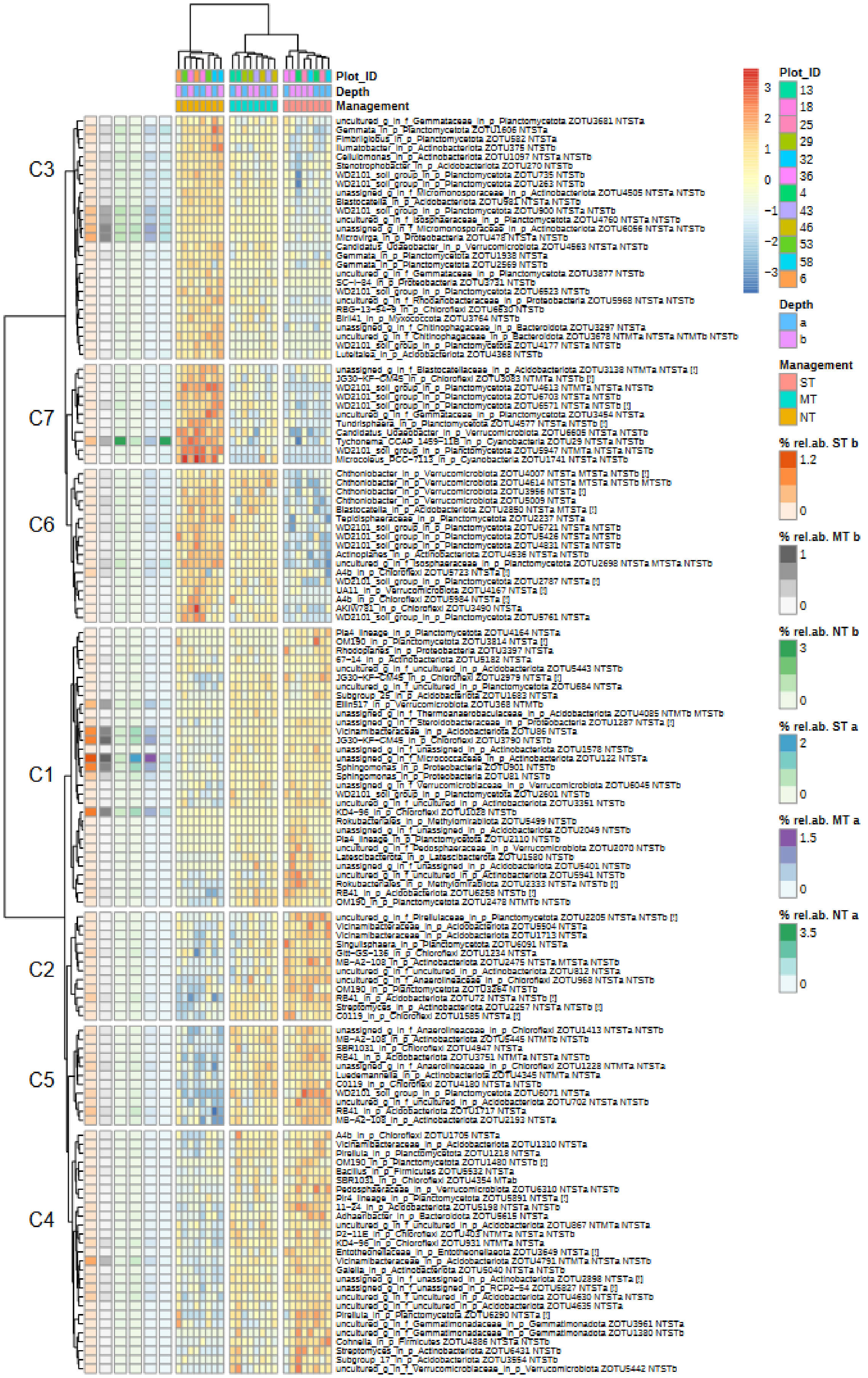

Plot_ID
Depth
Management
C3
C7
C6
C1
C2
C5
C4
3
2
1
0
−1
−2
−3
Plot_ID
13
18
25
29
32
36
4
43
46
53
58
6
Depth
a
b
Management
ST
MT
NT
% rel.ab. ST b
1.2
0
% rel.ab. MT b
1
0
% rel.ab. NT b
3
0
% rel.ab. ST a
2
0
% rel.ab. MT a
1.5
0
% rel.ab. NT a
3.5
0

**Figure A37: Bacteria, heatmap of log2-fold changes of taxon and sample against OTU-wise global mean. On the left average relative abundances of an OTU are resolved for treatment x depth combinations. Clustering of rows and columns was done with Euclidean distances and Ward`s clustering rule. Taxonomic information on OTUs is shown together with the contrast in which this taxon was indicated to be significantly differently abundant (e.g. contrast NT vs ST in depth a as NTSTa). Silhouette value was calculated, indicating the self-similarity of cluster members, and when negative indicated by [!] that the respective taxon could also be similar to a neighbor cluster. Clusters are indicated by “Ci” and correspond to numbering in Figure 2.**

#### 1.3 Taxonomic composition of fungi

The ITS2 region was amplified and sequenced. On average 44% of sequences were assigned to the phylum Ascomycota (lower in top soil at 40%), 25% to Basidiomycota (higher in NT top soil at 46%), 17% unassigned on phylum level, 5% to Chytridiomycota, 4% to Mortierellomycota (higher in ST at 6%), 3% to Glomeromycota (higher in lower depth 10-20 cm at 4%), 1% to Mucoromycota, 0.3% to Monoblepharomycota. Further fungal clades were also detected in lower relative abundance such as Aphelidiomycota, Basidiobolomycota, Olpidiomycota, Rozellomycota, and Zoopagomycota.

The phylogenetically most recent clade, subkingdom Dikarya, consists of the phyla Ascomycota and Basidiomycota. The next closely related subkingdom (Tedersoo et al., 2018) of Mucoromyceta consists of the phyla Glomeromycota, Mortierellomycota, Mucoromycota, and Calcarisporiellomycota. Chytridiomycota belong to the sub-kingdom of Chytridiomyceta that also contains Monoblepharomycota, Neocyllimastigomycota, and Olpidiomycota. The Aphelidiomyceta are phylogenetically a sister group to the Chytridiomyceta. Rozellomycota belong to a more basal fungal clade, the sub-kingdom Ophistosporidia (Tedersoo et al., 2018).

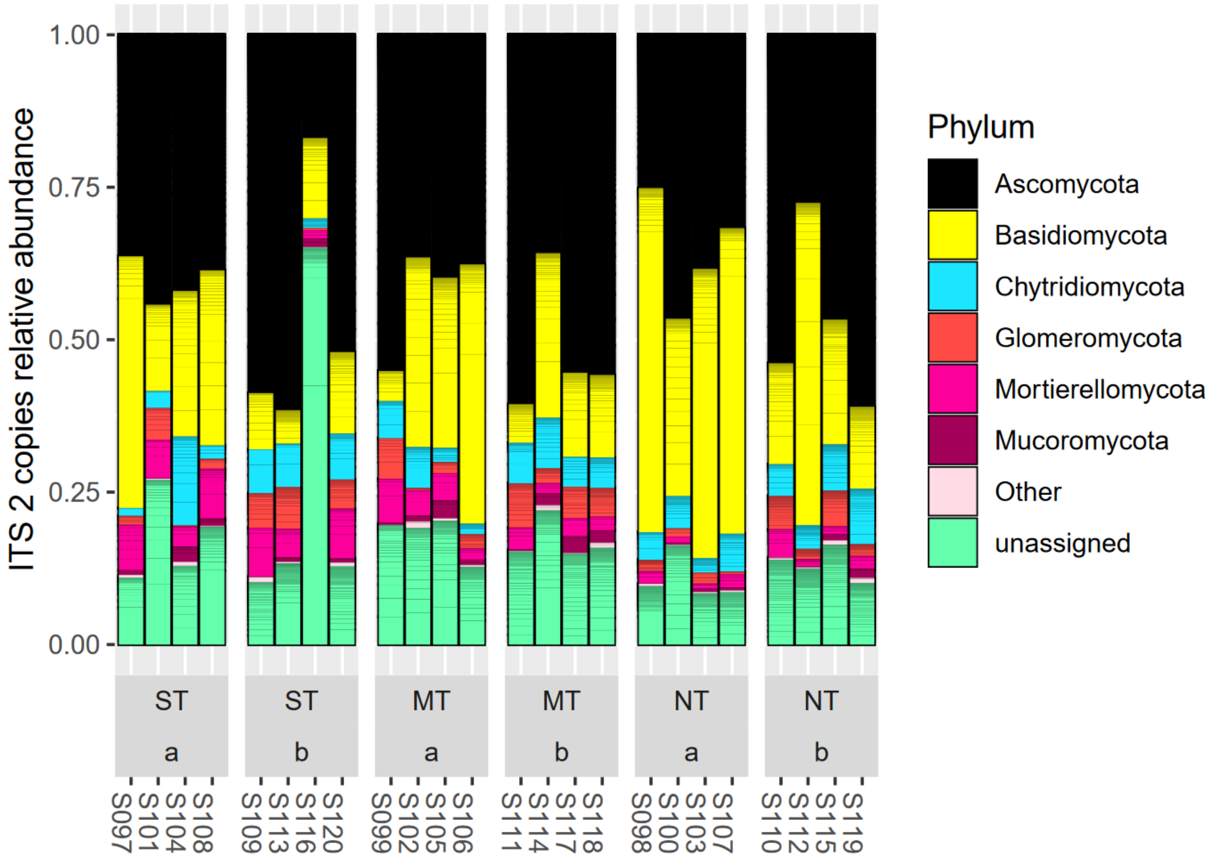


**Figure A38: Fungal ITS2 sequences classified on kingdom/phylum level. Phyla with less than 1% relative abundance were bulked as other.**

##### I.3.I Ascomycota – the largest and most diverse group containing free-living, opaque, endophytic and saprotrophic/disease types

This phylum was by relative abundance of ITS2 sequence counts, the most represented grouping of fungal sequences.

The following classes were found in the dataset in descending total relative abundance:

Sordariomycetes, Dothideomycetes, Leotiomycetes, Eurotiomycetes, unassigned class in Ascomycota, Pezizomycetes, Orbiliomycetes, Saccharomycetes, Lecanoromycetes, Taphrinomycetes, Laboulbeniomycetes, Arthoniomycetes, and Pezizomycotina class *incertae sedis*.

Ascomycota are the largest and most diverse group of fungi (Maddigan et al., 2021). Many are lichen forming (Grishkan et al., 2019).

Sordiariomycetes contained 8 taxa with significant changes to the tillage treatments (LinDA, FDR<5%).

Of those, 3 taxa were associated with clusters with relative increase under NT management (Cluster 1, Figure S2). One taxon was from an unassigned family from the order Xylariales and the other from the family *Delonicicolaceae* (order Delonicicolales). The third taxon was unassigned up to order level. These taxa were of medium relative abundance in this class with a maximum at 0.32% for the taxon from Xylariales under NT in top soil.

Three taxa were associated with a cluster exhibiting strong relative increases under MT management (cluster 4). These were a taxon from the genus *Fusarium* (family *Nectriaceae*, order Hypocreales) at 1.3% in MT depth b (10-20 cm, lower soil profile), a taxon from genus *Metacordyceps* (family *Clavicipitaceae*, order Hypocreales) at 0.8% in MT depth b, and a taxon from the genus *Achaetomium* (family *Chaetomiaceae*, order Sordariales) at 0.4% relative abundance in MT depth b. Further two taxa exhibited modest relative decrease under NT in lower soil layer (vs ST in 10-20 cm) with only one taxon assigned at order level to Microascales.

The most abundant taxon was at 5% from the order Sordariales with no significant changes from the tillage treatments (p>5%).

Generally, this class contains well-known genera such as *Trichoderma* (biocontrol agents, plant growth promoter, nutrient mobilisation; Harman, 2005), and *Fusarium* (often plant pathogenic). The order Sordariales is the most diverse in this class (Marin-Felix and Miller, 2022). It contains species that are ubiquitous and commonly found in soils and plant and insect hosts with a wide range of secondary metabolites production (Charria-Giron and Felix-Marin, 2022).

Specifically, Xylariales from the phylum Ascomycota with families *Xylariales, Xylariaceae,* and *Hypoxylaceae* have been described as endophytic or saprotrophic, and prolific producers of secondary metabolites spanning the full range of biochemical classes among which are key pharmaceutical compounds used (*viz.* non-growth associated biomolecules and highly regulated – could indicate K-strategist) (Becker and Stadler, 2021, Helaly et al., 2018). Indeed, endophytic Xylariales have been considered as biocontrol agents due to their antagonistic effects on fungi and other pathogens (Becker and Stadler, 2021).

The family *Nectriaceae* contains 55 genera, of which the majority are soil-borne saprobes, weak virulent facultative or obligate plant pathogens, and some facultative fungicolous and insecticolous. Among the genera are *Fusaria*, which contain important plant pathogen as well as medically important species (Lombard et al., 2015).

The genus *Metacordyceps* are generally recognized as insect pathogens and could be of use for biocontrol of insect pests (Kepler et al., 2011).

The *Achaetomium* were isolated from various environments such as swamps and arid soils and grew well as airy mycelium for example on malt extract agar (Cannon, 1986).

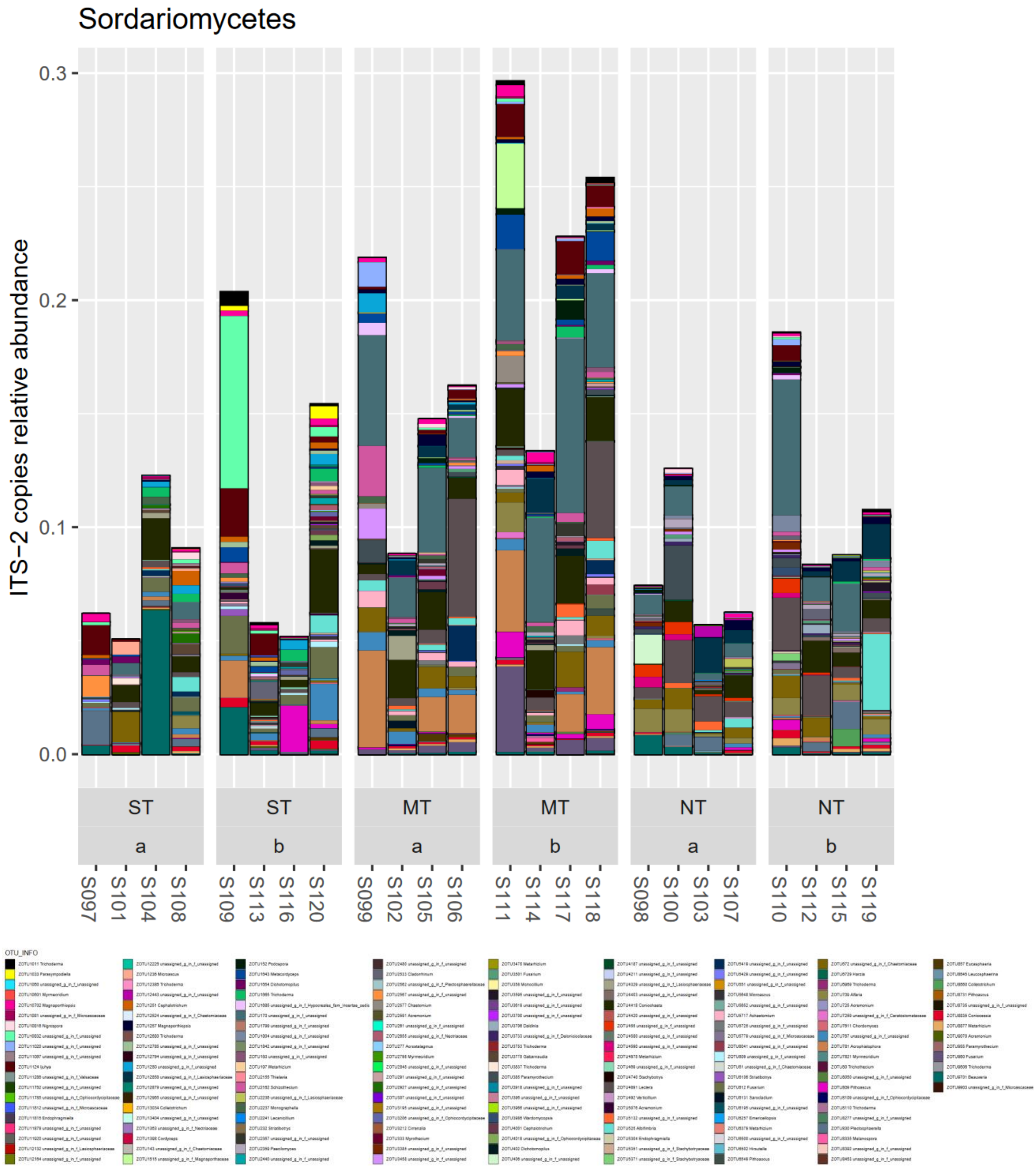


**Figure A39: Class Sordariomycetes in phylum Ascomycota**

In the class Dothideomycetes, 8 taxa with significant changes with tillage management were indicated ($p<5\%$). Most of these, 7 taxa, were associated with clusters with increase under NT management. Three were taxa from the genus *Keissleriella* (family *Lentitheciaceae*, order Pleosporales). Two more of those taxa were also from the order Pleosporales, with a genus *Sclerostagonospora* (family *Phaeosphaeriaceae*) and another taxon from the genus *Spegazzinia* (family *Didymosphaeriaceae*).

The one taxon with association to significant relative decrease under NT or vice versa strong increase under ST at 3.7% relative abundance in ST depth b was from the genus *Zymoseptoria* (hit to *Zymoseptoria tritici*, family *Mycosphaerellaceae*, order Capnodiales).

The class Dothideomycetes is the largest and most diverse grouping in the fungi with for example 191 families and over 19000 species. Living environments for members of this class span extremes (e.g. saline, rocks, marine), and genera living in soil such as *Ascochyta*, *Calophoma*, *Didymocyrtis*, *Paraboeremia*, and *Parathyridaria*, have been reported to be important to C and N nutrient cycles (Pem et al., 2021). None of the latter genera were found in the dataset, except for a *Neoascochyta* from the family *Didymellaceae*.

More specifically the order Pleosporales, encompasses important plant pathogens such as ***Alternaria (fam. Pleosporaceae)***, *Ascochyta (**fam. Didymellaceae)***, *Bipolaris (**fam. Pleosporaceae**)*, *Didymella (**fam. Didymellaceae)*** and ***Leptosphaeria*** (**fam. *Leptosphaeriaceae***) [highlighted in bold, were found in dataset]. The family *Didymellaceae* contains important plant pathogens of blackleg and ascochyta blight as well as endophytic, fungicolous and lichenicolous organisms (Valenzuela-Lopez et al., 2018). Taxa with strong increases under NT (cluster 5) were from the families *Didymosphaeriaceae* and *Phaeosphaeriacea, Spegazzinia radermacherae* and *Sclerostagonospora*, respectively. *Spegazzinia radermacherae* was isolated as a saprobe living on fallen seed pods of trees of *Radermachera sinica* (Hance tree, Asterids, Angiosperm) (Jayasiri et al., 2019). *Sclerostagonospora* (introduced as *S.Höhn* 1917, formerly *Hendersonia*, 1878, found on stems of perennial *Heracleum sphondylium* an umbelliferous plant, *Apiaceae,* Index Fungorum) is a pathogenic or saprobe of various monocotyledons and dicotyledons (Phookamsak et al., 2014).

The family in which *Keissleriella* species are placed, *Lentitheciaceae* (family, order Pleosporales), are known as saprophytes of herbaceous and woody plants (Calabon et al., 2021).

*Zymoseptoria* organisms are classified as a group of fungi, apart from the *Septoria* species, occuring on gramineous hosts and contain well-known plant pathogens such as *Z. tritici*, the causal agent of wheat blotch disease, Z. passerinii barley blotch disease and further species occurring on barley but also on weeds of these crops such as canary grass (Quadvlieg et al., 2011).

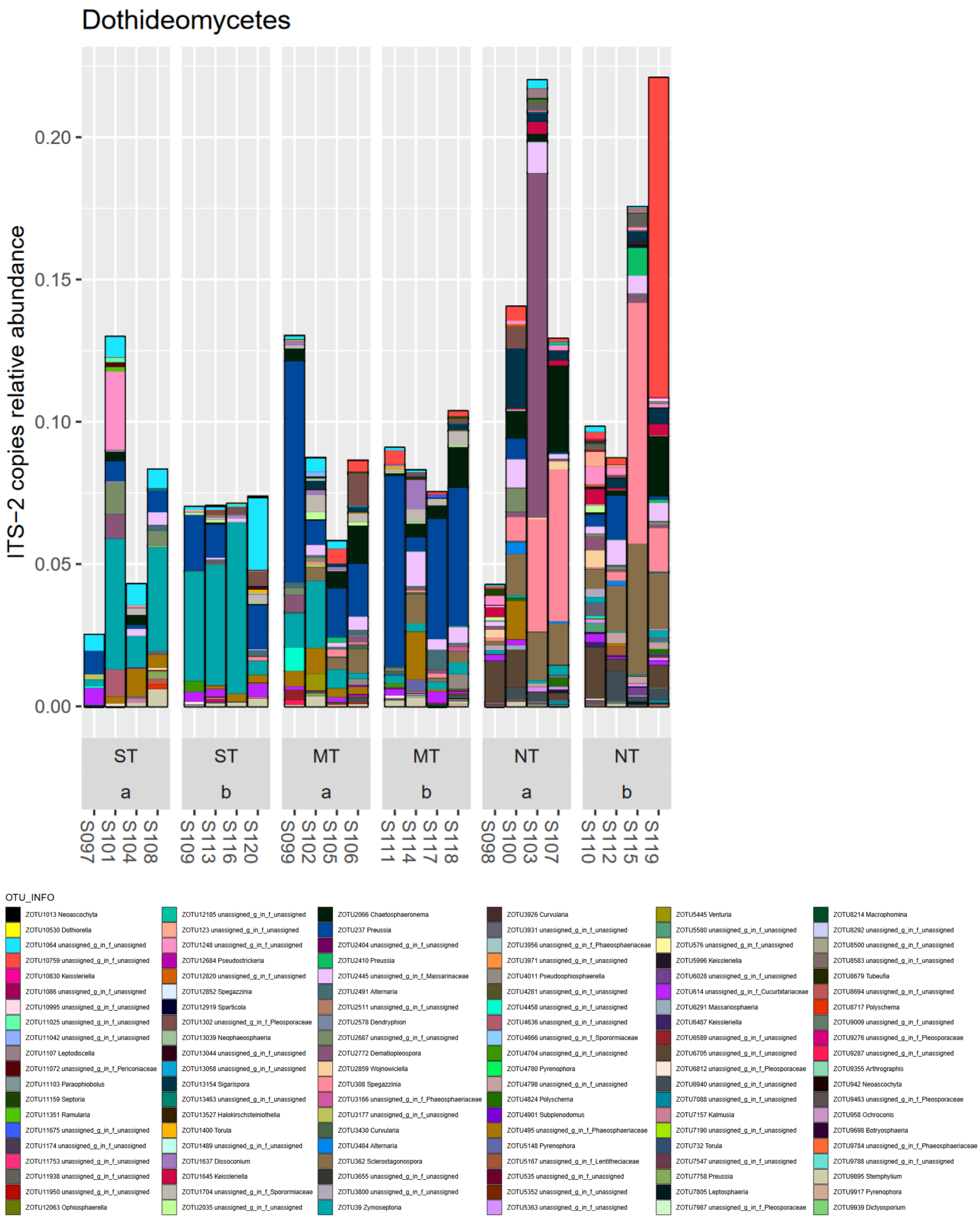


**Figure A40: Class Dothideomycetes in phylum Ascomycota**

In the class Leotiomycetes, two taxa with association to clusters with significant increase under NT were found (LinDA FDR<5%). Both were from the order Heliotales, one of those with assignment to *Clarireedia bennettii* (family *Sclerotiniaceae*). *Clarireedia bennettii* was the fourth most relative abundant taxon in this class at 1.3% in NT management in the top soil layer. The other was much less relatively abundant at below 0.4% relative abundance.

This class contains ecologically diverse members including mycorrhizae, endophytes, fungal parasites, root symbionts, and wood rot. Saprotrophs of plant material can be found (in dataset in bold) in the families ***Helotiaceae***, *Lachnaceae* and ***Hyaloscyphaceae***. Plant pathogens can be found in the Medeolariales, <u>***Sclerotiniaceae***</u>, **Erysiphales**, **Helotiales**, and Rhytismatales. Taxa of *Sclerotiniaceae* are saprobic or plant parasites of stems, leaves, flowers, fruits, seeds, and wood of various monocots and dicots (Ekanayaka et al., 2019). The *Clarireedia* are an economically important disease of C3 and C4 turfgrasses, causing the “dollar spot disease”. Particularly *Clarireedia bennettii* is specifically befalling C3 hosts of *Poaceae* family (i.e. Festuca rubrus) (Salgado-Salazar et al., 2018).

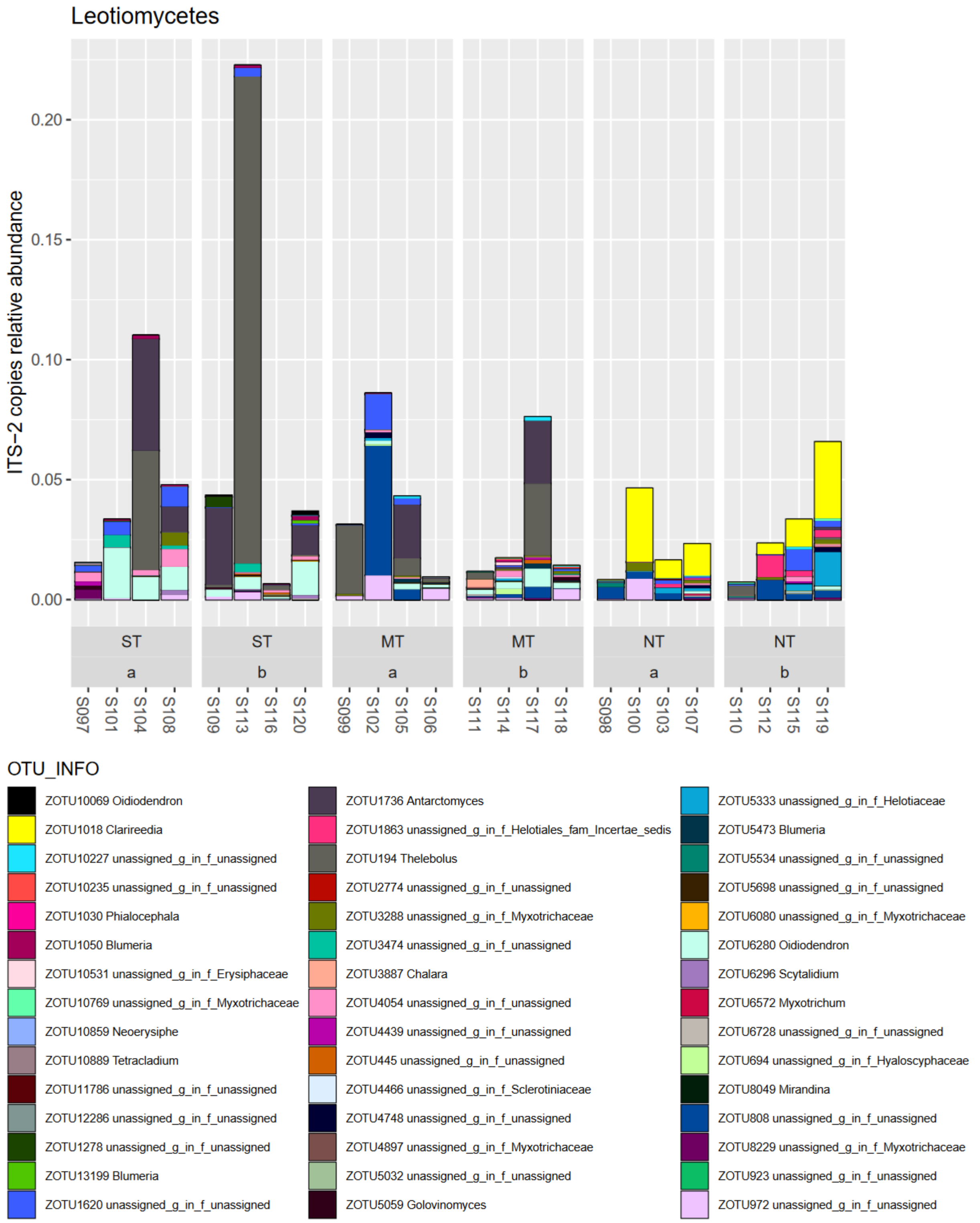


**Figure A41: Leotiomycetes in phylum Ascomycota**

In the class Eurotiomycetes, 3 taxa with significant associations to a cluster with significant decreases under NT / vice versa increases under ST and one taxon with significant associations to a cluster with moderate relative increase under NT were found. The latter taxon, *Exophiala crusticola* (family *Herpotrichiellaceae*, order Chaetothyriales), was at a low relative abundance <0.05%. It is discussed further in the section 1.2 on soil biocrust members. The former three taxa with increases under MT and/or ST, were *Aspergillus thesauricus* (family *Aspergillaceae*, order Eurotiales) at a relative abundance of 1.7% in MT depth b, a taxon from the order Onygenales at a lower relative abundance of 0.3% in ST depth b, and another taxon unassigned up to the class Eurotiomycetes at a relative abundance of 0.4% in ST top soil layer.

This class contains taxa such as *Penicillium* and *Aspergillus* from the family *Aspergillaceae*. They are well-known for antibiotic production and as bread-mould, respectively (Madigan et al., 2021). In soil, organisms of this family are frequently coprophilous on herbivore dung (Guevara-Suarez et al., 2019). *Aspergillus thesauricus* was isolated in a Spanish cave from air and growing on substrates such various human introduced organic sources such as hair, textile fibres, and candle residues. Isolates grew comparably fast on media such as yeast, malt, sucrose and creatine agars but did not exhibit strong pigmentation. They were shades of grey, green, to brown (Novakova et al., 2012).

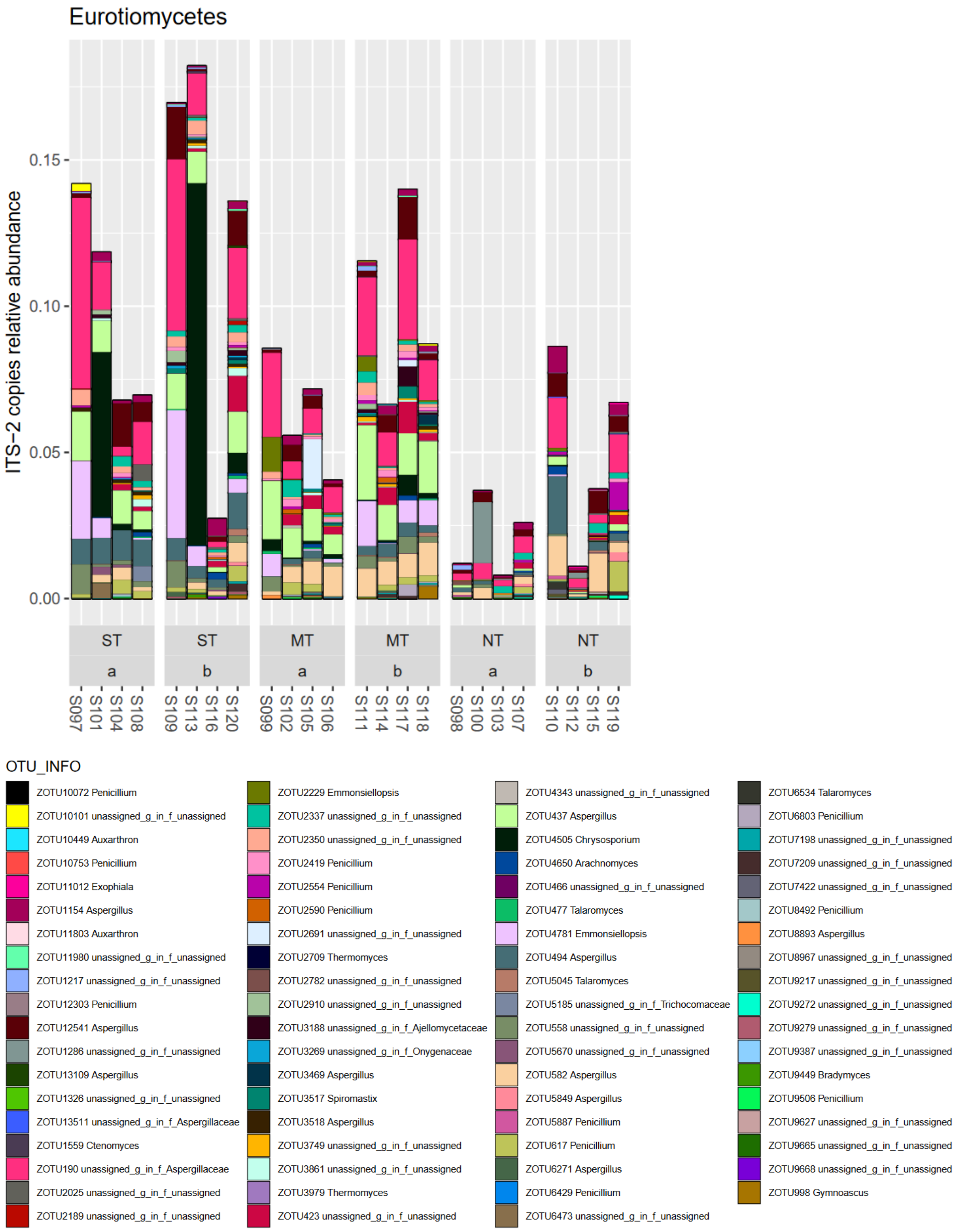


**Figure A42: Class Eurotiomycetes of phylum Ascomycota. This contains well-known genera such as *Penicillium* and *Aspergillus*.**

In the unassigned Ascomycota, 6 taxa were significantly associated with tillage management factor. Of those, 4 were in clusters with increases under NT (or vice versa increases under ST), and two were in clusters with relative decrease under NT.

Many unknown/unassigned taxa, such as free-living, dark endophytic, saprobic or soil unicellular organisms for example Archaeorhyzomycetes are to be found in Ascomycetes (Taylor and Sinsabaugh, 2015).

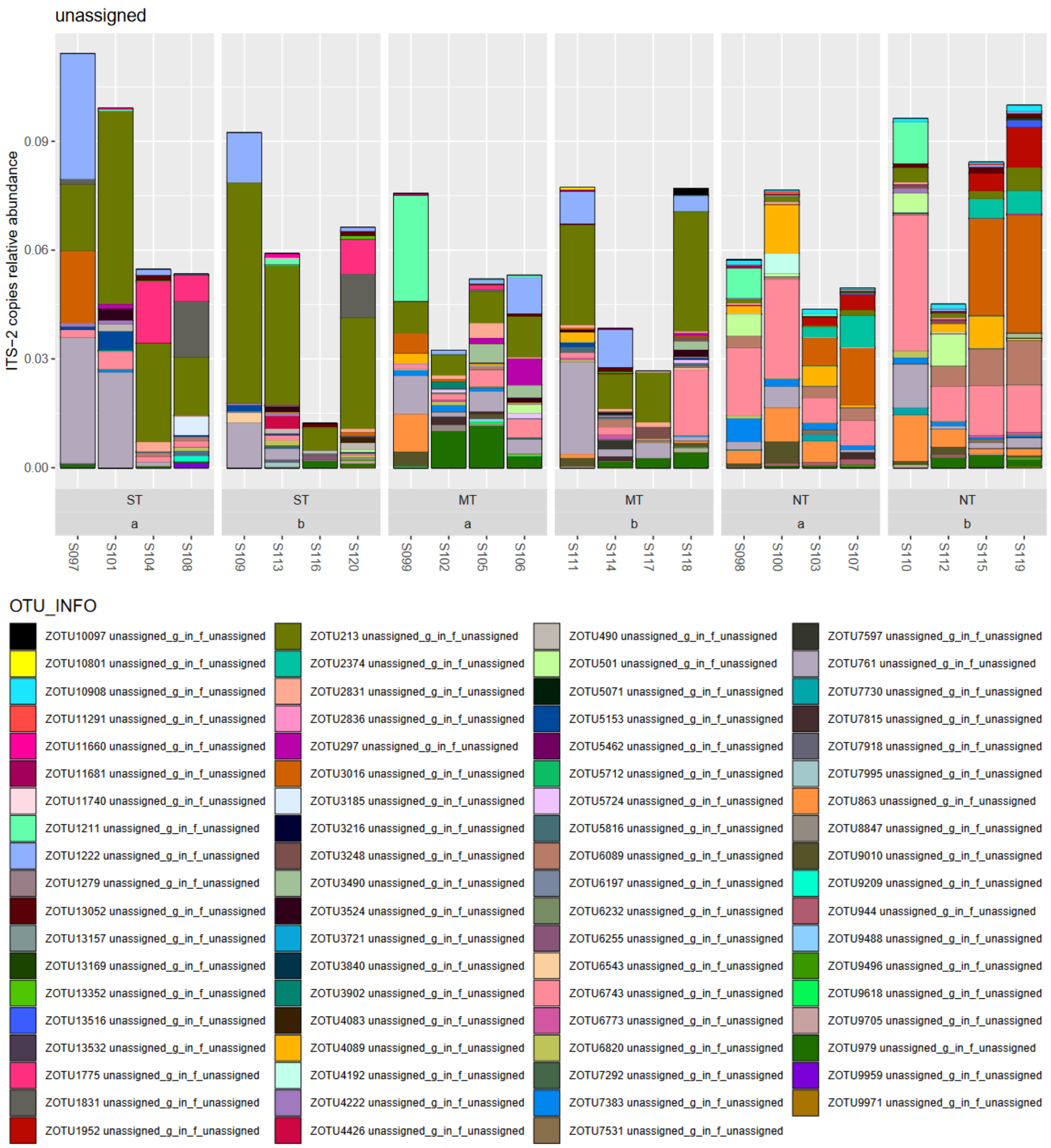


**Figure A43: Ascomycota unassigned on class level**

In the class Pezizomycetes, three taxa were in clusters associated with increases under NT management and one taxon was in a cluster with strong decreases under NT vs ST in the top layer. In the former group, taxa were assigned to the genus *Otidea* (family *Pyronemataceae*, order Pezizales), while the other two taxa were from the family *Ascobolaceae* (order Pezizales) and unassigned up to class. In the latter group, the taxon with strong decreases under NT vs ST in the top layer was from the genus *Ascobolus* (family *Ascobolaceae*, order Pezizales).

Members of the cup-fungus family *Pyronemataceae* are ecologically diverse. The majority is considered as saprotrophic but they also include ectomycorrhizal, bryosymbiotic and parasitic species such as terricolous, coprophilous, lignicolous, pyrophilous, urinophilic, and bryophilous members (Perry et al., 2007, Hansen et al., 2013).

*Otidea* are noted for their diversity of resin-like exudates. Their delineating feature within the *Pyronemataceae* family are the ear-shaped apothecia (Olariaga et al., 2015).

*Ascobolaceae* also form cups as fruiting bodies, the ascocarp, which are determining features for its genera (van Brummelen, 1972). The genera are: *Ascobolus*, *Saccobolus*, and *Thecotheus*. Most are coprophilic and some are also found on other organic substrates and soil. All organisms in this family are presumed to be saprobic/saptrotrophic (Hansen and Pfister, 2006). The genus *Ascobolus*, is widely reported to contain taxa with mostly coprophilic/dung degrading nutrition (Doveri, 2014).

Going back to the class level. In the Pezizomycetes, lychenization (symbiotic relation with photobiont such as Cyanobacteria or Algae) is widespread, with ca. 50% of species entering this relation. The minority of these, ca. 8-10%, enter a relation with Cyanobacteria (Lücking et al., 2009, Wedin et al., 2009).

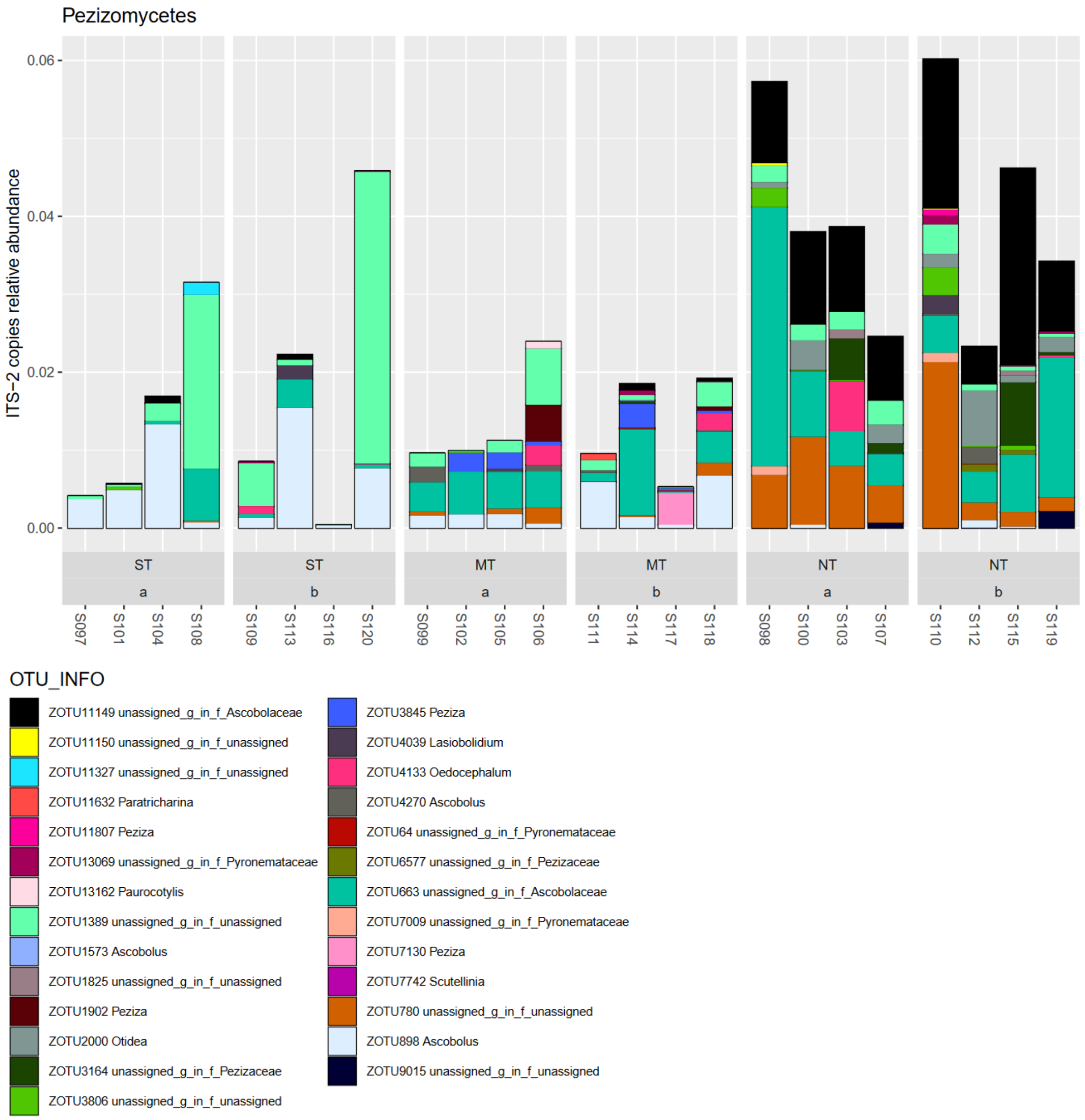


**Figure A44: Class Pezizomycetes in Ascomycota.**

In the class Orbiliomycetes, one taxon was significantly increased under NT vs ST in lower soil depth. The taxon is from the family *Orbiliaceae* (order Orbiliales).

The dataset contained two taxa from the genus *Arthrobotrys* (family *Orbiliaceae*, order Orbiliales), but without significant changes from tillage treatment (p<5%). Generally, the counts for taxa in this class already were sparser.

This class may contain nematode trapping fungi from the genera ***Arthrobotrys*** and *Drehslerella* (Zhang et al., 2023).

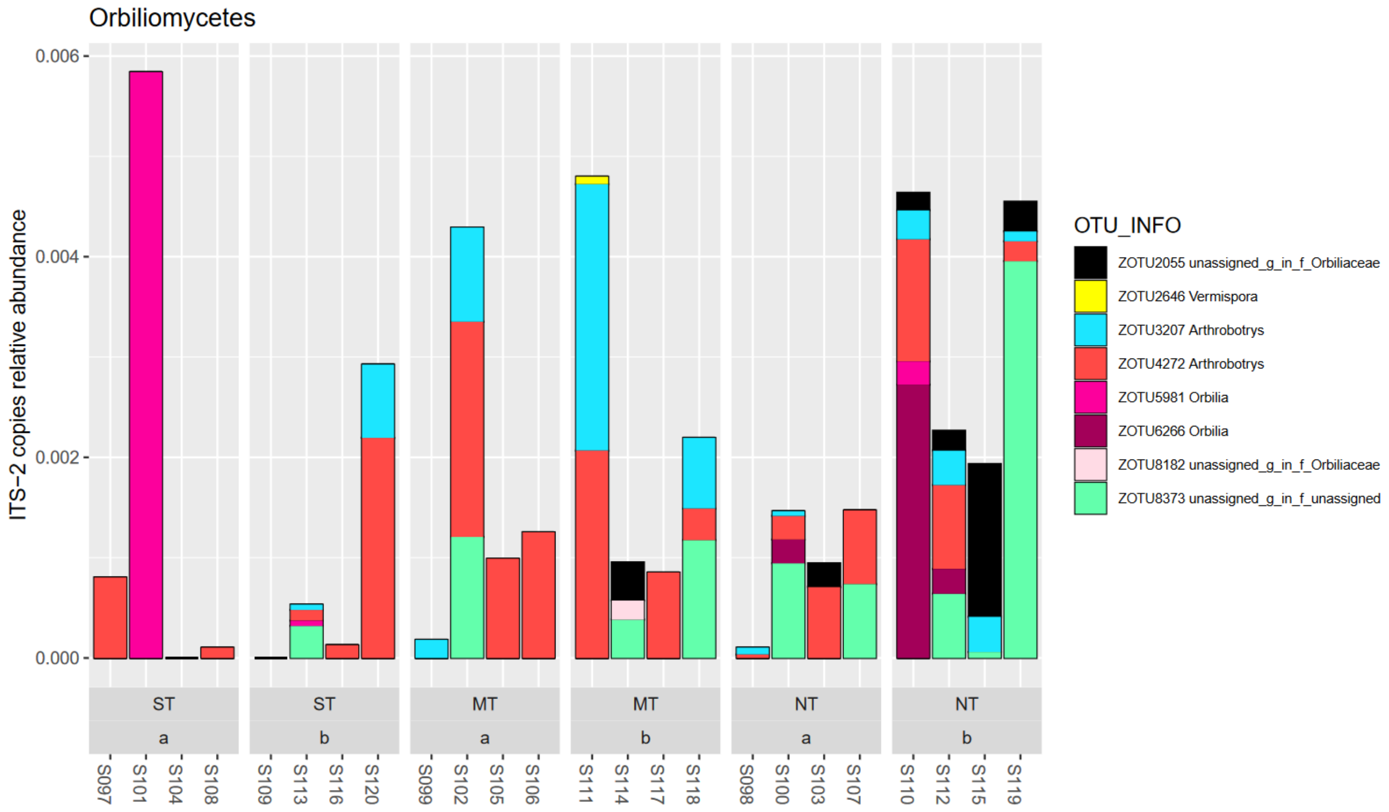


**Figure A45: Class Orbiliomycetes in Ascomycota**

Saccharomycetes, Lecanoromycetes, Taphrinomycetes, Laboulbeniomycetes, Arthoniomycetes, and Pezizomycotina class *incertae sedis* are not shown due to sparse counts.

Within the Lecanoromycetes taxa can be found that are “lycheniform” symbiont partners with cyanobacteria (see Cyanobacteria section, significant strong increase under NT). Taxonomies with cyanobacterial partner are in *Arctomiaceae* (*Ostropomycetideae*), Lichinales (Lichinomycetes) and Peltigerales (*Peltigerineae* or within Lecanorales). The fungi with *Nostoc* cyanobacteria as partner are from the *Collemataceae* family and are characteristically swollen such that the lichens are described as gelatinous lichens. The non-gelatinous fungi are from the *Pannariaceae* (Wedin et al., 2009).

In this dataset organisms from Lecanoromycetes were rare:

- Order Teloschistales family *Teloschistaceae* (0.08% in ST a)
- *Rinodina pyrina*, order Caliciales, family *Physciaceae* (0.04% in MT a)
- Order Lecanorales family *Lecanoraceae* (0.02% in MT a)

##### 1.3.2 Basidiomycota – also diverse ecology

The phylum Basidiomycota contained the second most ITS2 counts in the dataset, at on average 25%. Particularly in NT management topsoil, this phylum was very relatively abundant at 46%. This phylum with its richness of taxa ($n_{total}$= 208 OTUs, 5 taxa with significant changes) reacted less frequently to tillage treatments than the Ascomycota ($n_{total}$= 519 OTUs, 33 taxa with significant changes) (LinDA, FDR<5%). Taxa with significant changes in Basidiomycota were all but one from the class Agaricomycetes, and the other taxon from the class Tremellomycetes. All had at least genus level assignment.

In this dataset, the following classes of Agaricomycetes were found in order of decreasing total relative abundance: Agaricomycetes, Tremellomycetes, Exobasidiomycetes, unassigned Basidiomycota, and Geminibasidiomycetes. Further classes with sparse observations were taxa assigned to Cystobasidiomycetes, Microbotryomycetes, Agaricostilbomycetes, Wallemiomycetes, Ustilaginomycetes, and Pucciniomycetes.

Basidiomycota are a large clade of fungi with over 30,000 described species. It contains organisms visibly recognizable through their mushrooms such as *Agaricus* or *Amanita*. There are also well-known pathogens in this group such as smuts and rusts of grains such as *Ustilago* and *Puccinia* (Madigan et al., 2021).

In the class Agaricomycetes, four taxa associated to clusters with increases under NT management were indicated. Foremost, due to its high relative abundance and strong increases under NT management was a taxon with assignment to *Coprinellus curtus* (family *Psathyrellaceae*, order Agaricales). It increased in relative abundance of ITS2 counts to 20.5% under NT management in the top soil layer. Another taxon with significant increases from the same family was a *Psathyrella* at a much lower relative abundance of 0.2% in NT lower soil depth. Further taxa with significant relative increases, but on a similarly low relative abundance level were from the genera *Conocybe* (family *Bolbitiaceae*, order Agaricales) and a *Crepidotus* (family *Crepidotaceae*, order Agaricales).

Genera of note in the dataset were taxa belonging to *Psathyrella*, which is widely regarded as the archetypical little brown saprotrophic mushroom. It has to be noted that the family *Psathyrellaceae* was found to be polyphyletic, but phylogenetically closely clustered groups are *Coprinopsis*, and *Coprinellus (formerly Coprinus,* Redhead et al., 2001*)*. Generally, fungi in this class are saprotrophic. The former, *Coprinopsis*, is typically (but not exclusively) coprophagous, giving it its name (Padamsee et al., 2008). The latter genus, *Coprinellus* was found with several taxa in the dataset.

Especially one taxon increased strongly under NT management with assignment to the species *Coprinellus curtus*. *Coprinellus curtus* grows well in mycelia in the dark on malt extract agar, as others in this family, but has prominent rust stained cups (Badalyan et al., 2011). The fruiting bodies are short lived above ground, and this may lead to underestimation of its presence in soil – in Japan the common name is Hitoyo-take translated as “overnight fungus” (Nakasaki et al., 2007). The strain of *Coprinellus curtus*, isolated from vegetable plantations with organic amendments, had antagonistic behavior (direct hyphal interference) against several plant pathogens such as *Rhizoctonia solani* (Basidiomycota) that cause bottom rot disease on lettuce and Rhizoctonia-patch disease on Mascarene grass, and against *Fusarium oxysporum* (Ascomycota) that cause crown (foot) and root-rot disease of tomato and wilt of melon (Nakasaki et al., 2007).  A further example in this dataset is *Coprinellus xanthothrix*, which was isolated as a coprophil and grew well on glucose and peptone. It was found to have nematicidal activity (Liu and Bau, 2023, Liu et al., 2008).

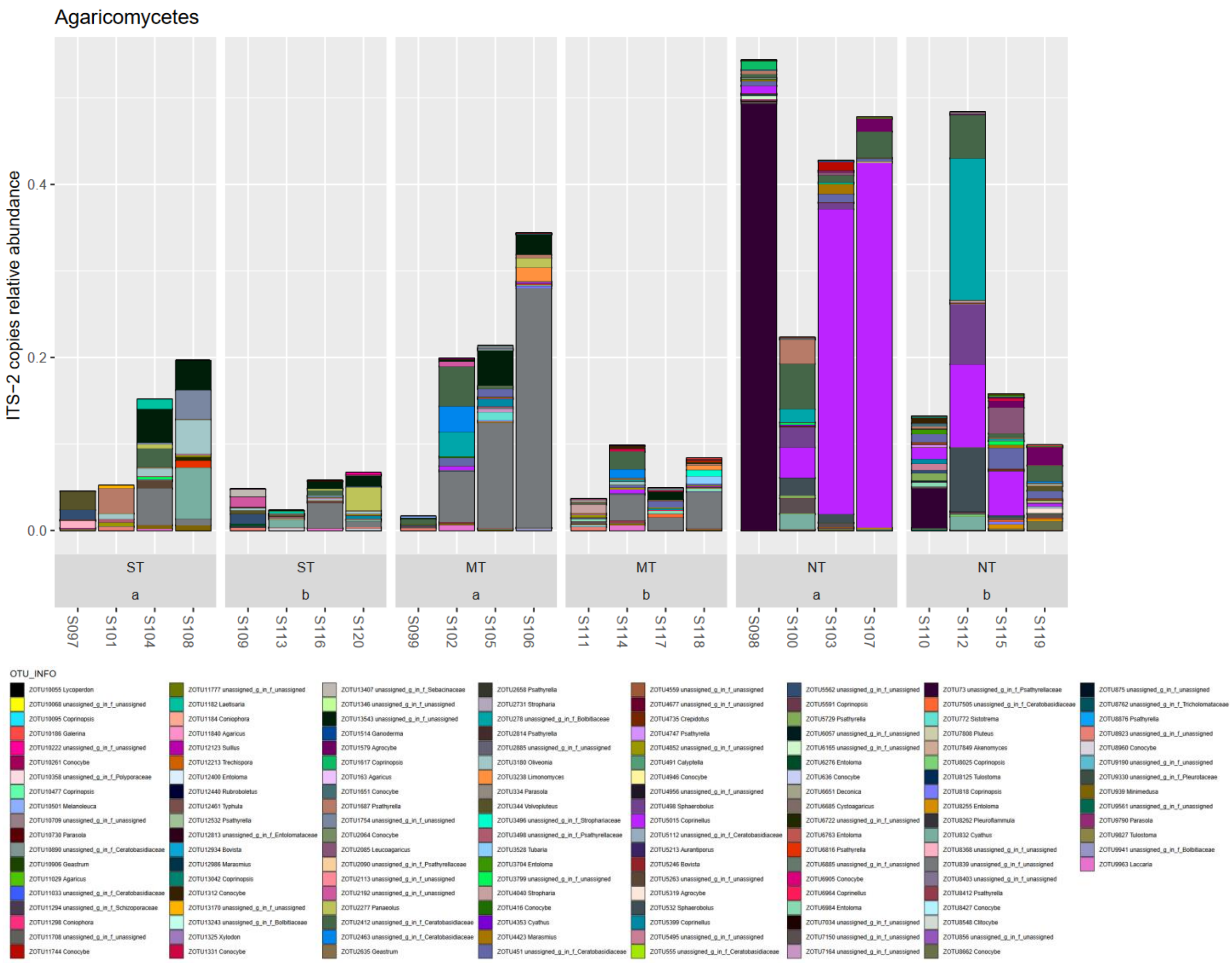


**Figure A46: Class Agaricomycetes in Basidiomycota.**

The class Tremellomycetes contained one taxon with significant changes to the tillage treatments. This was a taxon from the genus *Udeniomyces* (family *Mrakiaceae*, order Cystofilobasidiales) with significant decrease under NT vs ST (vice versa, increase under ST vs NT) in top soil layer. It had a relative abundance of 0.4% in ST top soil layer.

The Tremellomycetes are a polyphyletic grouping encompassing yeasts, dimorphic taxa and species that form hyphae and/or complex fruiting bodies (Liu et al., 2015).

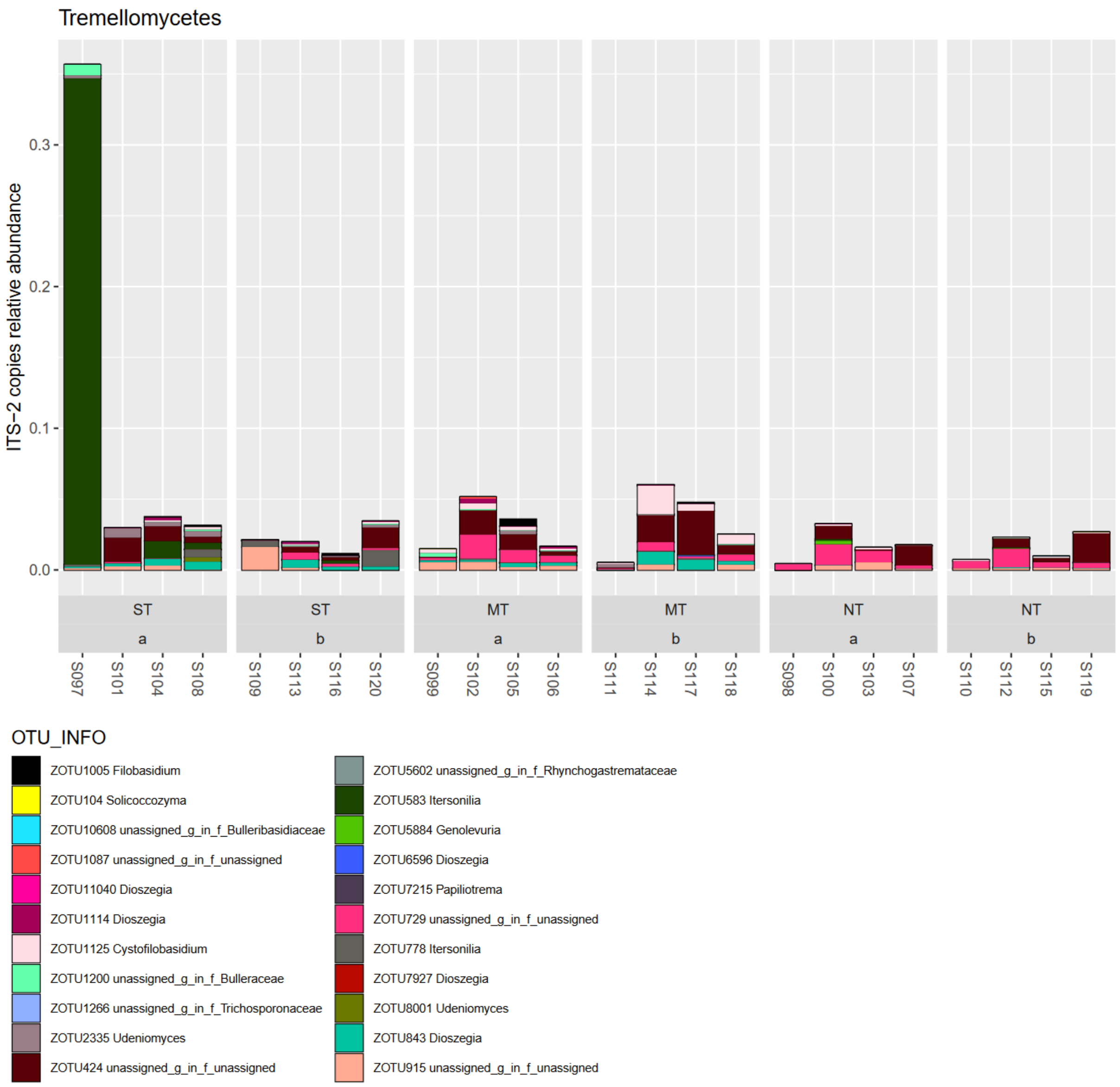


**Figure A47: Class Tremellomycetes in Basidiomycota**

No significant changing taxa were found in this class

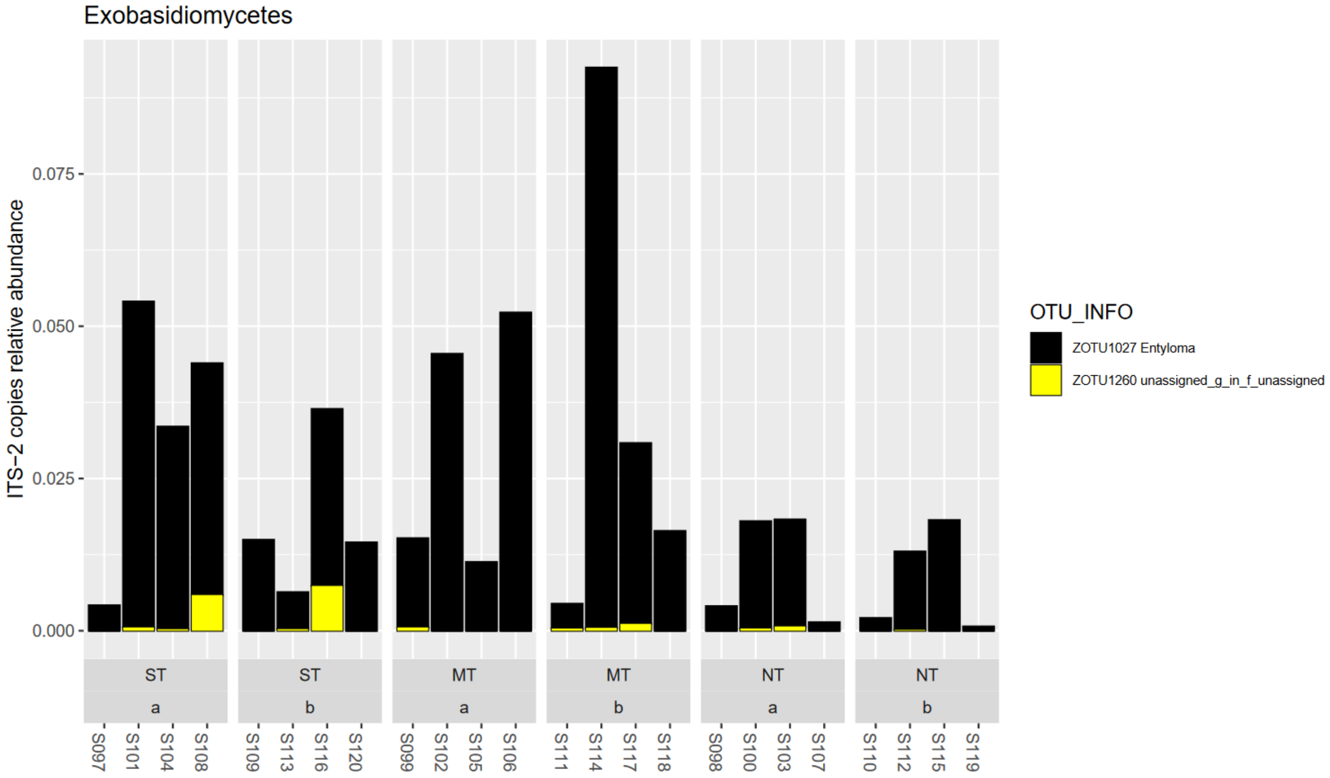


**Figure A48: Class Exobasidiomycetes in phylum Basidiomycota**

No significant changing taxa were found in this class

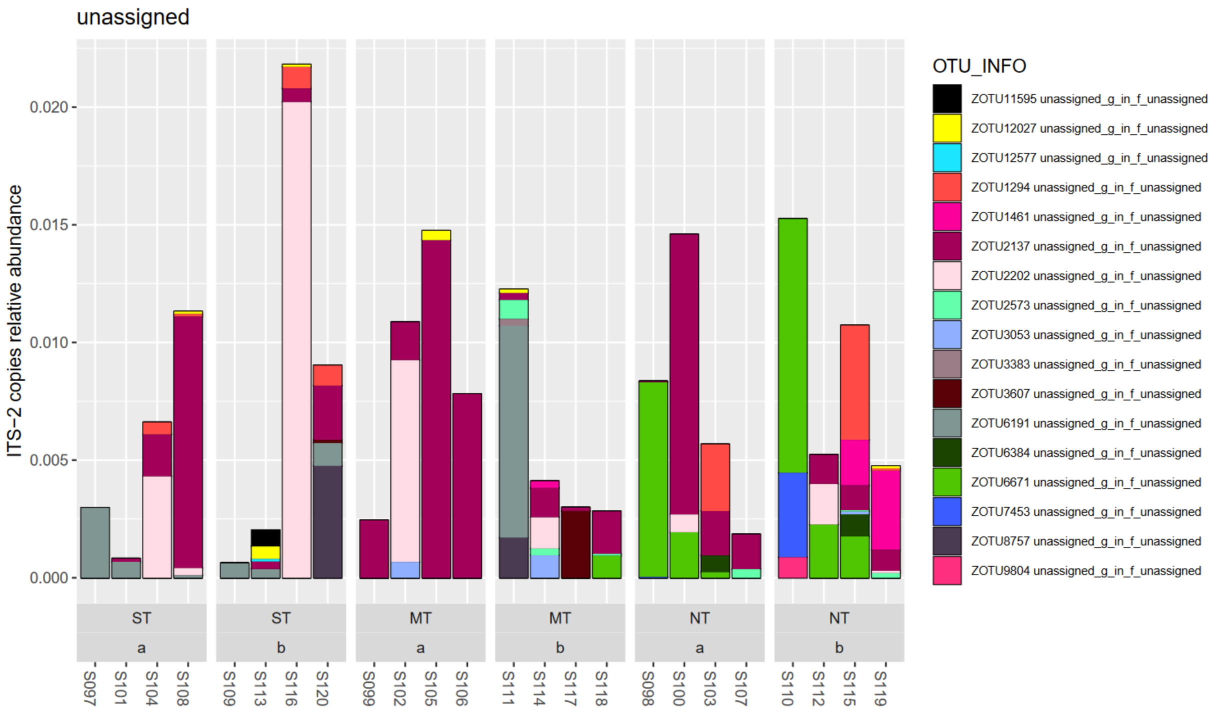


**Figure A49: Class unassigned in phylum Basidiomycota**

No significant changing taxa were found in this class

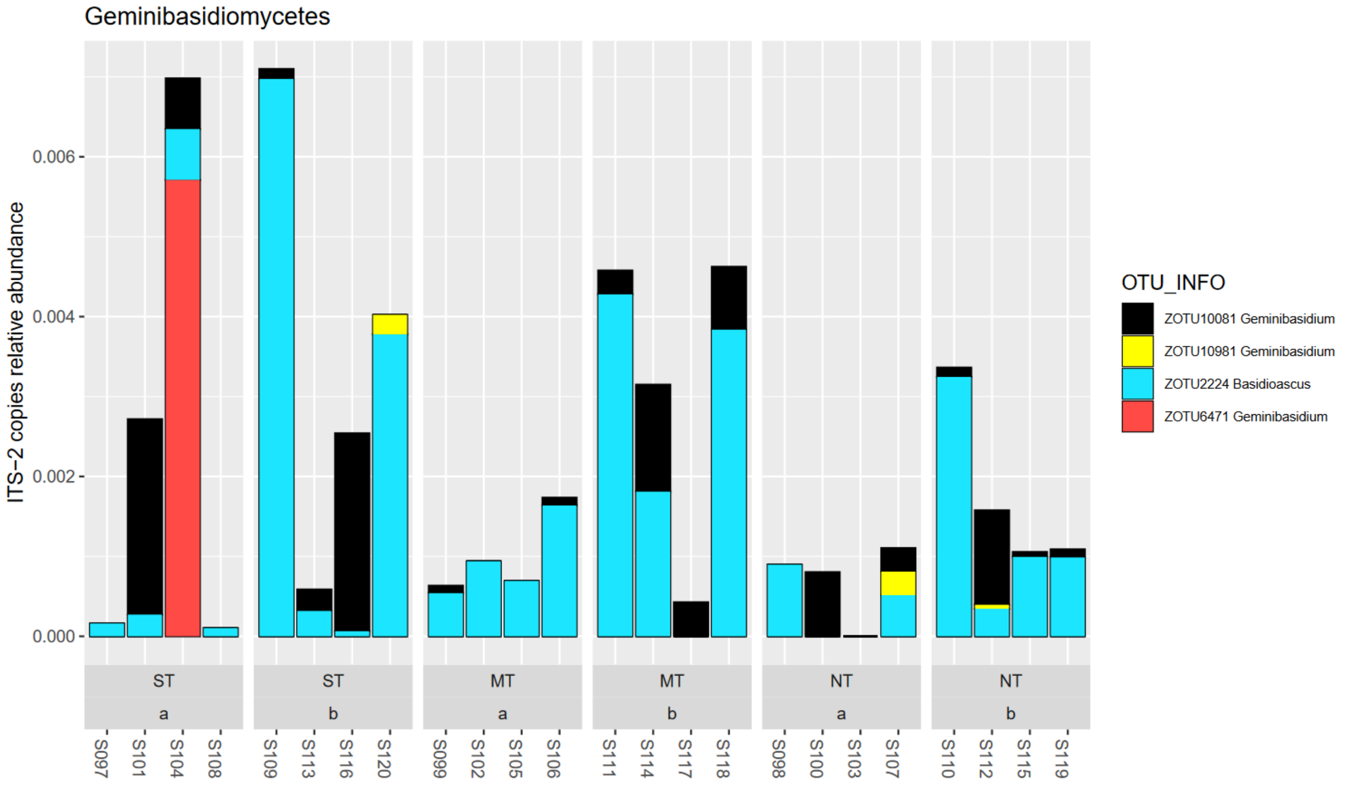


**Figure A50: Class Geminibasidiomycetes in phylum Basidiomycota**

##### 1.3.3 Glomeromycota

In the dataset, sequences belonging to Glomeromycota represented on average 3% of total reads. In lower depth (10-20 cm) the average was higher than in top soil layer, at 4%.

Three taxa in this phylum, which were from the richest class Glomeromycetes, were significantly changing with tillage treatments (LinDA FDR<5%). Two taxa from the family *Glomeraceae* (order Glomerales) were associated with a cluster with moderate increases under NT in lower soil depth. The most relatively abundant of these, at 0.7% was of a magnitude lower abundance than the most abundant taxon in this phylum.

Another taxon, unassigned up to class Glomeromycetes, with significant decreases in NT vs MT in lower soil depth layer was also not highly abundant in terms of ITS2 copies, at 0.1%

Generally, the phylum contains almost the entirety of known Arbuscular Mycorrhizal fungal (AMF) species. The family *Glomeraceae* contains most species such as *Glomus*, *Funneliformis*, *Sclerocystis*, *Rhizoglomus*, *Septoglomus*, *Simiglomus*, *Dominikia*, *Kamienskia*, *Halonatospora*, *Oehlia*, *Funneliglomus*, *Nanoglomus*, *Microdominikia*, *Orientoglomus*, *Sclerocarpum*, *Microkamienskia*, *Epigeocarpum*, *Silvaspora*, *Complexispora*, and *Blazkowskia* (Sieverding et al., 2014, da Silva et al., 2023).

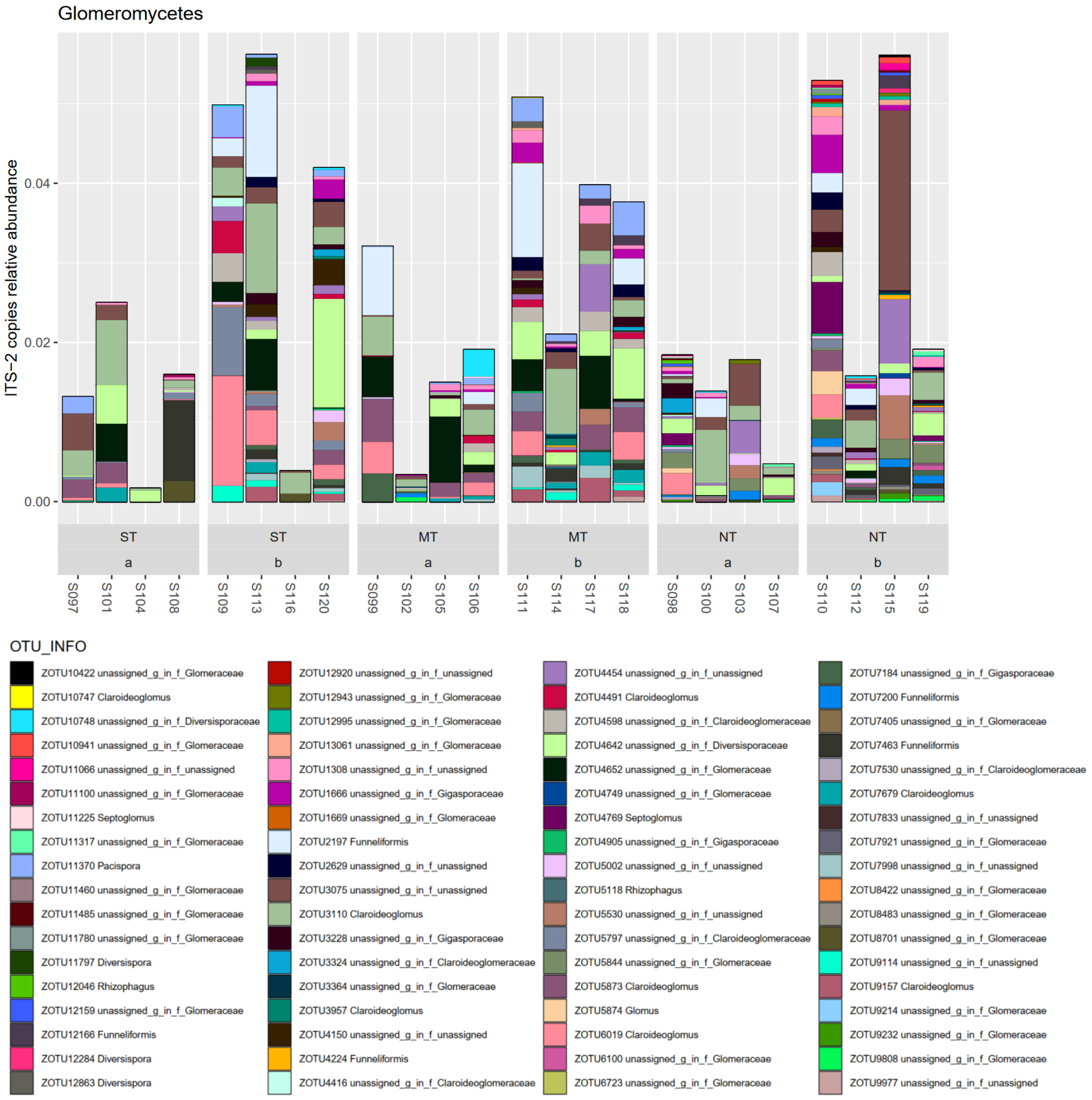


**Figure A51: Class Glomeromycetes in phylum Glomeromycota**

No significantly changing taxa (p>5%) were found in the class Archaeosporomycetes, likely also due to low counts/sparsity.

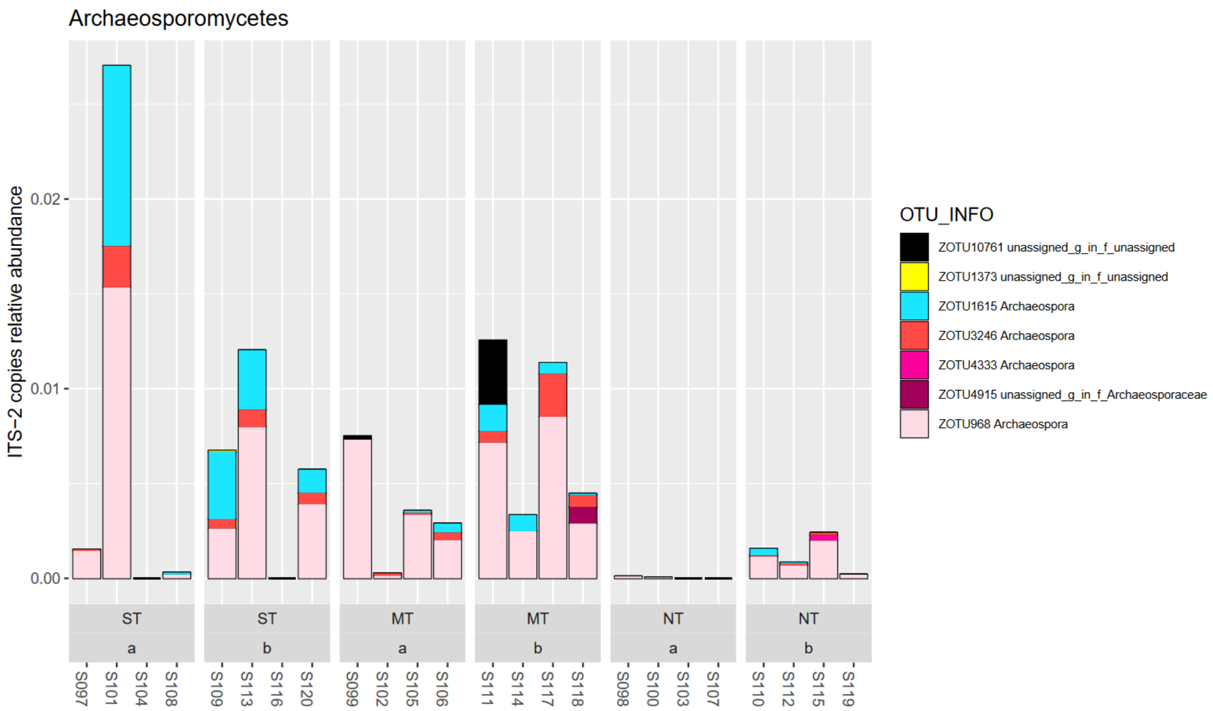


**Figure A52: Class Archaeosporomycetes in phylum Glomeromycota**

##### I.3.4 Chytridiomycota

The Chytridiomycota were the third most relatively abundant fungal phylum in the dataset and 84 taxa were clustered at 97% similarity of the ITS2 sequence. Here, the following classes in decreasing total relative abundance were found: Spizellomycetes, unassigned class, Rhizophydiomycetes, Chytridiomycetes, and Rhizophlyctidomycetes.

Of the 39 taxa assigned to the class Spizellomycetes, one taxon from the genus *Powellomyces* (family *Powellomycetaceae*, order Spizellomycetales) was moderately significantly increased under NT vs ST and MT in lower soil depth (LinDA FDR<5%). However, it was of very low relative abundance at 0.04%, compared to the most abundant taxon in this group (unassigned family) at 1.5% and the most relatively abundant *Powellomyces* in the dataset at 1.4% relative abundance.

*Powellomycetaceae* are soil-born basal fungi and a monophyletic clade within the order Spizellomycetales. Organisms from this family have been isolated from environments such as soils, plant detritus and manure by water solution and baiting with pollen (Simmons, 2011).

Several taxa from *Spizellomyces* genus were found in the dataset. Typical for this phylum are uniflagellar zoospores and in particular *Spizellomyces* are of importance in the soil environment through their range of beneficial-to-detrimental associations with mycorrhizal fungi, mildews, plants, and soil nematodes (Russ et al., 2016).

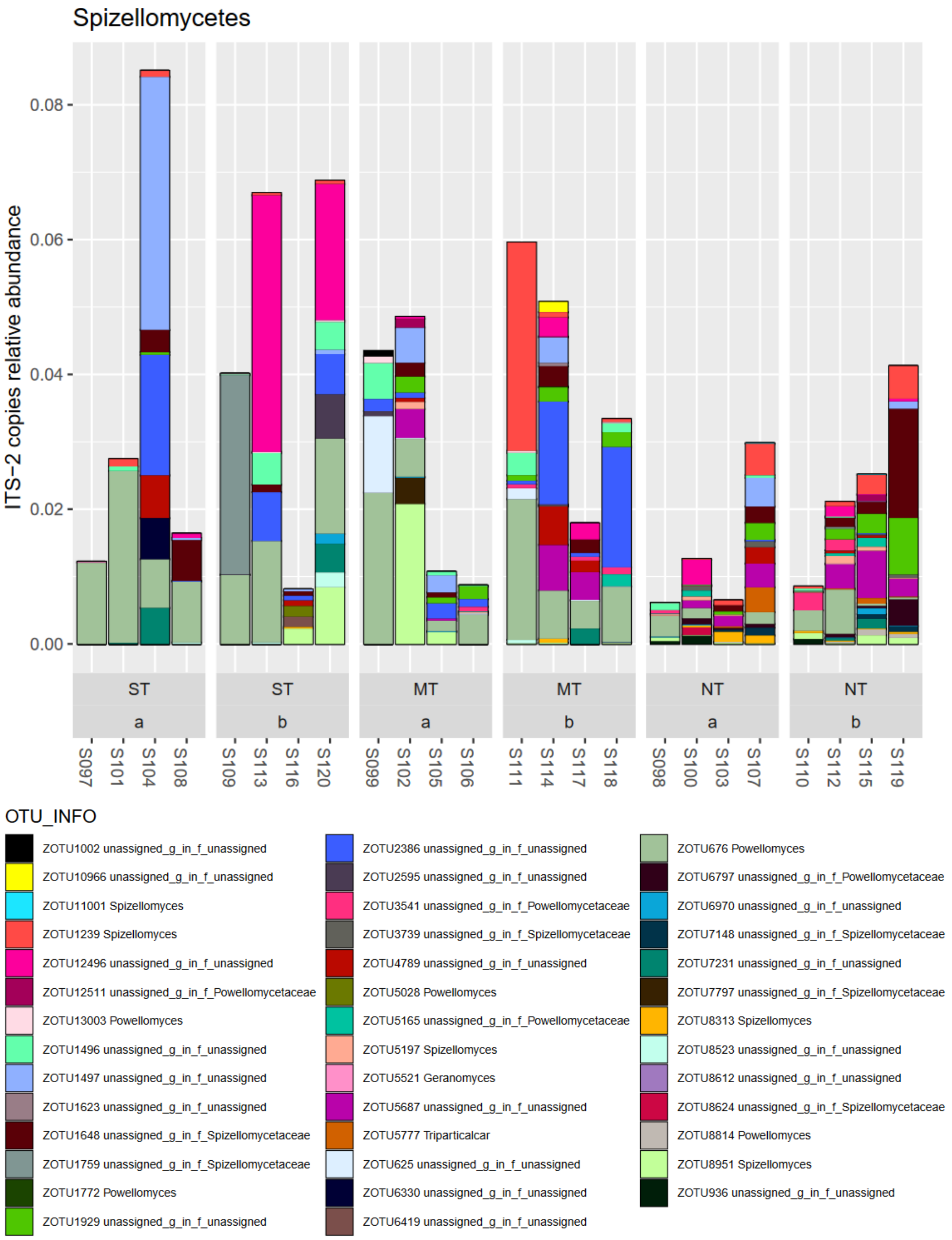


**Figure A53: Class Spizellomycetes in phylum Chytridiomycetes**

One taxon in the unassigned class of Chytridiomycetes was found to be significantly strongly increased in NT vs MT and ST in lower soil depth (p<5%). It was the fifth most relatively abundant taxon in this class at 0.5% in NT depth b.

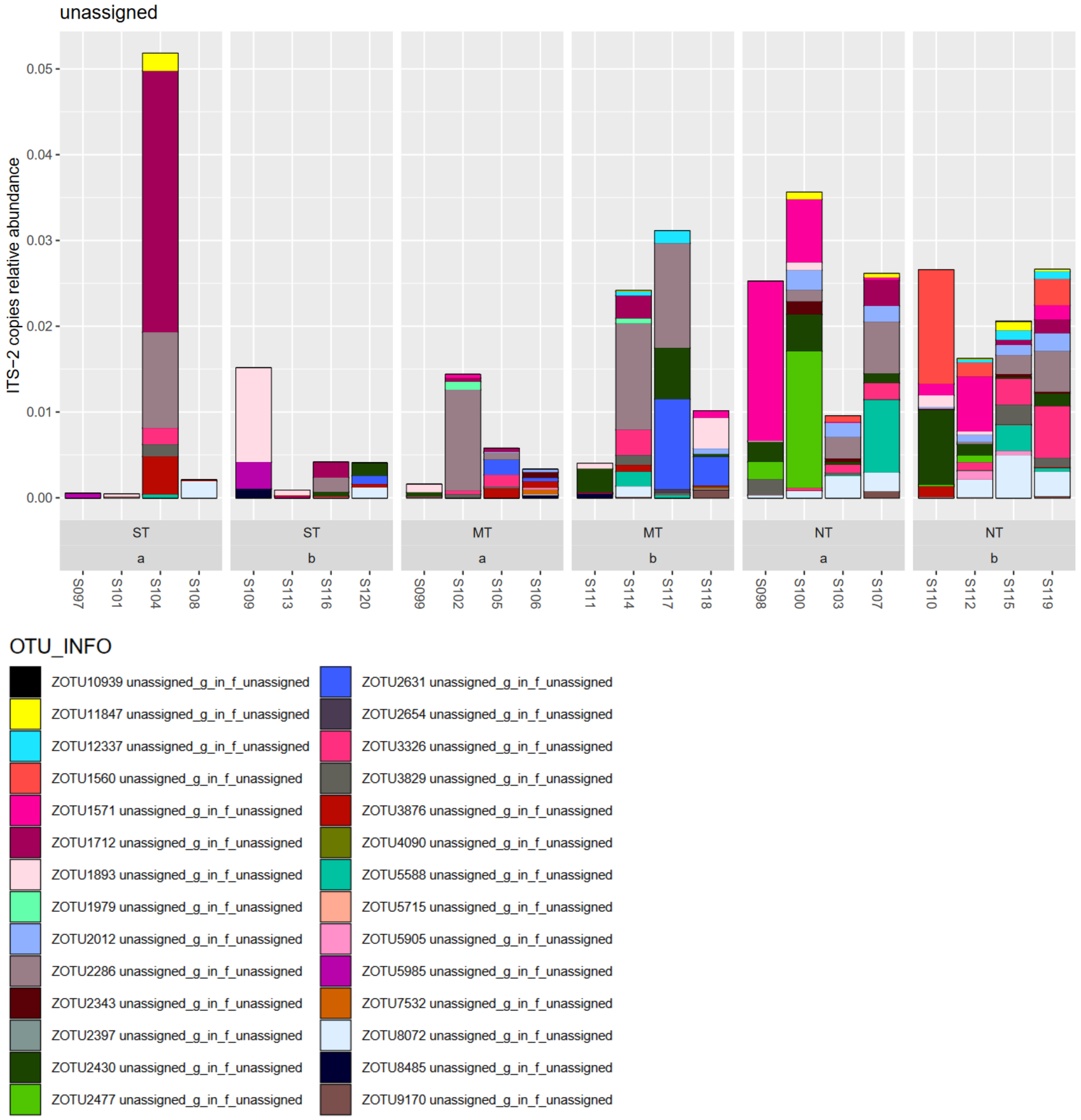


**Figure A54: Unassigned class in phylum Chytridiomycetes**

No significantly changing taxa were found in the class Rhizophydiomycetes (LinDA, FDR>5%), although some taxa appear to be more frequent under NT or MT, respectively.

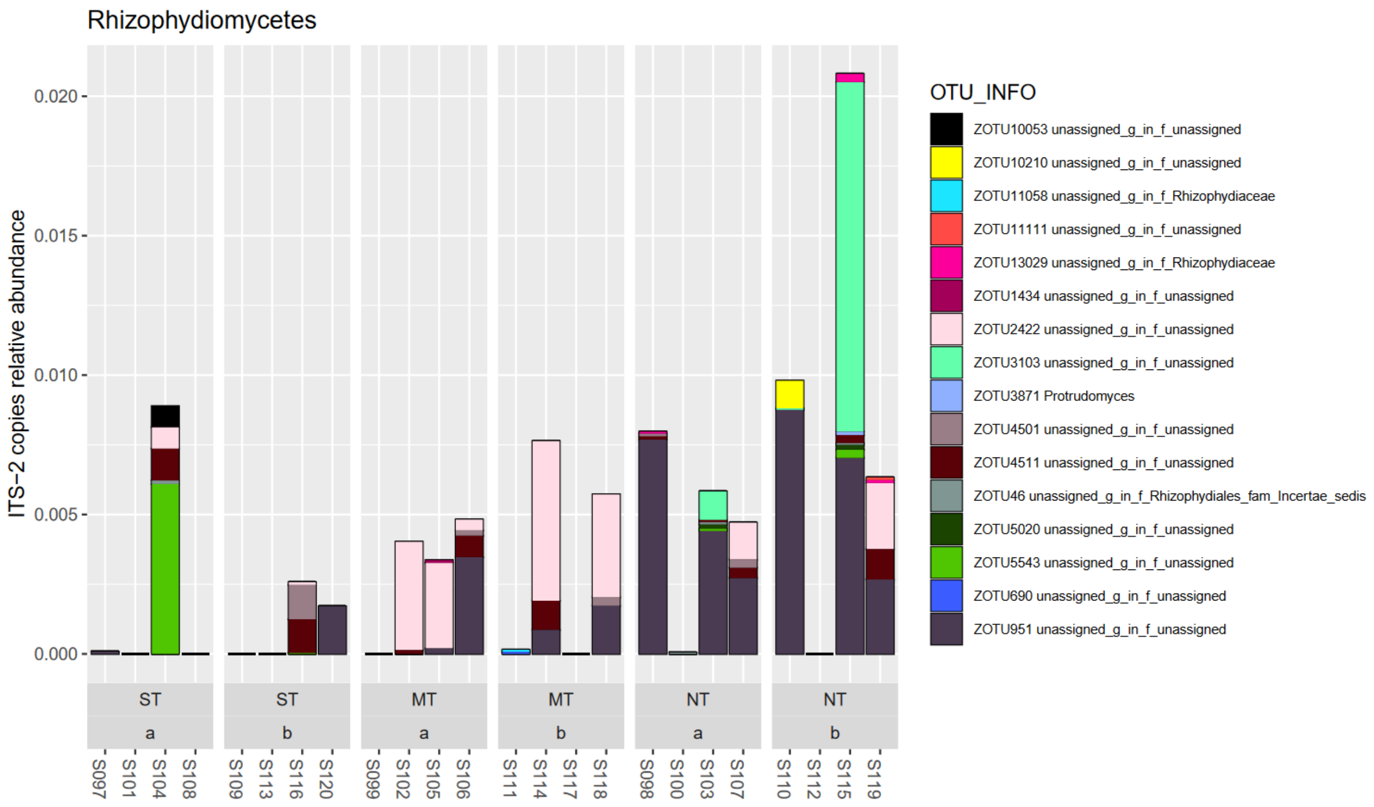


**Figure A55: Class Rhizophydiomycetes in phylum Chytridiomycetes**

No significantly different taxa were indicated in the class Chytridiomycetes (LinDA, FDR>5%), although the single taxon in this class appears more frequent in ST management.

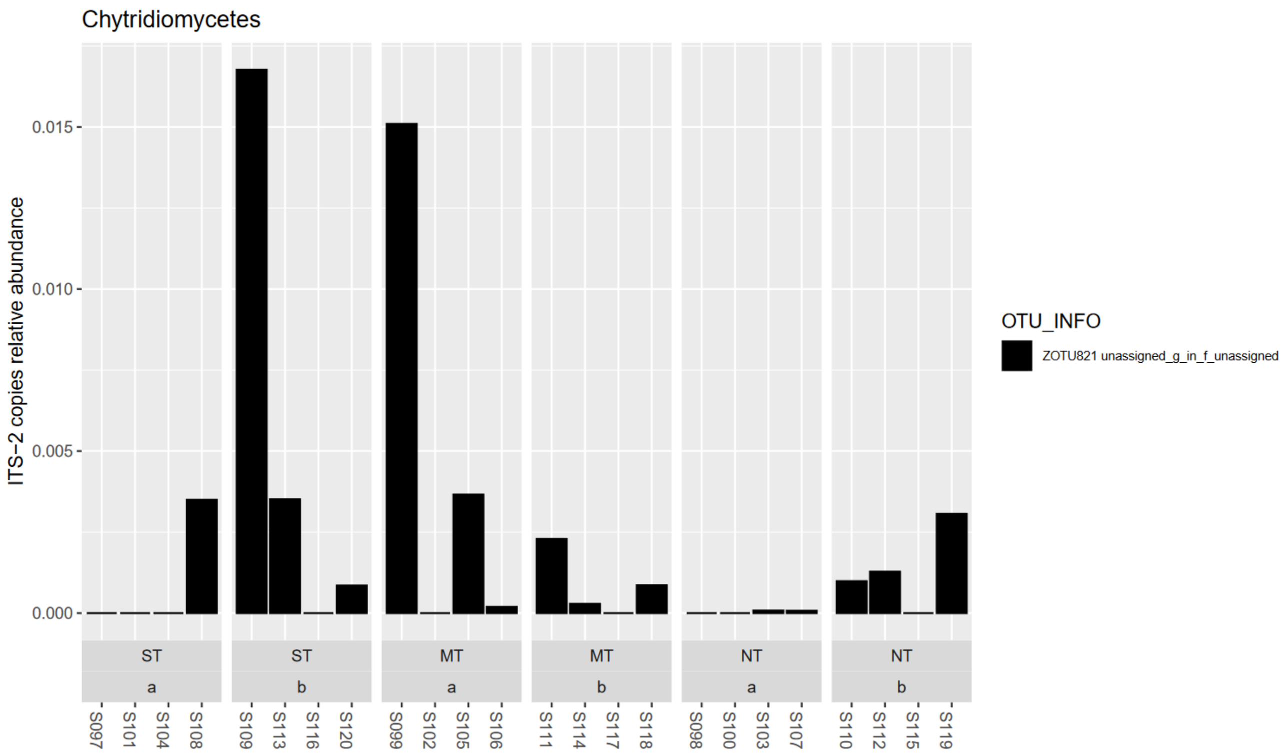


**Figure A56: Class Chytridiomycetes in phylum Chytridiomycota**

One taxon from the genus *Rhizophlyctis* (family *Rhizophlyctidaceae*, order Rhizophlyctidales) in the class Rhizophydiomycetes was found to be moderately significantly increased under NT vs ST and MT in lower soil depth (LinDA, FDR<5%).

The type organism of this genus can be cultured on yeast extract glucose chloramphenicol agar (CM+) and have been isolated from soil in many locations across the northern globe.

Depending on development stage, organisms attach to or penetrate plant pollen (Powell et al., 2015).

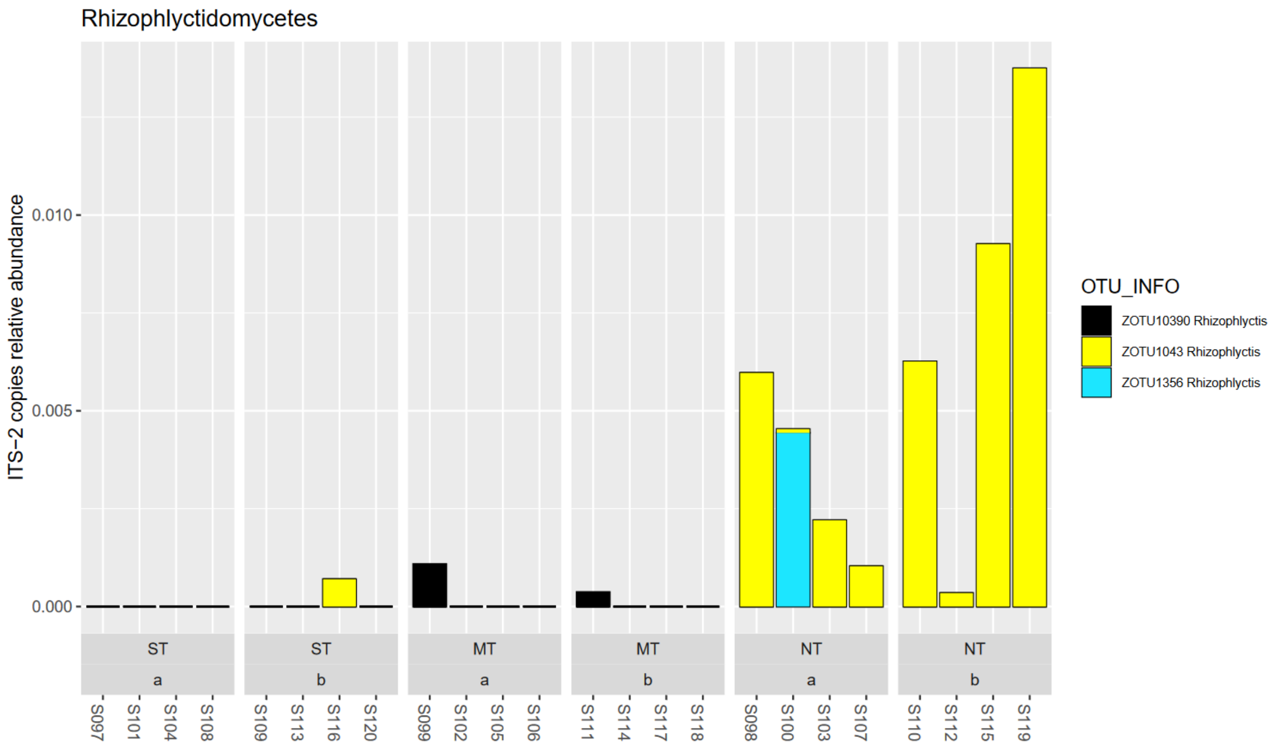


**Figure A57: Class Rhizophlyctidomycetes in phylum Chytridiomycota**

#### I.3.5 Morteriellomycota

In this dataset 26 taxa belonged to the class Morteriellomycota. Of those, 12 were assigned to the genus *Mortierella*. None of the taxa in this phylum were significantly associated to tillage management changes (LinDA, FDR>5%), although the total relative abundance appeared to be higher in ST management.

Phylogenetically Morteriellomycota are close to the Glomeromycota as they are classed under the sub-kingdom Mucoromyceta. Circa 100 species of *Mortierella* have been described, of which some taxa have been described that are beneficial to crops and mycorrhizal fungi in P acquisition, for example by synthesis and secretion of oxalic acid (Nguyen et al., 2019).

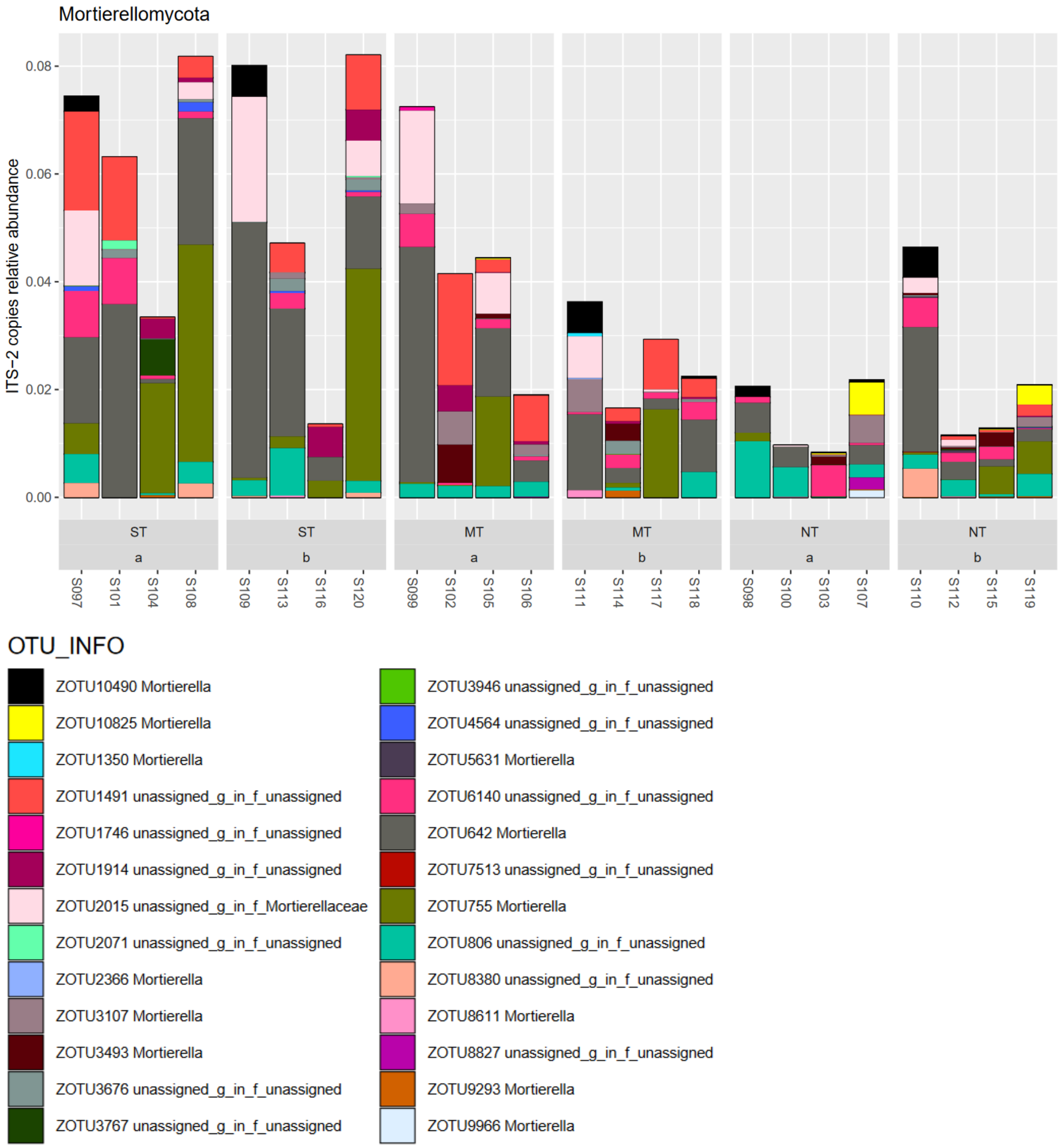


**Figure A58: Mortierellamycota phylum**

##### 1.3.6 Mucoromycota

In the dataset, 11 taxa were assigned to the phylum Mucoromycota. None were associated with tillage management changes (LinDA, FDR>5%).

Taxa from the genus *Mucor* were most relatively abundant, of which on species level assignments were made to *Mucor circinelloides, Mucor fragilis, Mucor hiemalis, Mucor moelleri*, *Mucor mucedo, Mucor plasmaticus,* and *Mucor racemosus*. This grouping of fungi contains very diverse ecologies from pioneers of organic matter degradation, rot and spoilage, food and beverage fermenters to soil born human pathogens (Walther et al., 2019).

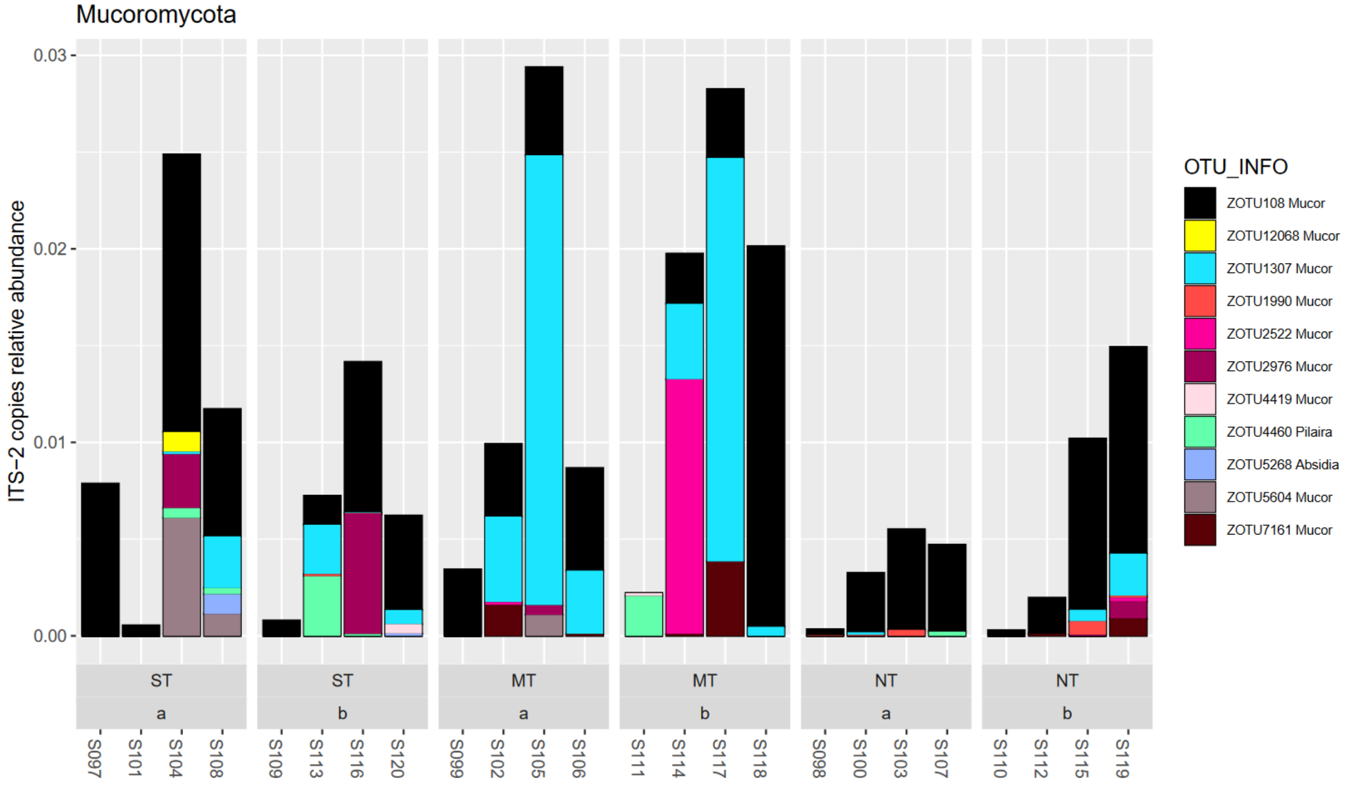


**Figure A59: Mucoromycota phylum**

##### I.3.7 Monblepharomycota

No significantly changing taxa were indicated in this relatively low abundance phylum consisting of one assigned taxon.

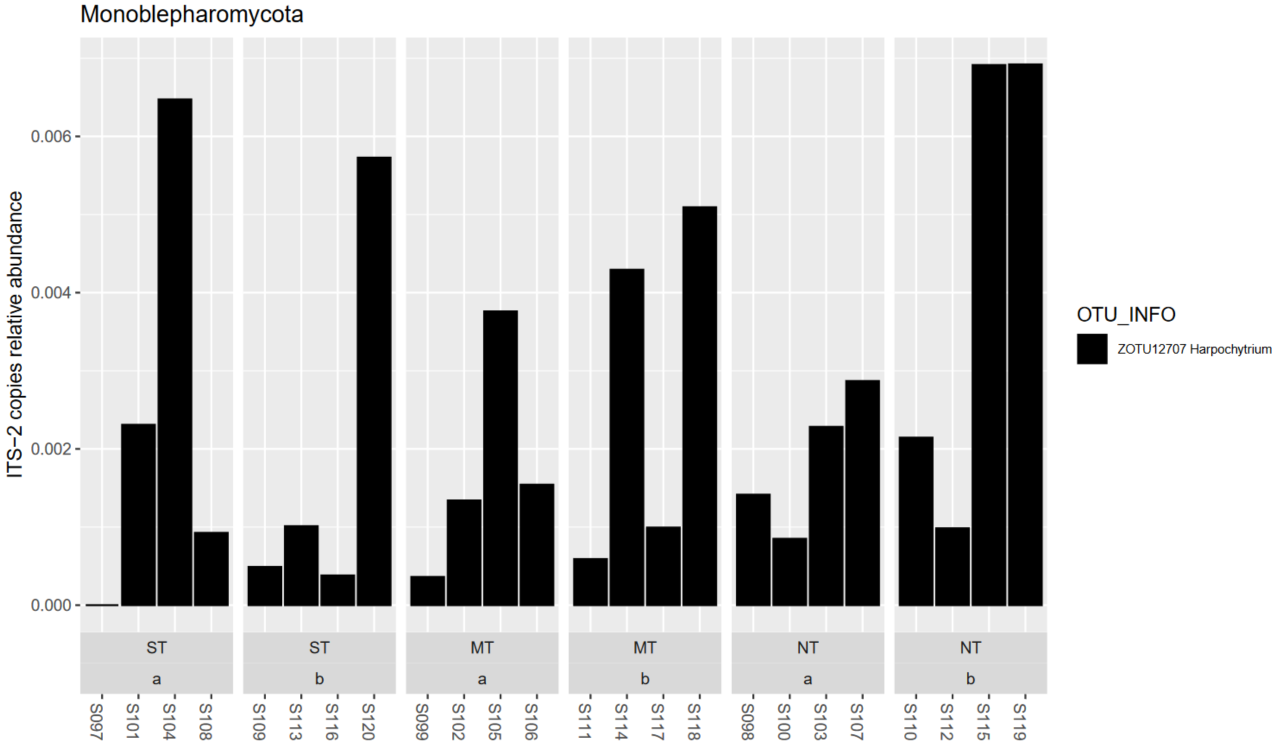


**Figure A60: Monoblepharomycota**

##### I.3.8 Rozellomycota

Several taxa were detected in this phylum, but in low relative abundance and none were indicated to be significantly differentially abundant.

The phylum Rozellomycota is placed in the basal fungal clade (as opposed to higher fungi such as Dikarya). Mostly, organisms in this grouping have been found to be parasites of aquatic animals, mites, insects, spiders, oligochaetes (segmented worms), amoebae, fungi and green algae (Wijayawardene et al., 2018).

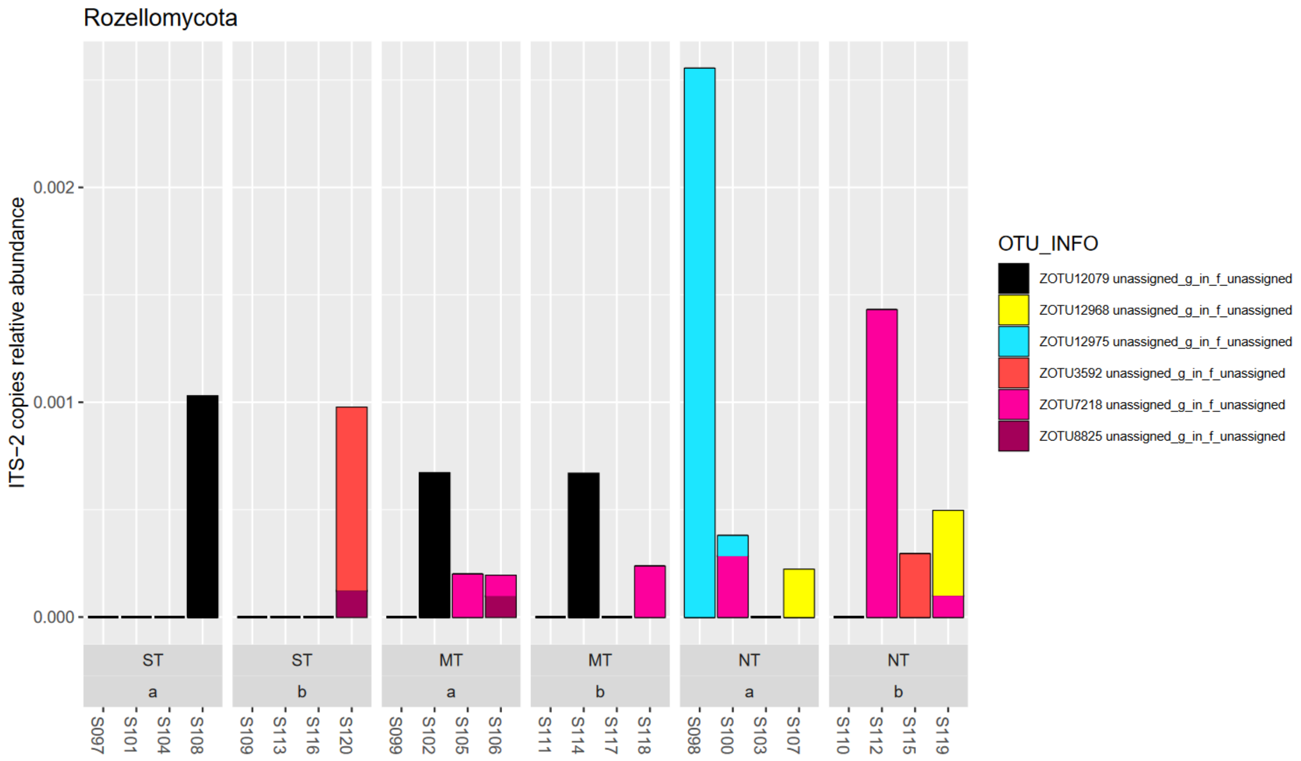


**Figure A61: Rozellomycota**

##### I.3.9 Unassigned fungi

In the dataset, 289 fungal taxonomic units without assignment up to phylum level were found. Of these, 7 were indicated to be significantly associated with tillage treatment factors. Four were associated with clusters with increase under NT and the others with clusters of decrease under NT (or vice versa increase under ST). The former group only contained taxa with low relative abundance < 0.05%, while the most abundant taxon in the latter group exhibited 1.3% relative abundance in ST top soil layer.

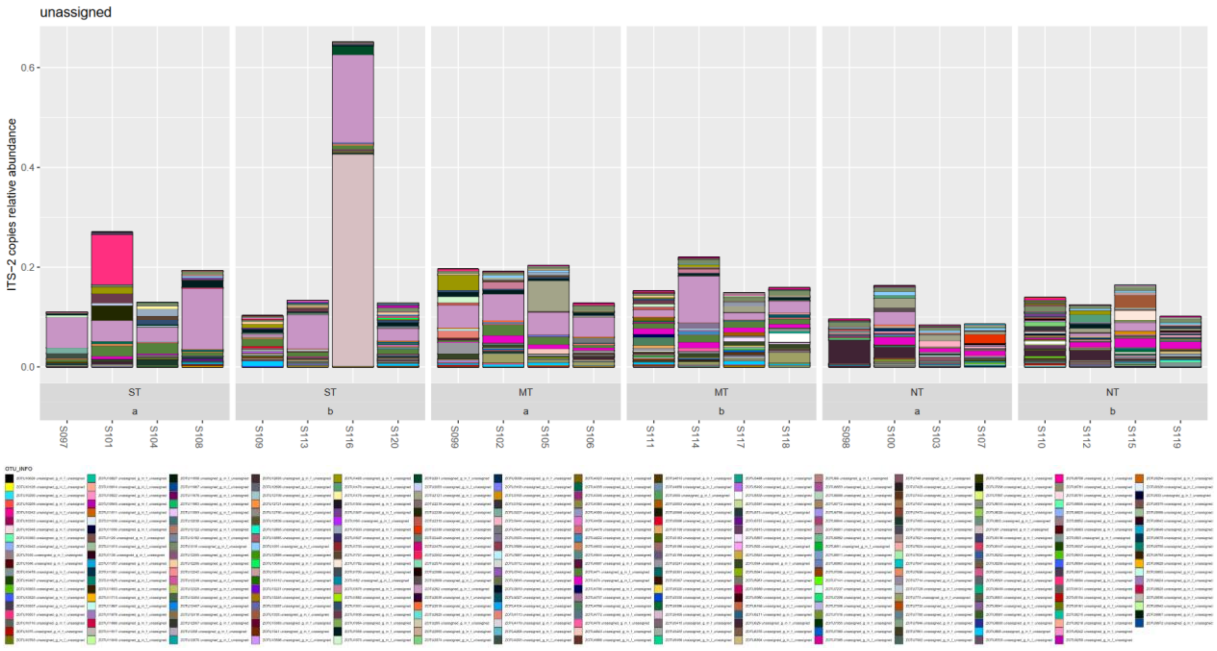


**Figure A62: Fungi with unassigned phylum**

#### 1.4 Differential abundance of fungi

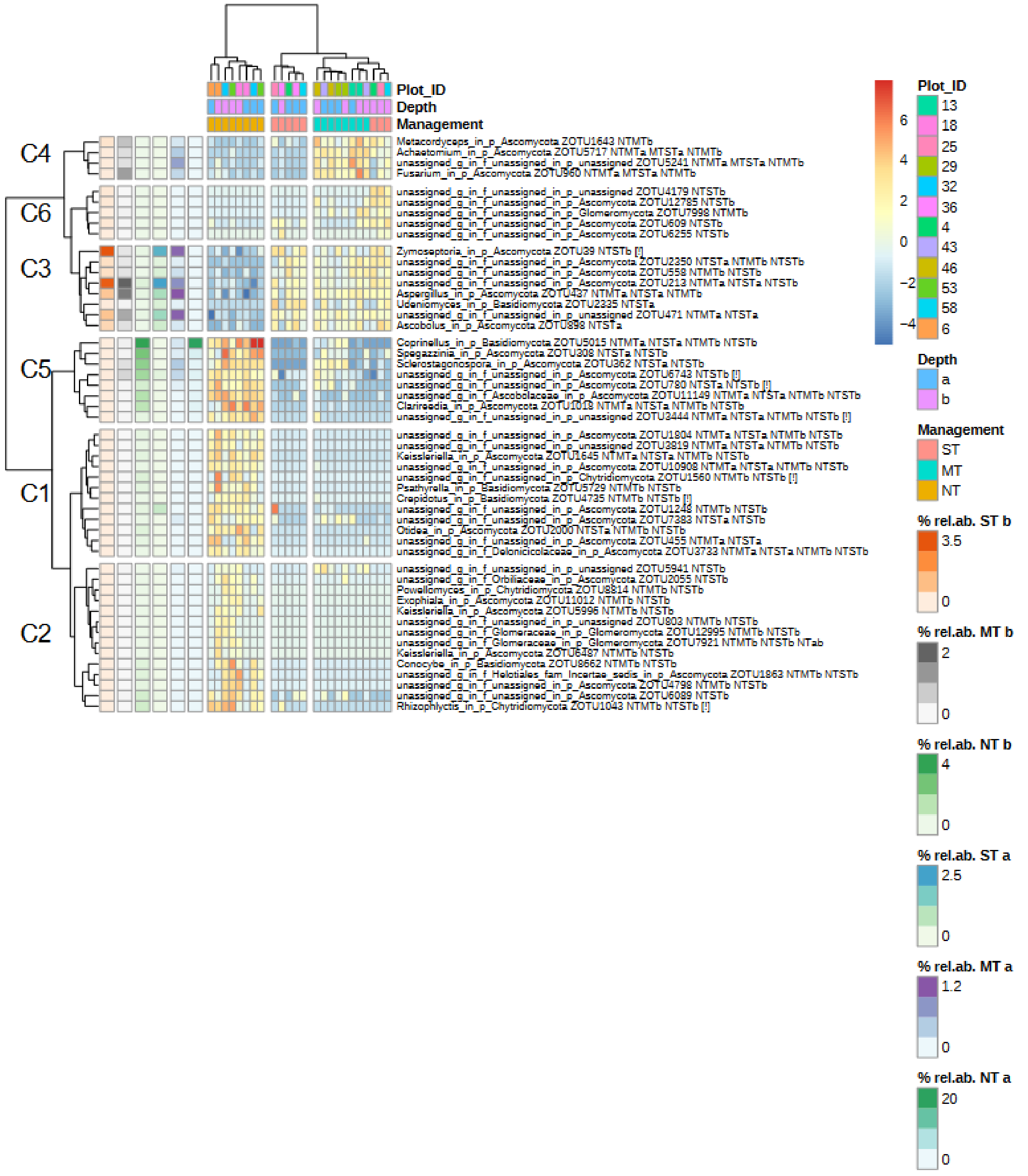


**Figure A63 Fungi, heatmap of log2-fold changes of taxon and sample against OTU-wise global mean. On the left average relative abundances of an OTU are resolved for treatment x depth combinations. Clustering of rows and columns was done with Euclidean distances and Ward`s clustering rule. Taxonomic information on OTUs is shown together with the contrast in which this taxon was indicated to be significantly differently abundant (e.g. contrast NT vs ST in depth a as NTSTa). Silhouette value was calculated, indicating the self-similarity of cluster members, and when negative indicated by [!] that the respective taxon could also be similar to a neighbour cluster. Clusters are indicated by "Ci" and correspond to numbering in Figure 2.**

#### 1.5 Biological soil crusts

Biological soil crusts are linkages of mineral soil particles with a community of photo-autotrophic cyanobacteria, algae, lichens, bryophytes, and associated heterotrophic fungi and prokaryotes that often develop in arid regions or on early habitats such as rocks. Heterotrophs may contribute to biocrusts with production of biofilm substances and pigments/resistance against UV radiation. In the arable context, the pioneer crusts developing from disturbance would be of more relevance than zonal crusts as climax communities limited by the respective biophysical conditions. In semi-arid and arid regions, the former are initially formed by cyanobacteria and develop more complexity with algae, lichens and bryophytes. The composition is structured by local degree of abiotic stresses such as aridity and solar radiation and is also vertically stratified (Büdel, 2005, Grishkan et al., 2019, Hu et al., 2003).

The experiment`s bacterial and fungal community contained some cyanobacteria, lychens with cyanobacteria or algae as partner (cyano-lichen fungi, algal-lichen/chlorolychen fungi, respectively), and micro-fungal and bacterial heterotrophs that were reported to occur in soil biocrusts. **Names highlighted in bold were found in the dataset**. Algae (Green micro-algae and diatoms) and Bryophytes were not analyzed in this study.

In regards to Cyanobacteria, two taxa *Tychonema* and *Microcoleus* were, as mentioned before, significantly increased under NT management. These are rather free living (Strunecky et al., 2023, Zhang et al., 2016, Powell et al., 2015, Büdel, 2005). Two other Cyanobacteria, one of them *Nostoc*, were found in low relative abundance of the 16S marker gene (Figure A36). Although the taxa in this dataset were placed in Cyanobacteriales order, they were ascribed to the orders **Oscillatoriales**, or **Nostocales** (Büdel, 2005), which are also the orders in LPSN. Taxa from Chroococcales are missing in this experiment. *Nostoc* are widely reported to be able to fix nitrogen and enter symbiotic relationships with lychenious fungi. Reported, cyanobacterial taxa of biocrusts are (Büdel, 2005):

- Chroococcales (order in class Cyanophyceae acc. LPSN): *Aphanocapsa, Aphanothece, Chroococcidiopsis, Chroococcus, Coccochloris, Cyanothece, Gloeocapsa, Gloeothece, Katagnymene, Myxosarcina, Pleurocapsa*, *Rhabdogloea, Synechococcus, Xenotholos*
- Oscillatoriales (order in class Cyanophyceae acc. LPSN): *Crinalium, Heteroleibleinia, Leptolyngbya, Lyngbya, Komvophoron,* ***Microcoleus***, *Oscillatoria*, *Phormidium (**Phormidiaceae**), Plectonema, Porphyrosiphon, Pseudanabaena, Pseudophormidium, Schizothrix, Symploca,* ***Tychonema***
- Nostocales (order in class Cyanophyceae acc. LPSN): *Anabaena, Calothrix*, *Cylindrospermum, Microchaete, Nodularia,* ***Nostoc***, *Rivularia, Scytonema*, *Tolypothrix, Trichormus,* Stigonematales, *Chlorogloea, Chlorogloeopsis, Fischerella, Hapalosiphon, Mastigocladus, Stigonema*

Fungi that form associations with Cyanobacteria (Cyanolichens) make up 8-10% of lichens and are found mainly in the Ascomycota but also Basidiomycota (Lücking et al., 2009, Wedin et al., 2009). Cyanolichen fungi in the Ascomycota are found in the classes Lichinomycetes, **Eurotiomycetes**, and **Lecanoromycetes** and there in at least five orders of the Lichinales, **Chaetothyriales**, Agyriales, Peltigerales (Lücking et al., 2009). Cyano-lichen genera were reported to be *Collema, Gloeoheppia, Gonohymenia, Heppia, Leciophysma, Leptogium, Lichinella, Peccania, Peltigera, Peltula, Pseudopeltula, Psorotichia,* and *Synalissa* (Büdel, 2005). In this dataset, *Bradymyces*, *Exophiala* and several unassigned genera from the order

Chaetothyriales were found. Noteworthy, a fungus from the *Exophiala crusticola* was found with significant, though weak, increases under NT – cluster 2 (Phyl. Ascomycota, class Eurotiomycetes, order Chaetothyriales, fam. *Herpotrichiellaceae*). This species is, however, not cyano-lichen forming. It is characterized as dark-pigmented yeast-like fungus (dematiaceous) with heterotrophic use of sugars and small mass organics. However, it is speculated to be thriving on the use of exudates and osmolites from microbial primary producers (i.e. cyanobacteria) that derive from drying and wetting cycles. This type may therefore be described as a "secondary saprophyte" (Bates et al., 2006).

Algal-lichen fungi (Chlorolichen) genera are according to Büdel (2005): *Acarospora*, *Arthrorhaphis, Aspicilia*, *Bacidia, Baeomyces*, *Buellia*, *Caloplaca*, *Candelariella, Catapyrenium*, *Catolechia, Cetraria, Chondropsis, Cladia, Cladonia, Coelocaulon, Dacampia, Dactylina, Dermatocarpon, Desmazieria, Dibaeis*, *Diploschistes*, *Endocarpon*, *Eremastrella*, *Fulgensia*, *Fuscopannaria, Gyalecta, Gypsoplaca, Heterodea, Heteroplacidium, Involucropyrenium, Lecanora* (fam. **Lecanoraceae**, ord. **Lecanorales**), *Lecidea*, *Lecidella*, *Lecidoma, Leprocaulon, Leproloma, Leptochidium, Massalongia, Megaspora, Multiclavula, Mycobilimbia, Neofuscelia, Ochrolechia, Pannaria, Paraporpidia, Parmelia, Pertusaria, Phaeophyscia, Phaeorrhiza, Physconia, Placynthiella, Polyblastia, Protoblastenia, Psora*, *Psoroma,* ***Rinodina*** *(ord. Caliciales, fam. Physciaceae, OTU to Rinodina pyrina), Siphula, Solorina*, *Squamarina*, *Stereocaulon, Teloschistes, Texosporium, Toninia*, *Trapelia, Trapeliopsis, Xanthoparmelia, Xanthoria*

None of the potentially algal-lichen fungi in this dataset were significantly associated with tillage management (LinDA FDR>5%).

Heterotrophic micro-fungi were observed in soil crusts for the following taxa (Büdel, 2005, Grishkan et al., 2019):

- Zygomycetes***:*** ***Mortierella*** (Mortierellomycota), *M. humilis*, *Rhizopus spp.*, *Syncephalastrum*, ***Absidia*** *crymbifera*, *Actinomucor elegans*, ***Mucor hiemalis,*** *M. plumbeus, Rhizopus arrhizus*
- Ascomycetes/teleomorphic Ascomycota: ***Chaetomium*** *perlucidum, Graphyllium permundum,* ***Kalmusia*** *utahensis, Macroventuria wentii, Pleospora richtophensis, Aspergillus nidulans, A. rugulosus, Canariomyces notabilis*, *Chaetomidium strumarium*, Ch. succineum, Ch. Spp., *Sordaria fimicola*, *Sporomiella minima*
- Basidiomycetes: *Podaxis pistilaris, Disporotrichum dimorphosporum*
- Pale Hyphomycetes: ***Aspergillus*** *leporis, Aspergillus ustus, Aspergillus wentii,* ***Chrysosporium*** *pannorum, Fusarium flocciferum,* ***Fusarium*** *spp., Penicillium oxalicum,* ***Penicillium*** *spp., Pseudozyma sp.,* ***Stemphylium*** *iliois*, (Yeast II),
- Coelomycetes and dark Hyphomycetes/anamorphic Ascomycota: ***Alternaria*** *tenuis, Alternaria tenuissima, Bipolaris ravenelii, Cladosporium herbarum, Cladosporium macrocarpum, Embellisia clamydospora, Embellisia telluster, Epicoccum purpurascens, Phoma anserine, Phoma fimeti, Phoma leveillei, Phoma nebulosi, Phoma sp., Sphaeronaema spp.,* ***Trichoderma*** *sp., Ulocladium chartarum, Ulocladium multiforme,* ***Acremonium*** *charticola, Alternaria alternata, A. atra., A. phragmospora, A. raphani, Amerosporium concinnum, Aphanocladium album,* ***Aspergillus*** *spp., Aureobasidium pullulans,* ***Beauveria*** *bassiana, Boeremia exigua, Camarosporium aequivocum, Cladosporium cladosporioides, Coleophoma empetri,* ***Curvularia***

*inaequales, Engyodontium album, Epicoccum nigrum, Fusarium gibbosum, F. oxysporum,* ***F. solani****,* ***Lecanicillium*** *psaliotae, Malbranchea pulchella,* ***Metarhizium marquandii****, Neoscytalidium dimidiatum, Papulaspora pannosa, Paraboeremia putaminum,* ***Penicillium*** *aurantiogriseum, P. brevicompactum, P. glabrum, P. griseoroseum, P. corylophylum, P. herquei, P. janczewskii, P. lividum, P. simplicissimum, P. waksmani, Phoma medicaginis, Phomatodes nebulosi, Pithomyces africanus, Plenodomus tracheiphilus, Pleospora tarda, Pseudogymnoascus pannorum, Pyrenochaeta cava,* ***Schizothecium*** *inaequale, Setophoma terrestris,* ***Stachybotrys*** *chartarum,* ***Talaromyces*** *purpurogenus, T. variabilis,* ***Trichoderma*** *koningii, T. viride, Xepicula leucotricha*

- (Unidentified: Non-sporulating: Mycelia sterilia (light colored), Mycelia sterilia (dark colored))

Of the above observed taxa in the dataset (in bold), two taxa from the *Achaetomium* (family *Chaetomiaceae*, same family as *Chaetomium*), a *Fusarium,* and *Aspergillus thesauricus* were associated with tillage management. These were in clusters with significant relative increases under MT management.

Although the biocrust prokaryotic community is mainly formed by phototrophs such as Cyanobacteria, the heterotrophic bacterial component, often mycelial, also adds to biocrust formation, particle binding and protection, for example by production of mucosum or other exopolysaccharides, or by producing pigments as protection against UV radiation (Powell et al., 2015, Abed et al., 2010, Gundlapally and Garcia-Pichel, 2006, Büdel, 2005).

The second largest group of prokaryotes after Cyanobacteria was constituted by Proteobacteria from the Alpha and Delta group in Abed et al., (2010), whereas Beta- and Gammaproteobacteria were missing (this was also the observation in our dataset). Bacteroidetes were also abundantly present. Actinobacteria were absent in Abed et al., (2010) but most abundant in Gundlapally and Garcia-Pichel (2006).

The Alphaproteobacteria observed in biocrust were (Abed et al., 2010, Gundlapally and Garcia-Pichel, 2006):

- *Deinococcus* (UV resistant), *Belnapia moabensis* (crust bacterium), *Rhodospirillales, Methylobacterium, Rhodovarius,* ***Rubellimicrobium****, Azospirillum,* ***Paracraurococcus,***
- *Paracoccus,* ***Sphingomonas*** (two OTUs with sign. decrease in depth b NT vs ST)
- ***Microvirga*** *subterraneus* (one taxon significantly increased under NT vs ST both depths), ***Bosea*** *thiooxidans,* ***Roseomonas***, *Chelatococcus asaccharovorans*,

The Betaproteobacteria observed were:

- ***Massilia****,* ***Nitrosospira****,* ***Ramlibacter*** *tataouinensis*
- *Rubrivivax gelatinosus, Ralstonia solanacearum,* ***Acidovorax*** *facilis, Duganella violaceusniger,*

The Deltaproteobacteria observed were:

- *Cystobacter, Corallococcus, Hyalangium, Spirobacillus, Polyangium* (Predatory bacteria, ***Haliangium*** in dataset)*, Stigmatella*

The Gammaproteobacteria isolated were:

- ***Pseudomonas** luteola*

The Bacteroidetes observed were:

- *Rhodocytophaga*, ***Adhaeribacter*** (sign. decrease in NT vs ST depth a), and unidentified
- *Flexibacter flexilis*, Flavobacterial symbiont of Acromyrmex (**Flavobacterium** in dataset), *Spirosoma escalantus (fam.* ***Spirosomaceae*** in dataset*)*
- **Flavobacterium**, *Hymenobacter roseosalivarius* (several OTUs from fam. ***Hymenobacteraceae***), *Bifissio spartinacae*, *Taxeobacter,* ***Dyadobacter** fermentans*

The Actinobacteria observed (molecular and culturing) were:

- ***Streptomyces*** (two OTUs with sign. decrease under NT vs ST depth a/b)*, Sphaerobacter, Actinomadura,* ***Rubrobacter****,* ***Nonomuraea***
- *Arthrobacter spp., Modestobacter multiseptatus,* ***Conexibacter** woesei, Microbacterium spp.,* ***Actinoplanes** nipponensis* (in dataset sign. increase under NT vs ST both depths)*,* ***Pseudonocardia** (several also from fam.* ***Pseudonocardiaceae**),* ***Nocardioides** fulvus,* ***Agromyces****,* ***Williamsia** murale,* ***Geodermatophilus****, Micrococcus,* ***Blastococcus****, Friedmanniella,*

Other:

- *Thermomicrobium roseum, Alicychlobacillus pomorum*, ***Bacillus** licheniformis* (one taxon with sign. decrease under NT vs ST depth a), *Sulfobacillus disulfidooxidans*
- *Paenibacillus, Brevibacillus, Enterococcus*

Of the above listed heterotrophic bacteria that were also reported in biocrusts two taxa were associated with increases under NT management. This was most notably due to its relative abundance a rhizobium taxon from *Microvirga* (ord. Rhizobiales, fam. *Beijerinckiaceae*). The other taxon was *Actinoplanes* (Actinobacteria, ord. Micromonosporales).

Several of the above listed taxa were associated with significant decreases under NT management. These were two *Sphingomonas* (Alphaproteobacteria), *Adhaeribacter* (Bacteroidota, ord. Cytophagales), two *Streptomyces* (Actinobacteria), and a *Bacillus* (Firmicutes).

A scheme summarizing the above findings is hypothesized below. The decrease in disturbance with NT management could explain also the finding of decrease of qPCR prokaryotic:eukaryotic ratio (Table 2). Also, the strong increase of *Coprinellus curtus* (hetertrophic Basidiomycete) under NT could link to increase of biocrust organic matter built-up with conditions able to support crust heterotrophs.

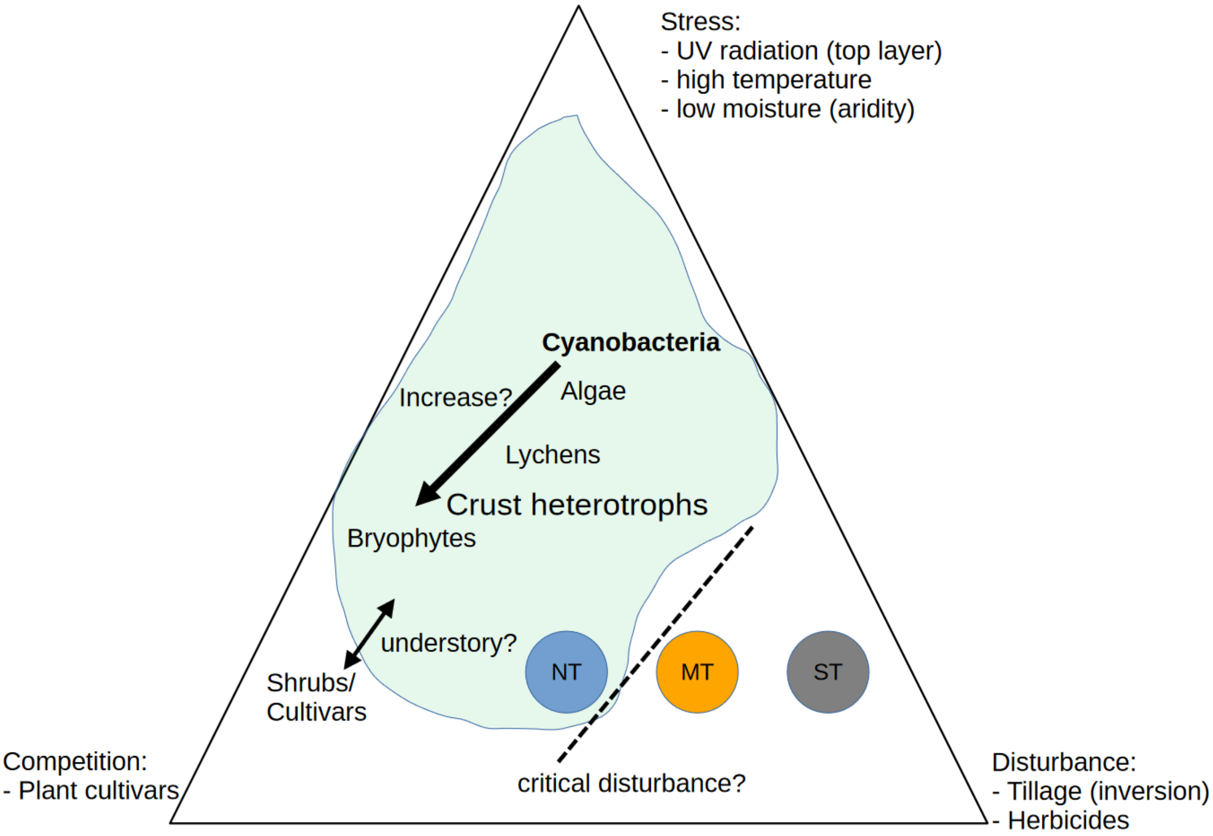


**Figure A64: CSR triangle after Grimes applied to biocrust and tillage management.**

#### 1.6 Well known bacterial plant pathogens

Well-known bacterial plant pathogens (Mansfield et al., 2012) are listed below. However, the short read 16S community metabarcoding approach is restricted from inferring on bacterial species and strains. Detected genera are highlighted in bold.

***Pseudomonas*** *syringae* pathovars: At least 29 pathovars each attacking a different host species

***Pseudomonas*** *savastanoi*

(n.s. with tillage treatments)

*Ralstonia solanacearum*

*Agrobacterium tumefaciens*

*Xanthomonas oryzae* pv. Oryzae

*Xanthomonas campestris* pathovars

*Xanthomonas axonopodis* pathovars

(hit to ***Pseudoxanthomonas***, n.s. with tillage treatments)

*Erwinia amylovora*

*Xylella fastidiosa*

*Dickeya* (*dadantii* and *solani*)

*Pectobacterium carotovorum* and *Pectobacterium atrosepticum*

*Clavibacter michiganensis* (*michiganensis* and *sepedonicus*),

Candidatus Liberibacter asiaticus

#### 2. Supplementary tables

##### 2.1 Bray-Curtis dissimilarities of soil biota

**Table A1** Treatment effect testing of tillage and depth (Nematodes only bulked depth 0-20 cm) on Bray-Curtis dissimilarities of community composition with permutational multivariate ANOVA (perMANOVA). Multivariate group dispersions were tested for homogeneity in betadisper test and were not found to be significant at p>0.05. Block was not found to be significant for acari and nematodes. Block was used to restrict permutation for bacteria and fungi. Model term significance: $p \leq 0.05$ *, 0.01 **, 0.001 ***.

| | Tillage | | Depth | | Tillage:Depth | |
|---|---|---|---|---|---|---|
| Bacteria | $F_{2,18}$=4.8 *** | $R^2$=32% | $F_{1,18}$=1.2 | $R^2$=4% | $F_{2,18}$=0.5 | $R^2$=4% |
| Fungi | $F_{2,18}$=3.8 *** | $R^2$=27% | $F_{1,18}$=1.0 | $R^2$=4% | $F_{2,18}$=0.7 | $R^2$=5% |
| Acari | $F_{2,18}$=5.3 *** | $R^2$=34% | $F_{1,18}$=0.9 | $R^2$=3% | $F_{2,18}$=0.6 | $R^2$=4% |
| Nematodes | $F_{2,9}$=1.1 | $R^2$=20% | | | | |

Dal Cortivo, C., Ferrari, M., Visioli, G., Lauro, M., Fornasier, F., Barion, G., Panozzo, A., Vamerali, T., 2020. Effects of seed-applied biofertilizers on rhizosphere biodiversity and growth of common wheat (Triticum aestivum L.) in the field. *Frontiers in Plant Science*, 11, 72.

da Silva, G. A., Corazon-Guivin, M. A., de Assis, D. M. A., Oehl, F., 2023. Blaszkowskia, a new genus in Glomeraceae. *Mycological Progress*, 22(11), 74.

Dedysh, S. N., Yilmaz, P., 2018. Refining the taxonomic structure of the phylum Acidobacteria. *International Journal of Systematic and Evolutionary Microbiology*, 68(12), 3796-3806.

Dedysh, S. N., Kulichevskaya, I. S., Beletsky, A. V., Ivanova, A. A., Rijpstra, W. I. C., Damsté, J. S. S., Mardanov, A.V., Ravin, N. V., 2020. Lacipirellula parvula gen. nov., sp. nov., representing a lineage of planctomycetes widespread in low-oxygen habitats, description of the family Lacipirellulaceae fam. nov. and proposal of the orders Pirellulales ord. nov., Gemmatales ord. nov. and Isosphaerales ord. nov. *Systematic and Applied Microbiology*, 43(1), 126050.

Doveri, F., 2014. An update on the genera Ascobolus and Saccobolus with keys and descriptions of three coprophilous species, new to Italy. *Mycosphere*, 5(1), 86-135.

Ekanayaka, A. H., Hyde, K. D., Gentekaki, E., McKenzie, E. H. C., Zhao, Q., Bulgakov, T. S., Camporesi, E., 2019. Preliminary classification of Leotiomycetes. *Mycosphere* 10(1): 310–489.

Franzmann, P. D., Skerman, V. B. D., 1984. Gemmata obscuriglobus, a new genus and species of the budding bacteria. *Antonie Van Leeuwenhoek*, 50, 261-268.

Fukunaga, Y., Kurahashi, M., Sakiyama, Y., Ohuchi, M., Yokota, A., Harayama, S., 2009. Phycisphaera mikurensis gen. nov., sp. nov., isolated from a marine alga, and proposal of Phycisphaeraceae fam. nov., Phycisphaerales ord. nov. and Phycisphaerae classis nov. in the phylum Planctomycetes. *The Journal of General and Applied Microbiology*, 55(4), 267-275.

Guevara-Suarez, M., García, D., Cano-Lira, J. F., Guarro, J., Gené, J., 2020. Species diversity in Penicillium and Talaromyces from herbivore dung, and the proposal of two new genera of penicillium-like fungi in Aspergillaceae. *Fungal Systematics and Evolution*, 5(1), 39-76.

Gundlapally, S. R., Garcia-Pichel, F., 2006. The community and phylogenetic diversity of biological soil crusts in the Colorado Plateau studied by molecular fingerprinting and intensive cultivation. *Microbial Ecology*, 52, 345-357.

Gupta, R. S., Chander, P., George, S., 2013. Phylogenetic framework and molecular signatures for the class Chloroflexi and its different clades; proposal for division of the class Chloroflexi class. nov. into the suborder Chloroflexineae subord. nov., consisting of the emended family Oscillochloridaceae and the family Chloroflexaceae fam. nov., and the suborder Roseiflexineae subord. nov., containing the family Roseiflexaceae fam. nov. *Antonie van Leeuwenhoek*, 103, 99-119.

Gupta, R. S., Patel, S., Saini, N., Chen, S., 2020. Robust demarcation of 17 distinct Bacillus species clades, proposed as novel Bacillaceae genera, by phylogenomics and comparative genomic analyses: description of Robertmurraya kyonggiensis sp. nov. and proposal for an emended genus Bacillus limiting it only to the members of the Subtilis and Cereus clades of species. *International Journal of Systematic and Evolutionary Microbiology*, 70(11), 5753-5798.

Grishkan, I., Lázaro, R., Kidron, G. J., 2019. Cultured microfungal communities in biological soil crusts and bare soils at the Tabernas Desert, Spain. *Soil systems*, 3(2), 36.

Hansen, K., Pfister, D.H., 2006. Systematics of the Pezizomycetes—the operculate discomycetes. *Mycologia*, 98(6), 1029-1040.

Hansen, K., Perry, B. A., Dranginis, A. W., Pfister, D.H., 2013. A phylogeny of the highly diverse cup-fungus family Pyronemataceae (Pezizomycetes, Ascomycota) clarifies relationships and evolution of selected life history traits. *Molecular Phylogenetics and Evolution*, 67(2), 311-335.

Harman, G. E., 2006. Overview of Mechanisms and Uses of Trichoderma spp. *Phytopathology*, 96(2), 190-194.

Helaly, S. E., Thongbai, B., Stadler, M., 2018. Diversity of biologically active secondary metabolites from endophytic and saprotrophic fungi of the ascomycete order Xylariales. *Natural Product Reports*, 35(9), 992-1014.

Huber, K. J., & Overmann, J., 2018. Vicinamibacteraceae fam. nov., the first described family within the subdivision 6 Acidobacteria. *International Journal of Systematic and Evolutionary Microbiology*, 68(7), 2331-2334.

Illescas, M., Rubio, M. B., Hernandez-Ruiz, V., Moran-Diez, M. E., Martinez de Alba, A. E., Nicolás, C., Monte, E., Hermosa, R., 2020. Effect of inorganic N top dressing and Trichoderma harzianum seed-inoculation on crop yield and the shaping of root microbial communities of wheat plants cultivated under high basal N fertilization. *Frontiers in Plant Science*, 11, 575861.

Index Fungorum. Accessed March 2023. Available under: http://www.indexfungorum.org/

Ivanova, A. A., Kulichevskaya, I. S., Dedysh, S. N., 2021. Gemmata palustris sp. nov., a novel Planctomycete from a fen in northwestern Russia. *Microbiology*, 90, 598-606.

Jayasiri, S. C., Hyde, K. D., Jones, E. B. G., McKenzie, E. H. C., Jeewon, R., Phillips, A. J. L., Bhat, D.J., Wanasinghe, D.N., Liu, J.K., Lu, Y.Z., Kang, J.C., Xu, J., Karunarathna, S. C., 2019. Diversity, morphology and molecular phylogeny of Dothideomycetes on decaying wild seed pods and fruits. *Mycosphere*, 10, 1-186.

Jung, B., Park, S. Y., Lee, Y. W., Lee, J., 2013. Biological efficacy of Streptomyces sp. strain BN1 against the cereal head blight pathogen Fusarium graminearum. *The Plant Pathology Journal*, 29(1), 52.

Kant, R., van Passel, M. W., Sangwan, P., Palva, A., Lucas, S., Copeland, A., Lapidus, A., del Rio, T. G., Dalin, E., Tice, H., Bruce, D., Goodwin, L., Pitluck, S., Chertkov, O., Larimer, F. W., Land, M. L., Hauser, L., Brettin, T. S., Detter, J. C., Han, S., de Vos, W. M., Janssen, P. H., Smidt, H., 2011. Genome sequence of" Pedosphaera parvula" Ellin514, an aerobic Verrucomicrobial isolate from pasture soil. *Journal of Bacteriology*, 193(11), 2900-2901.

Kepler, R. M., Sung, G. H., Ban, S., Nakagiri, A., Chen, M. J., Huang, B., Li, Z., Spatafora, J. W., 2012. New teleomorph combinations in the entomopathogenic genus Metacordyceps. *Mycologia*, 104(1), 182-197.

Khianngam, S., Tanasupawat, S., Akaracharanya, A., Kim, K. K., Lee, K. C., Lee, J. S., 2010. Cohnella thailandensis sp. nov., a xylanolytic bacterium from Thai soil. *International Journal of Systematic and Evolutionary Microbiology*, 60(10), 2284-2287.

Kulichevskaya, I. S., Suzina, N. E., Liesack, W., Dedysh, S. N., 2010. Bryobacter aggregatus gen. nov., sp. nov., a peat-inhabiting, aerobic chemo-organotroph from subdivision 3 of the Acidobacteria. *International Journal of Systematic and Evolutionary Microbiology*, 60(2), 301-306.

Kulichevskaya, I. S., Ivanova, A. A., Suzina, N. E., Rijpstra, W. I. C., Sinninghe Damsté, J. S., Dedysh, S. N., 2016. Paludisphaera borealis gen. nov., sp. nov., a hydrolytic planctomycete from northern wetlands, and proposal of Isosphaeraceae fam. nov. International Journal of Systematic and Evolutionary Microbiology, 66(2), 837-844.

Kulichevskaya, I. S., Ivanova, A. A., Baulina, O. I., Rijpstra, W. I. C., Sinninghe Damsté, J. S., Dedysh, S. N., 2017. Fimbriiglobus ruber gen. nov., sp. nov., a Gemmata-like planctomycete from Sphagnum peat bog and the proposal of Gemmataceae fam. nov. *International Journal of Systematic and Evolutionary Microbiology*, 67(2), 218-224.

Liu, Q., Liu, H. C., Zhou, Y. G., Xin, Y. H., 2019. Stenotrophobium rhamnosiphilum gen. nov., sp. nov., isolated from a glacier, proposal of Steroidobacteraceae fam. nov. in Nevskiales and emended description of the family Nevskiaceae. *International Journal of Systematic and Evolutionary Microbiology*, 69(5), 1404-1410.

Liu, X. Z., Wang, Q. M., Göker, M., Groenewald, M., Kachalkin, A. V., Lumbsch, H. T., Millanes, A.M., Wedin, M., Yurkov, A.M., Boekhout, T., Bai, F. Y., 2015. Towards an integrated phylogenetic classification of the Tremellomycetes. *Studies in Mycology*, 81(1), 85-147.

Liu, Y. J., Liu, Y., Zhang, K. Q., 2008. Xanthothone, a new nematicidal N-compound from Coprinus xanthothrix. *Chemistry of Natural Compounds*, 44, 203-205.

Liu, Y. H., Bau, T., 2023. Biological characteristics and ontogeny of Coprinellus xanthothrix. CABI. Available under: https://www.cabidigitallibrary.org/doi/pdf/10.5555/20230150617.

Lücking, R., Lawrey, J. D., Sikaroodi, M., Gillevet, P. M., Chaves, J. L., Sipman, H. J., Bungartz, F., 2009. Do lichens domesticate photobionts like farmers domesticate crops? Evidence from a previously unrecognized lineage of filamentous cyanobacteria. *American Journal of Botany*, 96(8), 1409-1418.

Lombard, L., Van der Merwe, N. A., Groenewald, J. Z., Crous, P. W., 2015. Generic concepts in Nectriaceae. *Studies in Mycology*, 80(1), 189-245.

Madigan, M. T., Martinko, J. M., Bender, K. S., Buckley, D. H., 2021. *Brock Biology of Microorganisms*. 16th Edition. Pearson.

Mansfield, J., Genin, S., Magori, S., Citovsky, V., Sriariyanum, M., Ronald, P., Dow, M., Verdier, V., Beer, S. V., Machado, M. A., Toth, I., Salmand, G., Foster, G. D., 2012. Top 10 plant pathogenic bacteria in molecular plant pathology. *Molecular Plant Pathology*, 13(6), 614-629.

Marin-Felix, Y., Miller, A. N., 2022. Corrections to recent changes in the taxonomy of the Sordariales. *Mycological Progress*, 21(8), 69.

Matsumoto, A., Kasai, H., Matsuo, Y., Ōmura, S., Shizuri, Y., Takahashi, Y., 2009. Ilumatobacter fluminis gen. nov., sp. nov., a novel actinobacterium isolated from the sediment of an estuary. *The Journal of General and Applied Microbiology*, 55(3), 201-205.

Matsumoto, A., Kasai, H., Matsuo, Y., Shizuri, Y., Ichikawa, N., Fujita, N., Omura, S., Takahashi, Y., 2013. Ilumatobacter nonamiense sp. nov. and Ilumatobacter coccineum sp. nov., isolated from seashore sand. *International Journal of Systematic and Evolutionary Microbiology*, 63(Pt_9), 3404-3408.

Nakasaki, K., Saito, M., Suzuki, N., 2007. Coprinellus curtus (Hitoyo-take) prevents diseases of vegetables caused by pathogenic fungi. *FEMS Microbiology Letters*, 275(2), 286-291.

Naushad, S., Adeolu, M., Wong, S., Sohail, M., Schellhorn, H. E., Gupta, R. S., 2015. A phylogenomic and molecular marker based taxonomic framework for the order Xanthomonadales: proposal to transfer the families Algiphilaceae and Solimonadaceae to the order Nevskiales ord. nov. and to create a new family within the order Xanthomonadales, the family Rhodanobacteraceae fam. nov., containing the genus Rhodanobacter and its closest relatives. *Antonie van Leeuwenhoek*, 107, 467-485.

Nguyen, T. T., Park, S. W., Pangging, M., Lee, H. B., 2019. Molecular and morphological confirmation of three undescribed species of Mortierella from Korea. *Mycobiology*, 47(1), 31-39.

Nouioui, I., Carro, L., García-López, M., Meier-Kolthoff, J. P., Woyke, T., Kyrpides, N. C., Pukal, R., Klenk, H.-P., Goodfellow, M., Göker, M., 2018. Genome-Based Taxonomic Classification of the Phylum Actinobacteria. *Frontiers in Microbiology*, 9, 2007.

Novakova, A., Hubka, V., Saiz-Jimenez, C., Kolarik, M., 2012. Aspergillus baeticus sp. nov. and Aspergillus thesauricus sp. nov., two species in section Usti from Spanish caves. *International Journal of Systematic and Evolutionary Microbiology*, 62(Pt_11), 2778-2785.

Olariaga, I., Vooren, N. V., Carbone, M., Hansen, K., 2015. A monograph of Otidea (Pyronemataceae, Pezizomycetes). *Persoonia-Molecular Phylogeny and Evolution of Fungi*, 35(1), 166-229.

Padamsee, M., Matheny, P. B., Dentinger, B. T., McLaughlin, D. J., 2008. The mushroom family Psathyrellaceae: evidence for large-scale polyphyly of the genus Psathyrella. *Molecular Phylogenetics and Evolution*, 46(2), 415-429.

Pascual, J., Wüst, P. K., Geppert, A., Foesel, B. U., Huber, K. J., Overmann, J., 2015. Novel isolates double the number of chemotrophic species and allow the first description of higher taxa in Acidobacteria subdivision 4. *Systematic and Applied Microbiology*, 38(8), 534-544.

Parte, A.C., Sardà Carbasse, J., Meier-Kolthoff, J.P., Reimer, L.C. Göker, M., 2020. List of Prokaryotic names with Standing in Nomenclature (LPSN) moves to the DSMZ. International Journal of Systematic and Evolutionary Microbiology, 70, 5607-5612; DOI: 10.1099/ijsem.0.004332

Peiris, D., Dunn, W. B., Brown, M., Kell, D. B., Roy, I., Hedger, J. N., 2008. Metabolite profiles of interacting mycelial fronts differ for pairings of the wood decay basidiomycete fungus, Stereum hirsutum with its competitors Coprinus micaceus and Coprinus disseminatus. *Metabolomics*, 4, 52-62.

Pem, D., Jeewon, R., Chethana, K. W. T., Hongsanan, S., Doilom, M., Suwannarach, N., Hyde, K. D., 2021. Species concepts of Dothideomycetes: classification, phylogenetic inconsistencies and taxonomic standardization. *Fungal Diversity*, 109(1), 283-319.

Perry, B. A., Hansen, K., Pfister, D. H., 2007. A phylogenetic overview of the family Pyronemataceae (Ascomycota, Pezizales). *Mycological Research*, 111(5), 549-571.

Phookamsak, R., Liu, J. K., McKenzie, E. H., Manamgoda, D. S., Ariyawansa, H., Thambugala, K. M., Dai, D.Q.,Camporesi, E., Chukeatirote, E., Wijayawardene, N.N., Bahkali, A.H., Mortimer, P.E., Xu, J.C., Hyde, K.D., 2014. Revision of Phaeosphaeriaceae*. Fungal Diversity*, 68(1), 159-238.

Powell, J. T., Chatziefthimiou, A. D., Banack, S. A., Cox, P. A., Metcalf, J. S., 2015. Desert crust microorganisms, their environment, and human health. *Journal of Arid Environments*, 112, 127-133.

Powell, M. J., Letcher, P. M., Chambers, J. G., Roychoudhury, S., 2015. A new genus and family for the misclassified chytrid, Rhizophlyctis harderi. *Mycologia*, 107(2), 419-431.

Quaedvlieg, W., Kema, G. H. J., Groenewald, J. Z., Verkley, G. J. M., Seifbarghi, S., Razavi, M., Mirzadi Gohari, A., Mehrabi, R., Crous, P. W. (2011). Zymoseptoria gen. nov.: a new genus to accommodate Septoria-like species occurring on graminicolous hosts. *Persoonia-Molecular Phylogeny and Evolution of Fungi*, 26(1), 57-69.

Redhead, S. A., Vilgalys, R., Moncalvo, J. M., Johnson, J., Hopple Jr, J. S., 2001. Coprinus Pers. and the disposition of Coprinus species sensu lato. *Taxon*, 50(1), 203-241.

Russ, C., Lang, B. F., Chen, Z., Gujja, S., Shea, T., Zeng, Q., Young, S., Cuomo, C. A., Nusbaum, C., 2016. Genome sequence of Spizellomyces punctatus. *Genome Announcements*, 4(4), 10-1128.

Salgado-Salazar, C., Beirn, L. A., Ismaiel, A., Boehm, M. J., Carbone, I., Putman, A. I., Tredway, L.P., Clarke, B.B., Crouch, J. A., 2018. Clarireedia: A new fungal genus comprising four pathogenic species responsible for dollar spot disease of turfgrass. *Fungal Biology*, 122(8), 761-773.

Sangwan, P., Chen, X., Hugenholtz, P., Janssen, P. H., 2004. Chthoniobacter flavus gen. nov., sp. nov., the first pure-culture representative of subdivision two, Spartobacteria classis nov., of the phylum Verrucomicrobia. *Applied and Environmental Microbiology*, 70(10), 5875-5881.

Sangwan, P., Kovac, S., Davis, K. E., Sait, M., Janssen, P. H., 2005. Detection and cultivation of soil Verrucomicrobia. *Applied and Environmental Microbiology*, 71(12), 8402-8410.

Sieverding, E., da Silva, G. A., Berndt, R., Oehl, F., 2015. Rhizoglomus, a new genus of the Glomeraceae. *Mycotaxon*, 129(2), 373-386.

Simmons, D. R., 2011. Phylogeny of Powellomycetaceae fam. nov. and description of Geranomyces variabilis gen. et comb. nov. *Mycologia*, 103(6), 1411-1420.

Singh, H., Du, J., Won, K., Yang, J. E., Yin, C., Kook, M., Yi, T. H., 2015. Massilia arvi sp. nov., isolated from fallow-land soil previously cultivated with Brassica oleracea, and emended description of the genus Massilia. *International Journal of Systematic and Evolutionary Microbiology*, 65(Pt_10), 3690-3696.

Strunecký, O., Ivanova, A. P., Mareš, J., 2023. An updated classification of cyanobacterial orders and families based on phylogenomic and polyphasic analysis. *Journal of Phycology*, 59(1), 12-51.

Taylor, D. L., Sinsabaugh, R. L., 2015. The soil fungi: occurrence, phylogeny, and ecology. In: *Soil Microbiology, Ecology, and Biochemistry*. Chapter 4, pp. 77-109. Ed. Paul, E.A.

Tedersoo, L., Sánchez-Ramírez, S., Koljalg, U., Bahram, M., Döring, M., Schigel, D., May, T., Ryberg, M., Abarenkov, K., 2018. High-level classification of the Fungi and a tool for evolutionary ecological analyses. *Fungal Diversity*, 90, 135-159.

Yamada, T., Sekiguchi, Y., Hanada, S., Imachi, H., Ohashi, A., Harada, H., Kamagata, Y., 2006. Anaerolinea thermolimosa sp. nov., Levilinea saccharolytica gen. nov., sp. nov. and Leptolinea tardivitalis gen. nov., sp. nov., novel filamentous anaerobes, and description of the new classes Anaerolineae classis nov. and Caldilineae classis nov. in the bacterial phylum Chloroflexi. *International Journal of Systematic and Evolutionary Microbiology*, 56(6), 1331-1340.

Yoon, J., Matsuo, Y., Adachi, K., Nozawa, M., Matsuda, S., Kasai, H., Yokota, A., 2008. Description of Persicirhabdus sediminis gen. nov., sp. nov., Roseibacillus ishigakijimensis gen. nov., sp. nov., Roseibacillus ponti sp. nov., Roseibacillus persicicus sp. nov., Luteolibacter pohnpeiensis gen. nov., sp. nov. and Luteolibacter algae sp. nov., six marine members of the phylum 'Verrucomicrobia', and emended descriptions of the class Verrucomicrobiae, the order Verrucomicrobiales and the family Verrucomicrobiaceae. *International Journal of Systematic and Evolutionary Microbiology*, 58(4), 998-1007.

Valenzuela-Lopez, N., Cano-Lira, J. F., Guarro, J., Sutton, D. A., Wiederhold, N., Crous, P. W., Stchigel, A. M., 2018. Coelomycetous Dothideomycetes with emphasis on the families Cucurbitariaceae and Didymellaceae. *Studies in Mycology*, 90(1), 1-69.

van Brummelen, J., 1972. Ascocarp ontogeny and a natural classification of the Ascobolaceae. *Persoonia-Molecular Phylogeny and Evolution of Fungi*, 6(4), 389-394.

Visioli, G., Lauro, M., Vamerali, T., Dal Cortivo, C., Panozzo, A., Folloni, S., Piazza, C., Ranieri, R., 2020. A comparative study of organic and conventional management on the rhizosphere microbiome, growth and grain quality traits of tritordeum. *Agronomy*, 10(11), 1717.

Wachowska, U., Irzykowski, W., Jędryczka, M., Stasiulewicz-Paluch, A. D., Głowacka, K., 2013. Biological control of winter wheat pathogens with the use of antagonistic Sphingomonas bacteria under greenhouse conditions. *Biocontrol Science and Technology*, 23(10), 1110-1122.

Waite, D. W., Chuvochina, M., Pelikan, C., Parks, D. H., Yilmaz, P., Wagner, M., Loy, A., Naganuma, T., Nakai, R., Whitman, W. B., Hahn, M.W., Kuever, J., Hugenholtz, P., 2020. Proposal to reclassify the proteobacterial classes Deltaproteobacteria and Oligoflexia, and the phylum Thermodesulfobacteria into four phyla reflecting major functional capabilities. *International Journal of Systematic and Evolutionary Microbiology*, 70(11), 5972-6016.

Walther, G., Wagner, L., Kurzai, O., 2019. Updates on the taxonomy of Mucorales with an emphasis on clinically important taxa. *Journal of Fungi*, 5(4), 106.

Wedin, M., Wiklund, E., Jørgensen, P. M., Ekman, S., 2009. Slippery when wet: phylogeny and character evolution in the gelatinous cyanobacterial lichens (Peltigerales, Ascomycetes). *Molecular Phylogenetics and Evolution*, 53(3), 862-871.

Wijayawardene, N. N., Pawłowska, J., Letcher, P. M., Kirk, P. M., Humber, R. A., Schüßler, A., Wrzosek, M., Muszewska, A., Okrasinska, A., Istel, L., Gesiorska, A., Mungai, P., Lateef, A.A., Rajeshkumar, K.C., Singh, R.V., Radek, R., Walther, G., Wagner, L., Walker, C., Wijesundara, D.S.A., Papizadeh, M., Dolatabadi1, S., Shenoy, B.D., Tokarev, Y.S., Lumyong, S., Hyde, K. D.,, 2018. Notes for genera: Basal clades of fungi (Including aphelidiomycota, basidiobolomycota, blastocladiomycota, calcarisporiellomycota, caulochytriomycota, chytridiomycota, entomophthoromycota, glomeromycota, kickxellomycota, monoblepharomycota, mortierellomycota, mucoromycota, neocallimastigomycota, olpidiomycota, rozellomycota and zoopagomycota). *Fungal Diversity*, 92, 43-129.

Wilhelm, R. C., van Es, H. M., Buckley, D. H., 2022. Predicting measures of soil health using the microbiome and supervised machine learning. *Soil Biology and Biochemistry*, 164, 108472.

Willems, A., De Ley, J., Gillis, M., Kersters, K., 1991. Comamonadaceae, a new family encompassing the acidovorans rRNA complex, including Variovorax paradoxus gen. nov., comb. nov., for Alcaligenes paradoxus (Davis 1969). *International Journal of Systematic and Evolutionary Microbiology*, 41(3), 445-450.

Wüst, P. K., Foesel, B. U., Geppert, A., Huber, K. J., Luckner, M., Wanner, G., Overmann, J., 2016. Brevitalea aridisoli, B. deliciosa and Arenimicrobium luteum, three novel species of Acidobacteria subdivision 4 (class Blastocatellia) isolated from savanna soil and description of the novel family Pyrinomonadaceae. *International Journal of Systematic and Evolutionary Microbiology*, 66(9), 3355-3366.

Zhang, F., Yang, Y. Q., Zhou, F. P., Xiao, W., Boonmee, S., Yang, X. Y., 2023. Morphological and Phylogenetic Characterization of Five Novel Nematode-Trapping Fungi (Orbiliomycetes) from Yunnan, China. *Journal of Fungi*, 9(7), 735.

Zhang, H., Song, G., Shao, J., Xiang, X., Li, Q., Chen, Y., Yang, P., Yu, G., 2016. Dynamics and polyphasic characterization of odor-producing cyanobacterium Tychonema bourrellyi from Lake Erhai, China. Environmental Science and Pollution Research, 23, 5420-5430.

Zhang, J., Song, F., Xin, Y. H., Zhang, J., Fang, C., 2009. Microvirga guangxiensis sp. nov., a novel alphaproteobacterium from soil, and emended description of the genus Microvirga. *International Journal of Systematic and Evolutionary Microbiology*, *59*(8), 1997-2001.

Zhang, X. J., Zhang, J., Yao, Q., Feng, G. D., Zhu, H. H., 2019. Microvirga flavescens sp. nov., a novel bacterium isolated from forest soil and emended description of the genus Microvirga. *International Journal of Systematic and Evolutionary Microbiology*, *69*(3), 667-671.

Zhi, X. Y., Li, W. J., Stackebrandt, E., 2009. An update of the structure and 16S rRNA gene sequence-based definition of higher ranks of the class Actinobacteria, with the proposal of two new suborders and four new families and emended descriptions of the existing higher taxa. *International Journal of Systematic and Evolutionary Microbiology*, 59(3), 589-608.
